# Anthropogenic climate change will likely outpace coral range expansion

**DOI:** 10.1101/2024.07.23.604846

**Authors:** Noam S. Vogt-Vincent, James M. Pringle, Christopher E. Cornwall, Lisa C. McManus

## Abstract

Past coral range expansions suggest that high-latitude environments may serve as refugia, potentially buffering tropical biodiversity loss due to climate change. We explore this possibility for corals globally, using a dynamical metacommunity model incorporating temperature, light intensity, pH, and four distinct, interacting coral assemblages. This model reasonably reproduces the observed distribution and recent decline of corals across the Indo-Pacific and Caribbean. Our simulations suggest that there is a mismatch between the timescales of coral reef decline and range expansion under future predicted climate change. Whereas the most severe declines in coral cover will likely occur within 60–80 years, significant tropical coral range expansion requires centuries. The absence of large-scale coral refugia in the face of rapid anthropogenic climate change emphasises the urgent need to reduce greenhouse gas emissions, and mitigate non-thermal stressors for corals, both in the tropics and high-latitudes.

## Introduction

Coral reefs are immeasurably valuable marine environments, supporting a third of marine species and the livelihoods of millions of people, particularly in low- and middle-income countries (*1,2*). Unfortunately, reef-building corals are also highly sensitive to environmental change, with the significant observed decline in coral cover expected to continue under future climate projections (*3*).

As the ocean warms and isotherms migrate poleward, thermal niches shift to higher latitudes, and a rise in marine species diversity in subtropical and temperate seas is expected to accompany the decline in the tropics (*4*). This expected redistribution of marine life has led to the suggestion that high-latitude seas may act as climate change ‘refugia’ for reef-building corals (*5*), potentially serving as a long-term buffer against declining coral diversity. The scope for latitudinal range expansion for corals may be limited by light attenuation, acidification, competition with macroalgae, local anthropogenic stressors, and a lack of settlement cues (*6*). Nevertheless, many statistical species distribution models support predictions of coral range expansion (*7, 8*), and the geological record also provides evidence that coral reef distribution expands and contracts in response to geologically recent environmental change (*9–11*). Indeed, a rise in coral recruitment has already been observed at some subtropical field sites (*12*), and recent apparent poleward expansion has occurred in Australia, Florida, and Japan (*13–15*).

On the other hand, much of this recent range expansion may have been driven by the proliferation of coral species already present in high-latitude environments, rather than the introduction of new species from the tropics (*16, 17*), and these marginal communities are environmentally and functionally distinct from tropical coral reefs (*18, 19*). The recent rate of warming is also exceptional within the context of the late Pleistocene (*20*) and exceeds the rate that can be resolved from most geological records (*21*), making it challenging to contextualise the response of coral reefs to rapid palaeoenvironmental change. It is therefore unclear how analogous or useful past coral range expansions are in predicting coral reef dynamics over the coming century. Corals are also relatively slow-growing organisms and, in contrast to many other marine taxa, are wholly reliant on passive larval dispersal to expand their range. As a result, it is unrealistic to expect the distribution of corals to be in equilibrium with the rapidly changing physical environment, an assumption made by most species distribution models used to forecast coral range shifts.

The capacity for coral range expansion into high-latitude seas therefore remains uncertain, limiting our ability to predict the impact of climate change on coral reef ecosystems. Range expansion has important implications for global coral distribution, diversity and ecosystem functioning. Furthermore, predicting where and when refugia for tropical corals are likely to emerge may also permit proactive or dynamic management plans for environments that may not have previously been prioritized (*18*).

In this study, we expand on earlier work (*22*) to develop a simple process-based, ecoevolutionary metacommunity model, the Coral Eco-evo Range Expansion Simulation (CERES), which reasonably reproduces the recent distribution of coral reefs and decadal trends in coral cover. Using CERES, we demonstrate that there is a mismatch between the expected rate of future climate change and the rate of coral range expansion, indicating that range expansion alone is unlikely to buffer coral diversity against anthropogenic climate change.

## Results

### Reproducing observed coral biogeography and trends

CERES simulates the response of four interacting coral assemblages to environmental change using an eco-evolutionary metapopulation framework, assuming that populations grow through the linear extension of coral colonies and the establishment of new coral colonies through larval dispersal (figure 1, see also *Materials and Methods*). We consider competing *tropical* (high diversity, analogous to low-latitude coral reefs) and *subtropical* (low diversity, analogous to high-latitude non-reefal coral communities) assemblages, further divided into *fast-growing* and *slow-growing* life history strategies (*23*). We assume that habitat shallower than 20 m is ‘potentially habitable’ for corals, thereby simulating the size of each assemblage across 87,965 subpopulations at monthly temporal resolution.

**Figure 1:**
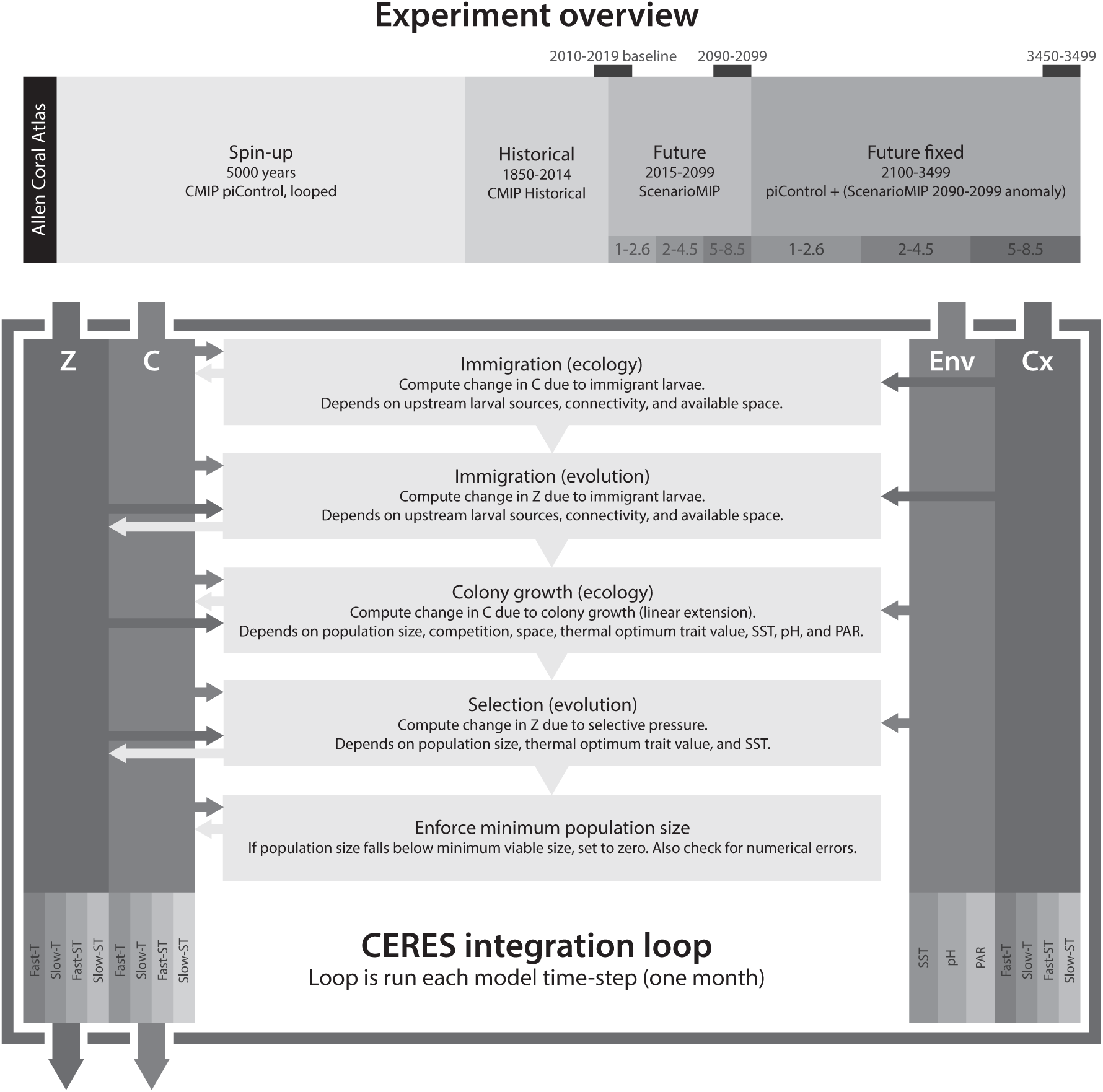
Experimental design used in this study (*top*) and schematic overview of the CERES integration loop (*bottom*). *Z* and *C* are the model dependent variables, the thermal optima and coral cover for each assemblage and subpopulation. ‘Env’ and ‘Cx’ represent the model independent variables, the environmental variables (SST, pH, and PAR), and potential connectivity. Inward and outward arrows respectively represent variables being read and updated by the model.

Colony linear extension rate is modelled as a function of the monthly sea surface temperature (SST), pH, photosynthetically available radiation (PAR) at the seafloor, and inter-assemblage competition; whereas population growth through the establishment of new colonies is based on estimates of fecundity, post-spawning processes such as mortality, and potential connectivity (*24*). CERES further simulates a quantitative trait, the mean thermal optimum, for each assemblage and subpopulation. This trait value evolves based on immigration and selection, the latter being proportional to the additive genetic variance and the gradient of the fitness landscape with respect to the trait value (*22*). The colony linear extension rate falls with distance from optimal environmental conditions, and assemblages experience mortality beyond low and high thermal stress thresholds (referenced to the thermal optimum) and under compound environmental stress. The fitness of each assemblage with respect to pH, PAR and competition does not evolve. We parameterise the model based on empirical physiological measurements where possible, which we translate into a population growth rate by assuming that the size structure of coral colonies is log-normal (*25*) and static. Unlike statistical models, CERES is fully process-based and predicts coral cover and thermal optima for each assemblage and subpopulation as emergent properties of the system.

For each of twelve CMIP6 models (*26*) with a transient climate response in the ‘very likely’ 1.2–2.4 °C range (*27*), we impose a 5000-year spin-up period under repeating preindustrial forcing, followed by historical forcing from 1850 to 2014. Despite not being explicitly constrained by present-day species distributions, the emergent biogeography from the model strongly resembles the true distribution of reef-building corals in the Indo-Pacific and NW Atlantic (figure 2). The simulated latitudinal distribution and limit of tropical corals generally corresponds well to the extent of coral reefs in the Allen Coral Atlas (*28*) for the major coral reef fronts in Japan, Florida, and Australia, with some tropical coral cover predicted beyond the known distribution of coral reefs in South Africa (figure S1). The ‘patchiness’ of coral reefs tends to be underestimated by CERES as geomorphology and reef accretionary processes are not considered, resulting in some erroneous predictions such as high coral cover across the sandy Bahama Banks (figure 2(c,d)), and overly continuous fringing reefs along continental coastlines. CERES significantly overestimates the abundance of corals along eastern ocean boundaries, the Southwest Atlantic, and upwelling equatorial regions (figure S1). These discrepancies are likely due to coral evolutionary history, and the effects of nutrients and/or sedimentation (see supplementary materials). Since corals in these environments represent less than 1% of global coral cover (*29*), we exclude them from our analyses and focus on the Indo-Pacific and NW Atlantic.

**Figure 2:**
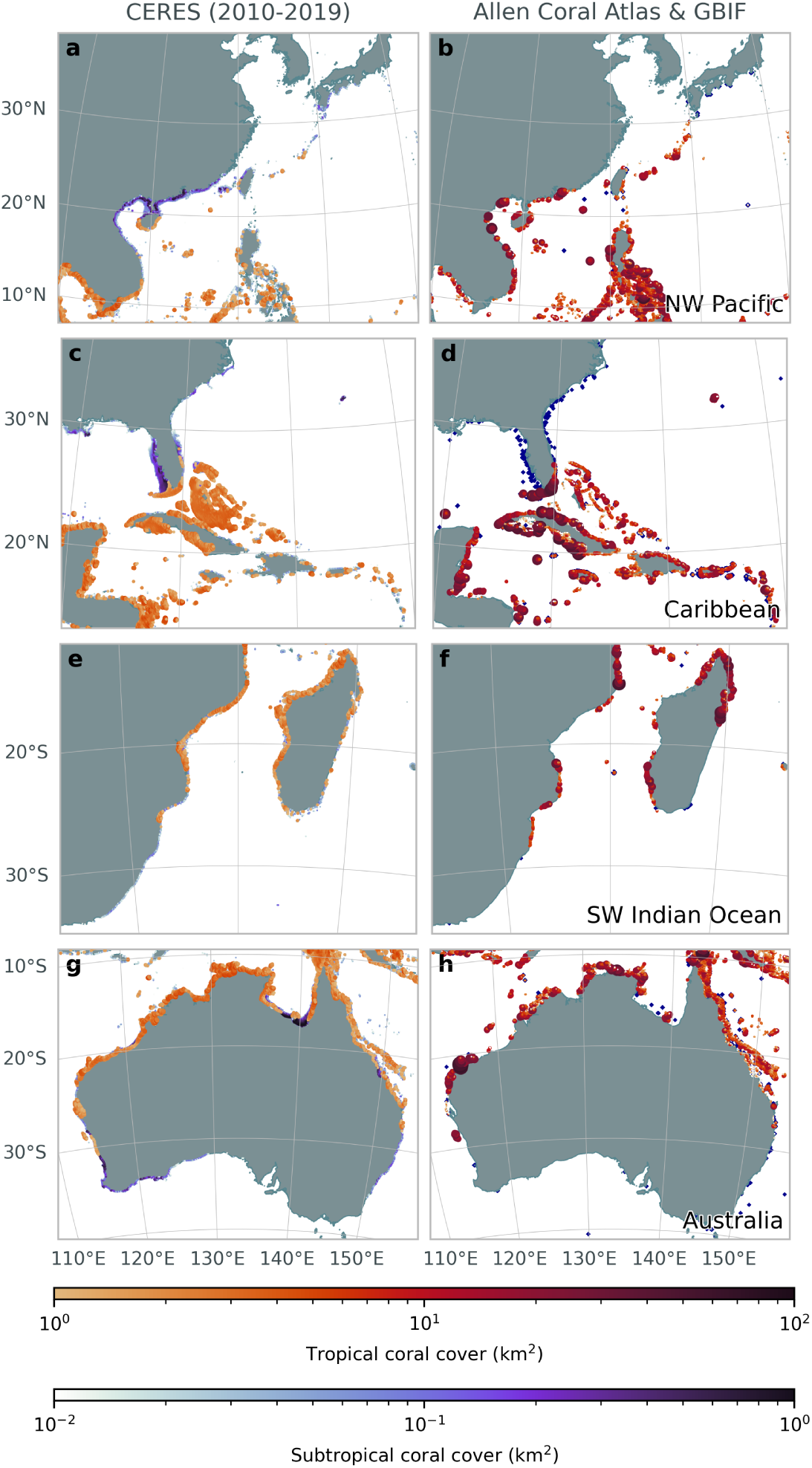
Mean coral cover from 2010-2019 across all ensemble members in CERES (*left*); and coral cover from the Allen Coral Atlas (*28*) and (as blue points) scleractinian coral occurrence records within 20 m depth (*30*) (*right*). To show both tropical and subtropical coral cover data from CERES, we only plot tropical coral cover where it exceeds 1 km^2^.

CERES also successfully reproduces both regional and global declines in coral cover observed since the late 20^th^ century by the Global Coral Reef Monitoring Network (*29*), although is on average slightly more conservative (figure S2). For instance, the GCRMN estimated that global live hard coral cover declined by 3.2% (in absolute terms) between the 2005-2009 and 2015-2019 monitoring periods, compared to 2.0% (1.0-3.4% across the model ensemble) for CERES. CMIP models are not assimilative, so CERES would not be expected to reproduce the correct timing of global bleaching events, but CERES nevertheless simulates lower coral cover variability compared to observations (figure S2). Whilst records of recent coral range expansion are limited, coral community composition in Japan has been periodically monitored since the 1930s (*15*). It is unclear to what extent observed changes represent a genuine range shift as opposed to an expansion of existing high-latitude coral communities (*17*), but most CERES ensemble members predict an increase in coral cover at sites where coral communities may have expanded over recent decades (figure S3), in good agreement with observations (*15*).

The ability of the model to largely independently (on the basis of empirical physiological data) reproduce coral reef distributions and decadal trends in coral cover suggests that it reasonably represents coral population dynamics, indicating that CERES may have predictive capacity for future trends.

### The future of coral reefs under anthropogenic climate change

At the end of the CMIP6 historical period, we impose SST and pH from the ‘sustainable’ SSP1-2.6, ‘middle of the road’ SSP2-4.5, and ‘fossil-fuelled development’ SSP5-8.4 ScenarioMIP scenarios (*26*) until the end of the 21^st^ century, again from twelve CMIP6 models. CERES predicts that coral cover declines by an average of 58% by the end of the 21^st^ century relative to 2010-2019 under SSP2-4.5 (figure 3(a)). There is considerable variability in the predicted decline across the CMIP6 ensemble, ranging from 41 – 71%. Although there is geographic variability, this decline exceeds 25% at almost all locations (figures 4(a), S4).

**Figure 3:**
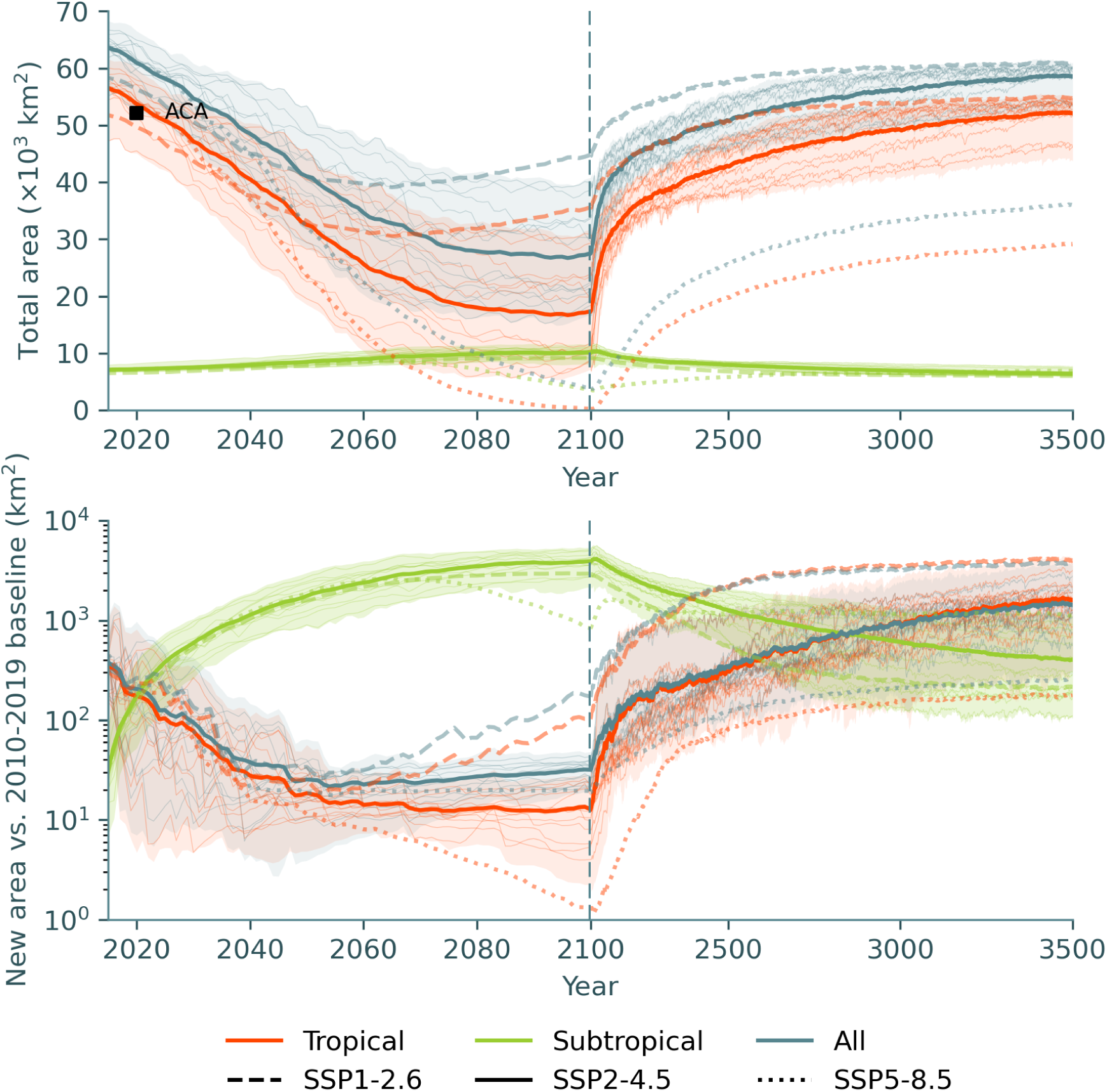
(a) Total modelled area of all (blue), tropical (orange), and subtropical (green) corals across the Indo-Pacific and NW Atlantic. Thick lines show the ensemble mean for SSP2-4.5 (solid), SSP1-2.6 (dashed), and SSP5-8.5 (dotted). Thin lines and the shaded area show trajectories for individual models for SSP2-4.5. The corresponding coral cover from the Allen Coral Atlas (*28*) is marked as a black square for reference, but note that the model habitable fraction parameter, *r_f_*, was tuned for this fit. (b) Total new coral cover relative to the mean model state from 2010-2019 (the sum across all sites where coral cover increased). New coral cover is high during the 2010-2019 baseline due to interannual variability. Note that the time axis scale changes in the year 2100, exaggerating the post-2100 recovery.

**Figure 4:**
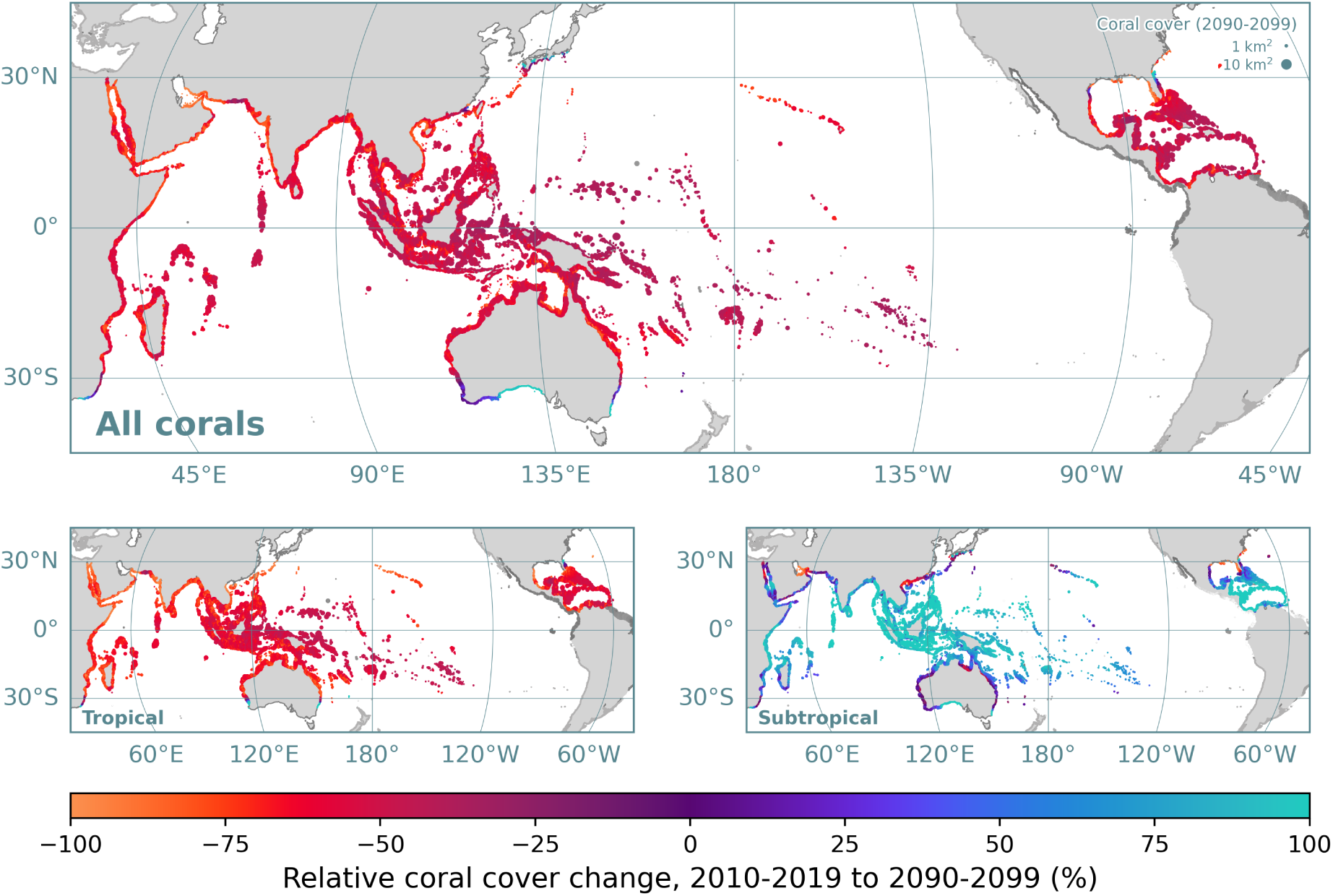
Mean relative change in coral cover between 2010-2019 and 2090-2099 under SSP2-4.5 across all ensemble members in CERES for (a) all, (b) tropical, and (c) subtropical coral. Sites are scaled by the absolute coral cover between 2090-2099. Sites outside the Indo-Pacific and NW Atlantic that were simulated by CERES but not considered by this study are plotted in greyscale. For clarity, only sites with a final coral cover exceeding 0.001 km^2^ are plotted.

The ensemble-mean coral cover decline of 58% masks the considerably higher decline of 71% (49 – 86% across the ensemble) predicted for tropical assemblages under SSP2-4.5 (figures 3(a), 4(b)), with many present-day reefs experiencing near or complete collapse (figure S4, supplementary animation 1). In contrast, the model predicts that the spatial cover of subtropical coral communities will increase by 41% (24 – 59%) over the coming century (figure 4(c)).

This increase is largely in the form of tropical assemblages transitioning into subtropical assemblages (the model analogue for the degradation of diverse coral reefs into low-diversity coral communities) rather than the formation of new coral habitat. Under the ‘sustainable’ SSP1-2.6 scenario, CERES predicts an average decline of 25% (11 – 45%) for coral cover, rising to 34% (15 – 59%) for tropical coral assemblages. Unsurprisingly, the ‘fossil-fuelled development’ SSP5-8.5 scenario results in a catastrophic 92% (85 – 99%) fall in coral cover, and a virtual elimination (96 – 100%) of diverse coral reefs.

In contrast to predictions from some statistical models, CERES predicts almost zero range expansion for tropical coral assemblages. Compared to a 2010-2019 baseline, CERES predicts that new tropical coral habitat will amount to less than 25 km^2^ by the end of the century, which is 3 orders of magnitude lower than the estimated net decline (figure 3(a)). Most of this marginal simulated range expansion occurs in Australia (figure 4(b), figure S5), although two ensemble members predict greater expansion in the NW Atlantic, to the north of the Florida Reef Tract (figure S6). Simulated tropical coral expansion in the NW Pacific and SW Indian Ocean is negligible this century (figures S7-8). Range expansion is predicted for subtropical coral communities, particularly in southern Australia (figure S5). However, as stated above, the largest ‘expansions’ of subtropical coral communities are actually due to the degradation of higher diversity tropical coral reefs (figure 4(c)). As a result, the ensemble-mean predicts that the centroid of subtropical coral communities will shift equatorward over the coming century, with the tropical coral centroid generally remaining static or also shifting equatorward (figure S9).

In most ensemble members and regions, the coral cover decline plateaus around the year 2080 under the SSP2-4.5 scenario (figure 3(a)). If temperatures and pH are kept stable after 2100 (with natural variability), coral cover recovery takes place over centennial timescales, with similar total coral cover to the present achieved by the year 2500. Under SSP1-2.6, coral cover instead reaches a minimum around 2060, whereas coral cover reaches almost zero under SSP5-8.5 by the end of the 21^st^ century. Recovery timescales are similar for all scenarios, although this is unsurprising as we treat all scenarios similarly after 2100 (holding the mean temperature and pH to the 2090-2099 mean). Coral cover under SSP5-8.5 does not fully recover within the simulation timespan (by 3500), although post-2100 predictions for this scenario may not be meaningful, as the model does not include any fundamental changes in coral ecology and evolution that might result from a near-total loss of coral reefs.

Over centennial timescales, CERES predicts genuine latitudinal range expansion in tropical corals (figures 3, 5, S10-11). Most of this range expansion is predicted to occur in the NW Atlantic and Australia (figure S12), with a potential northward expansion of the Florida Reef Tract, increased coral cover on Bermuda, and considerable southward expansion on the west and east coasts of Australia. CERES also predicts some minor latitudinal range expansion for tropical corals in Japan and South Africa, but these changes are small compared to the NW Atlantic and Australia.

Finally, as well as changes in coral cover, the model predicts significant changes in community composition between the four assemblages considered in CERES (figure 6, supplementary animation 2). At both high and low latitudes, the environmental stress due to 21^st^ century warming causes the model to predict a general shift away from fast-growing tropical corals. Slow-growing (stress-tolerant) tropical corals dominate low-latitude assemblages under peak environmental stress, although the model also predicts an increase in subtropical assemblages due to reef degradation. This new community composition peaks after 2050, slightly earlier than the coral cover minimum. As coral cover plateaus and begins to recover, there is a strong rebound in community composition towards fast-growing, competitive corals over a timescale of approximately 100 years. These corals dominate tropical assemblages during the recovery and colonisation phase once warming ceases, and continue to dominate community composition for centuries. By the end of the simulation in 3500, community composition is similar to the early 21^st^ century, although with a still elevated proportion of fast-growing corals in the tropics.

## Discussion

### High latitude coral reefs are ineffective large-scale refugia for corals under anthropogenic climate change

The geological record shows that coral reef fronts advance and recede in response to long-term climate change (*9–11*). Indeed, CERES simulates range expansions for tropical corals over centennial timescales (figure 3), which is fast compared to the temporal resolution of many geological records (*21*). Our simulations are therefore broadly consistent with palaeoecological observations. However, our simulations also suggest that the observed and expected rate of ocean warming in the 21^st^ century exceeds the evolutionary adaptive capacity of corals, with a considerable gap emerging between the mean SST and coral thermal optima. Therefore, whilst significant range expansion of coral reefs may eventually occur, any new high-latitude reefs will emerge centuries too late to act as refugia for the diverse coral assemblages characteristic of modern tropical coral reefs, which face a potentially catastrophic decline within decades.

In CERES, the timescale of this long-term range expansion (coral cover growth beyond the early 21^st^ century baseline) is primarily driven by the maximum coral colony linear extension rate. Under very high growth rates, tropical coral assemblages reach their equilibrium range as early as 100 years after the cessation of warming. Conversely, this expansion requires over 1000 years for very slow colony growth rates (figure S13). This timescale is only sensitive to the effective fecundity, or the rate of population growth due to the new coral colony establishment, under exceptionally high values (figure S14). High additive genetic variance, which sets the rate of change of subpopulation thermal optima in response to selective pressure, also drives faster range expansion (figure S15). However, the sensitivity of the range expansion rate to the additive genetic variance appears lower than to the colony growth rate. These findings suggest that the establishment of tropical coral reefs at higher latitudes is limited by the capacity for initially-small subpopulations to grow, partly due to maladaption to local conditions, rather than being limited by larval dispersal.

Observations of major reef-building coral species apparently colonising high-latitude environments such as Japan have been cited as potential evidence for the capacity of temperate seas to act as large-scale refugia for tropical corals (*15*). Our results instead support the interpretation that these observations primarily represent a local expansion of existing high-latitude coral communities that are distinct from low-latitude coral reefs (*17*). It is therefore unrealistic to expect these environments to harbour coral diversity comparable to low-latitude coral reefs until well beyond 2100, by which time the tropics may have already conceivably experienced an irreversible loss in diversity.

In fact, far from acting as refugia for low-latitude corals, high-latitude coral reefs experienced some of the greatest declines in coral cover in our simulations, in agreement with predictions from a previous study (*31*). The simulated decline in coral reefs over the 21^st^ century (for sites beginning with *>* 10% tropical coral cover) was reasonably predicted by a linear combination of benthic light intensity and the amplitude of the seasonal cycle in sea-surface temperature with a mean absolute error of 9.7% (figure 7), both of which tend to increase with distance from the equator. It may be surprising that these environmental variables were more important in determining the simulated coral reef decline than, for instance, the change in temperature or pH (figure S16). However, the spatial variability in warming and acidification across the tropics and subtropics is relatively low, and in the case of warming, exceeds the rate of evolutionary adaptation everywhere. Therefore, whilst ocean warming drives *global* trends of coral reef degradation, other environmental factors may be key to determining regional reef resilience. Since seasonal variability in temperature and benthic light intensity tend to respectively increase and decrease with latitude, existing high-latitude coral reefs experience particularly high compound environmental stress under climate change, exacerbating their simulated decline. For instance, whilst CERES reproduces the recent observed expansion of high-latitude coral communities in mainland Japan (figure S3), it also predicts that this expansion may be transient (figure S7) (*32*). These results are complementary to a recent study predicting coral species richness under climate change using environmental niche models (*33*). This study predicted that coral species richness would not increase at high-latitude coral habitats under SSP1-2.6, but could increase in Australia under SSP5-8.5. Taken together with our results, where a dynamic model suggests that the rate of climate change under SSP5-8.5 far exceeds the capacity of corals to expand their range, it appears unlikely that high-latitude environments will see a meaningful increase in coral species richness over the coming century, regardless of the emissions trajectory.

Although hyperthermal events from the geological record are not ideal climatic or ecological analogues to the present day, the rate of warming during the Palaeocene-Eocene Thermal Maximum (PETM) may have intermittently resembled contemporary warming rates (*21*). Palaeontological evidence from the PETM suggests that a long-term expansion towards higher latitudes was preceded by greater extinction risk for high-latitude colonial corals (*34*), which is consistent with our model predictions. Evidence from the PETM also highlights the importance of non-thermal stressors in explaining the response of corals to rapid environmental change (*35*). Conversely, although the PETM severely reduced coral reef formation (*35*), the hyperthermal event did not stem the long-term trend of increasing coral diversity throughout the Cenozoic (*34, 35*). The considerable decline in coral cover predicted by CERES therefore does not necessarily imply a loss of diversity, although caution is required when interpreting these past changes due to the dramatically different climatological and ecological context (*35*).

### Decoupling between winter temperatures and the coral reef front

In addition to explaining simulated spatial patterns of reef decline over the 21^st^ century, nonthermal stressors also explain much of the variability in the simulated mobility of the latitudinal limits of reef formation, also known as the coral reef front, over centennial timescales and longer (figure 5). CERES predicts that the greatest long-term latitudinal expansion of tropical corals will occur in Australia, where the seasonal cycle in temperature is relatively low, allowing reefs to follow the migration of isotherms that set their present-day latitudinal limits. The predicted expansion of the Florida Reef Tract is more moderate, likely limited by the relatively large seasonality in surface temperature.

**Figure 5:**
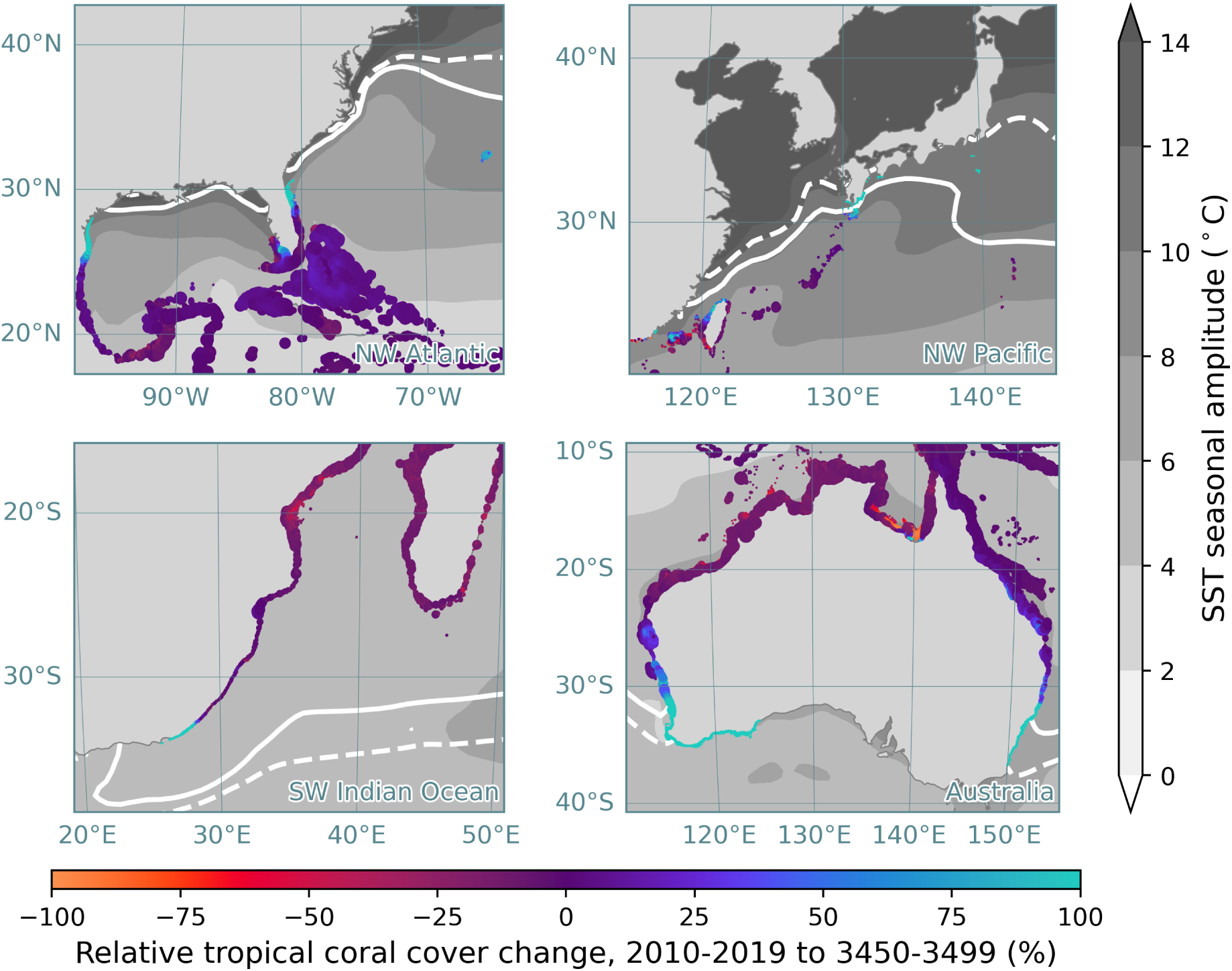
Mean relative change in tropical coral cover between 2010-2019 and 3450-3499 under SSP2-4.5 across all ensemble members in CERES for tropical coral assemblages. Sites are scaled by the absolute tropical coral cover between 3450-3499. The ocean is shaded by the amplitude of the seasonal cycle of sea-surface temperature, and the solid and dotted white contours represent the annual minimum 18°C isotherm for the pre-industrial and post-2090 periods, respectively. For clarity, only sites with a final coral cover exceeding 0.001 km^2^ are plotted.

This temperature variability is even more dramatic in Japan, where the seasonality is greater in amplitude, and the simulated expansion of tropical corals is small and largely limited to the northernmost Ryūkyū and Izu islands (figure 5). It has previously been suggested that tropical coral range expansion in Japan may be limited by ocean acidification (*32*). CERES simulations with pH fixed to pre-industrial levels suggest that acidification is largely responsible for the failure of some coral reefs to fully recover by the year 3500 (figure S17), since the fitness of corals with respect to pH is fixed in our model. In this scenario, tropical coral assemblages appear on the Pacific coast of Kyūshū, but range expansion otherwise remains limited (figure S18). This suggests that temperature variability may be a more important bottleneck than acidification in throttling the poleward expansion of tropical corals in Japan. Latitudinal light attenuation also limits coral reef range expansion in CERES. In simulations with reduced sensitivity to light, there is greater range expansion in all regions, particularly in South Africa and Japan (figure S19). Light intensity, whilst not being the only constraint on the latitudinal limit of tropical corals (*36, 37*), may therefore still play an important role.

Since CERES has only been assessed against short-term (sub-centennial scale) observations, these long-term predictions should be viewed with considerable skepticism. Nevertheless, these simulations support previous assertions that the coral reef front is not solely set by winter seasurface temperatures, but rather by a range of environmental parameters (*6, 38*).

### Disproportionate impact of climate change on acroporid corals may be transient

As with previous studies (*22, 31*), peak environmental stress during the 21^st^ century disproportionately affects fast-growing tropical corals in CERES, the model assemblage that is dominated by acroporid corals (*23*). Indeed, after individual mass bleaching events, coral community composition in reefs have been widely observed to shift away from (branching) acroporids and towards slow-growing corals with stress-tolerant life-history strategies (*23, 39*).

CERES predicts that this initial shift in community composition may be rapidly followed by a rebound *towards* fast-growing tropical corals (figure 6), which lead the recovery of low-latitude reefs from the end of the 21^st^ century. Acroporids proliferated during the Pleistocene, possibly due to their rapid growth rate, allowing them to efficiently shift their distribution in response to rapid sea level change (*40*). The proliferation of acroporid corals in response to the availability of new habitat is consistent with our model predictions: once fast-growing corals (primarily acroporids) are able to adapt to a warmer climate, their population size grows rapidly and allows them to dominate the recently depopulated tropical reefs. This ‘overcorrection’ persists for over 1000 years in our simulations, more than an order of magnitude longer than the initial decline. Based on these simulations, one could hypothesise that (1) the currently observed decadal shifts in coral community composition may be a poor indicator of the long-term future of coral reefs, and (2) the dominant geological signature of past rapid warming may be an increase in the abundance of acroporid corals in fossilised reefs, even if the initial response is the opposite. These are interesting hypotheses but, given the poor constraints on inter-assemblage competition in CERES, we leave these as open questions.

**Figure 6:**
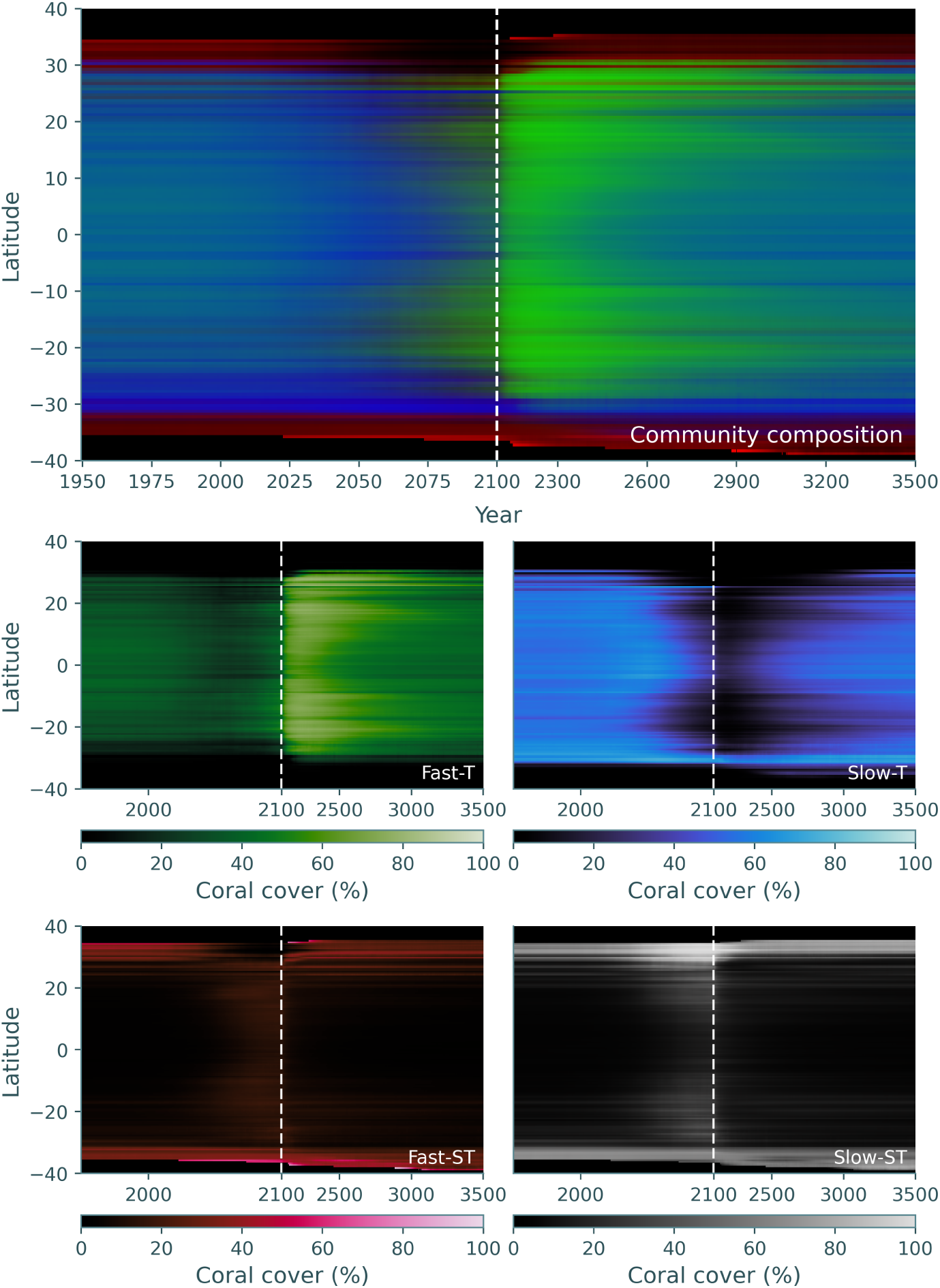
Mean community composition per latitude band (*y* axis) as a function of time (SSP2-4.5), with fast-growing tropical, slow-growing tropical, and fast-growing subtropical proportions broadcast to green, blue and red channels respectively (there are three degrees of freedom as proportions sum to 1, but slow-growing subtropical proportion is effectively represented by brightness).

### Limited scope for 21**^st^** century range expansion despite model uncertainty

CERES reasonably reproduces the present day distribution of corals and coral reefs (figure 2), as well as observed trends since the late 20^st^ century (figure S2-3), and most parameters are constrained by empirical data (see *Materials and Methods*). However, many parameters are nevertheless subject to considerable uncertainty. Here, we demonstrate that the key conclusion of this study - that range expansion alone is unlikely to buffer coral diversity against climate change by the end of the 21^st^ century - is robust with respect to this uncertainty.

The model assumes that all coral colonies follow the same log-normal size distribution, which does not evolve through time. This is a simplifying assumption, as colony size distributions change through space (*41*) and time (*42*). However, the growth rate of coral cover has a relatively low sensitivity to the colony size distribution (see supplementary materials, figure S20). In particular, since high-latitude reefs tend to support larger coral colonies (*41*) (and therefore slower coral cover growth rate), uncertainty in colony size structure does not translate into major uncertainty in range expansion potential over the coming century.

As previously described, light limitation and acidification contribute to the limited capacity for range expansion for tropical corals simulated by CERES. However, although a lower physiological sensitivity to light intensity and pH would permit greater range expansion, new reef habitat remains low (≪ 100 km^2^, considerably lower than the simulated early 21^st^ century interannual variability) for all values tested (figure S21-22). A broader thermal tolerance and more relaxed heat stress threshold and sensitivity result in delayed coral cover decline and more new reef habitat, but the effects are quite small (figure S23-25).

The additive genetic variance (with respect to the coral thermal optimum) is poorly constrained, although a growing number of studies have found phenotypic variance that translates into additive genetic variance *V* of order 0.05°C^2^ (*43, 44*). Greater additive genetic variance reduces the predicted fall in coral cover over the 21^st^ century (figure S26), due to faster evolution. More coral habitat is generated under both very high and low additive genetic variance, although the latter is likely an artefact of the model initial conditions (the spin-up timescale used in this study may be insufficient for *V ∼* 0°C^2^). The most extreme case (*V* = 0.5°C^2^) increases the potential for new coral cover by an order of magnitude relative to the base case we consider in this study. However, a very high value of *V* = 0.5°C^2^ is inconsistent with empirical data (*43, 44*).

The competition parameters in CERES that govern interactions between coral assemblages are one of the few parameters that are not explicitly based on empirical data. Values qualitatively similar to those chosen in our simulations are required to reproduce the observations that (1) corals of different life-history strategies coexist at most latitudes (*45*); (2) coral cover in marginal coral communities is considerably lower than coral cover in reefs; and (3) ‘tropical’ (high-diversity, reef-forming) coral assemblages dominate over ‘subtropical’ non-reefal coral communities at low latitudes. At least within the CERES modelling framework, competition parameters that fulfil the above criteria will tend to produce qualitatively similar (if quantitatively different) behaviour to figure 6. There was also no empirical data to constrain the difference in behaviour under cold stress between fast-growing and slow-growing assemblages, although the model is fairly insensitive to these parameters (figures S27-30). There is some evidence to suggest that acroporid (broadly fast-growing) corals may be more sensitive to changes in pH (*46*), potentially exacerbating the shifts in community composition predicted by CERES. However, this was not incorporated in the present study due to insufficient evidence, and we would not expect them to alter our conclusions given the secondary role acidification plays compared to warming (*7*).

Our sensitivity analysis shows that a key parameter setting the resilience of tropical corals in response to 21^st^ century warming is the effective fecundity. Although higher effective fecundity appears less important in setting the rate of tropical coral range expansion beyond 2100 (figure S14), it does increase the new coral cover generated during the 21^st^ century (figure S31). Effective fecundity is a complex parameter, translating the existing coral cover into new coral cover generated through the production of larvae, thereby incorporating the actual fecundity (eggs generated per polyp), fertilisation rate, larval mortality, settlement likelihood, and survival to sexual maturity. The parameters used in this study represent a best estimate based on values reported in the literature (*23, 47, 48*), and considerably higher values of fecundity are inconsistent with the present-day distribution of coral reefs, particularly in the NW Atlantic and Australia (figure S32-33). However, although these parameters may reasonably represent coral assemblages as a whole, it is possible that individual species (with particularly high biological fecundity, and/or very low larval and juvenile mortality) may avoid major decline this century.

We found no meaningful correlation between any connectivity metric and simulated coral reef trajectories (figure S34). This contrasts with previous studies that suggested the longdistance dispersal of coral larvae across thermal gradients could, in some cases, improve the capacity of coral reefs to rapidly adapt to ocean warming (*22, 49*). We hypothesise that the simpler fitness landscape and annual (as opposed to monthly) mean SST considered by previous studies may have inflated the signal of long-distance connectivity on coral reef trajectories. Our findings suggest that evolutionary processes at a reef scale, perhaps driven by intra-reef connectivity between small-scale thermal regimes (*50*), may play a more important role in shaping the response of coral reefs to short-term environmental change than long-distance connectivity.

Finally, CERES does not comprehensively represent all processes that may be relevant to the dynamics of a coral reef, including sea level rise (*51*), settlement cues and substrate suitability (*52, 53*), interactions with macroalgae and temperate biota (*6, 54*), and unresolved scales of oceanographic variability causing our predictions of long-distance connectivity to represent upper bounds (*55,56*). However, these unrepresented processes would all generally be expected to *further* inhibit range expansion, and therefore do not affect the primary conclusions of this study.

In summary, this comprehensive sensitivity analysis demonstrates that it is highly likely that tropical coral assemblages as a whole will (1) face a major decline this century under realistic emissions scenarios, and (2) fail to expand their range rapidly enough to buffer diversity against this decline. However, our findings also suggest that certain coral species may avoid catastrophic declines, particularly those with very high effective fecundity, rapid growth rates, and high heritability and phenotypic variance in thermal tolerance.

### Emissions cuts and management are critical for the future of coral reefs

Despite high-latitude seas still frequently being referred to as potential refugia, our conclusion to the contrary will be unsurprising for many ecologists, given the abundance of evidence that the functioning of marginal coral communities is fundamentally distinct from tropical reefs (*18, 19*). What may be more surprising are our pessimistic projections for high-latitude coral communities and subtropical reefs in general. Although these ecosystems lack the diversity of tropical coral reefs (*57*), they nevertheless provide important ecosystem services, and the proactive implementation of management strategies may be essential to mitigate the impacts of climate change on these unique communities (*18*). In particular, despite the recent success of corals in some high-latitude communities in Japan (*15, 16*), this success may be transient and should not be misinterpreted by policymakers as evidence that management is low priority. Indeed, most recent studies exploring the potential of high-latitude environments in Japan to act as refugia for corals emphasise the pressing need for active management to conserve these communities (*16, 17*).

Although we did not consider anthropogenic stressors such as overfishing and pollution in our simulations, our results suggest that the presence or absence of non-thermal stressors (such as light limitation) that reduce the baseline growth potential for corals can be the difference between tolerable and catastrophic coral cover decline (figure 7). Indeed, coral reefs in Hawai’i exposed to less pollution and greater fishing restrictions have demonstrated greater resilience to thermal stress (*58*), so reducing non-thermal stressors where possible will be increasingly essential for limiting unmitigable damage from heat stress.

**Figure 7:**
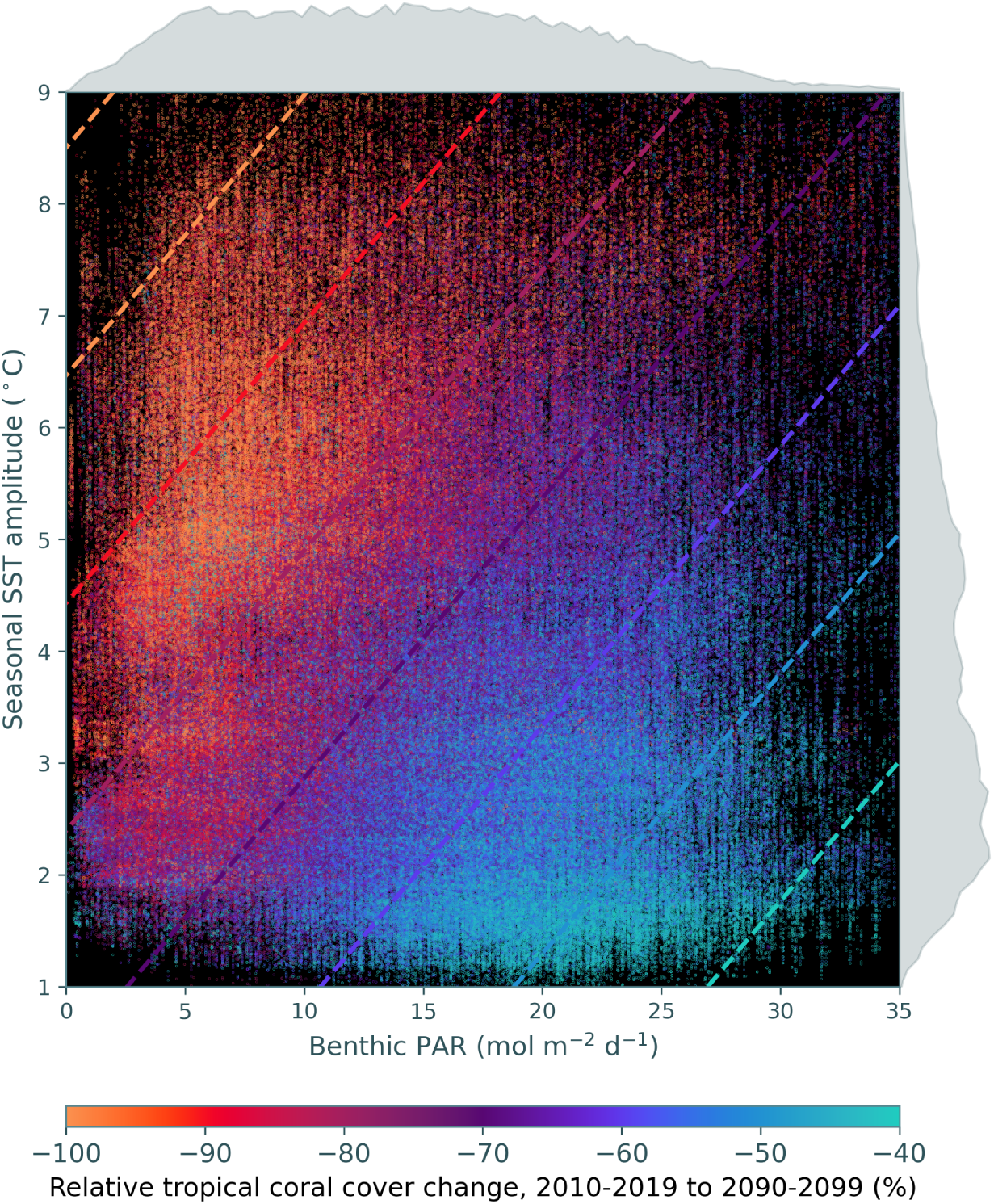
Predicted (contours) and simulated (points) relative change in tropical coral cover over the 21^st^ century, as a function of mean benthic light intensity and the amplitude of the seasonal cycle in sea-surface temperature. Only sites where the tropical coral cover originally exceeded 10% are included. The predicted change in coral cover was calculated from a linear mixed model, with random intercepts varying by ensemble member. The mean absolute error is 9.7%, and the sensitivity of the change in tropical coral cover with respect to PAR and seasonal SST amplitude is 1.2%/mol m*^−^*^2^ d*^−^*^1^ and -4.9%/°C respectively. Simulated data are plotted with random effects removed. Histograms show the distribution of predictor variables across reef sites.

Finally, and perhaps most importantly, our results underline the sensitivity of the future of tropical coral reefs to greenhouse gas emissions trajectories over the 21^st^ century. The geological record demonstrates that a major reduction in tropical coral reef formation is not necessarily equivalent to a reduction in coral diversity (*34, 35*). Likewise, the proposed inability of high-latitude seas to buffer coral diversity against climate change does not preclude the existence of other refugia, such as microrefugia or mesophotic reefs (although, similarly to the high-latitudes, these environments are also home to functionally distinct communities (*5, 19*)). Nevertheless, diversity considerations aside, the predicted differences between coral reef futures under SSP1-2.6, SSP2-4.5, and SSP5-8.5 are considerable (figure 3), and these differences will have an enormous impact on the communities that depend on healthy coral reefs for their livelihoods (*2*). Our results support a growing body of evidence (*22, 31, 33, 59*) that any reduction in emissions will be highly consequential for coral reefs, even if specific climate targets are missed. These impacts will shape not only coral reef trajectories but also the lives of those who depend on them well beyond the end of the century.

## Materials and Methods

### Model overview

CERES (Coral Eco-evolutionary Range Expansion Simulation) simulates eco-evolutionary and range expansion dynamics for shallow-water corals under climate change using a simple metacommunity framework, expanding on earlier studies (*22, 60*). For a set of *i* discrete subpopulations and *j* competing coral groups over *k* time-steps, CERES simulates the response of live coral cover, *C_ijk_* to sea-surface temperature (*T_ik_*), pH (*P_ik_*), and benthic light intensity (*I_ik_*) (table 1). We model a single quantitative trait under selection, the population-mean thermal optimum *z_ijk_*, with additive genetic variance *V_j_*. We consider two groups of corals: ‘tropical’ corals, representing the high-diversity coral assemblages characteristic of low-latitude coral reefs, and ‘subtropical’ corals, representing the low-diversity coral assemblages characteristic of high-latitude non-reefal coral communities. We emphasise that, although our tropical coral group is analogous to coral reefs in the present day, CERES does not consider the process of reef accretion (*59*), and we use the terms ‘tropical’ and ‘subtropical’ as a descriptive generalisation rather than precise geographic designations. Similarly to previous studies (*22, 31, 59*), we model two competing life-history strategies for ‘tropical’ corals - fast-growing (**Fast-T**), and slow-growing (**Slow-T**) - and likewise for ‘subtropical’ corals (**Fast-ST** and **Slow-ST**). We assume that fast-growing life history strategies have greater maximum growth rates, lower thermal stress thresholds, and greater dispersal potential. Based on the observed latitudinal distribution of tropical coral reefs and non-reefal coral communities, we parameterise tropical coral assemblages as having narrower thermal tolerance and stress thresholds, but a competitive advantage over subtropical coral assemblages under optimal conditions (table 2). We consider anywhere within 20 m depth and within 45° of the equator to be ‘potentially habitable’ for shallow-water corals, and assume that reef can cover at most a certain fraction *r_f_* of this area. We set 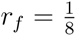, based on comparison with the Allen Coral Atlas (*28*).

**Table 1:**
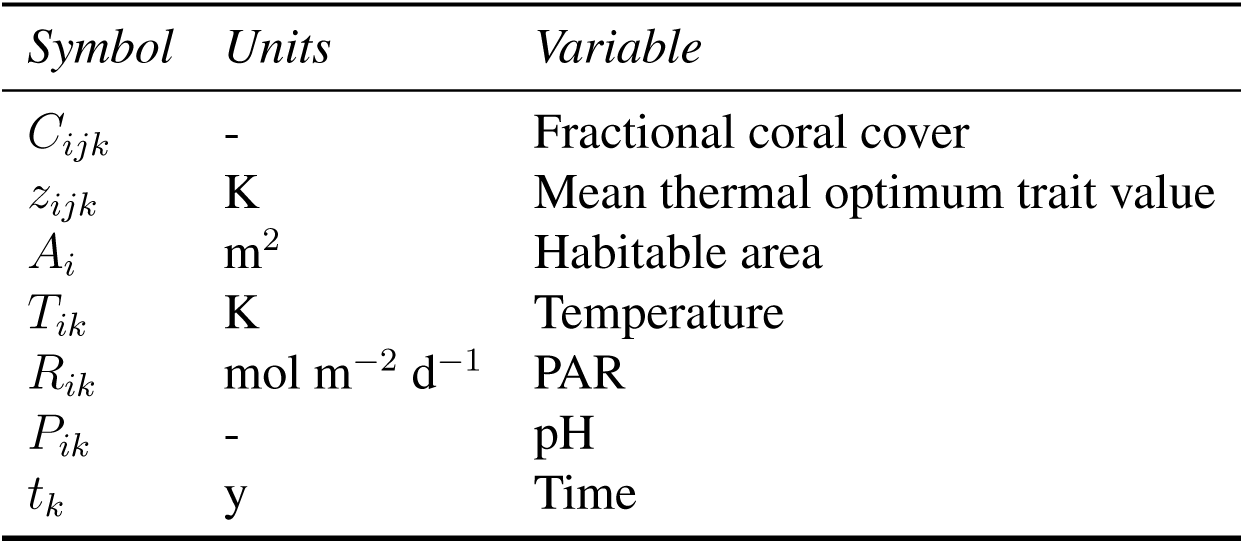
Model variables. Intermediate variables used solely for computation (e.g. *g_ijk_*) are not listed.

| *Symbol* | *Units* | *Variable* |
| --- | --- | --- |
| $C_{ijk}$ | - | Fractional coral cover |
| $z_{ijk}$ | K | Mean thermal optimum trait value |
| $A_i$ | m<sup>2</sup> | Habitable area |
| $T_{ik}$ | K | Temperature |
| $R_{ik}$ | mol m<sup>-2</sup> d<sup>-1</sup> | PAR |
| $P_{ik}$ | - | pH |
| $t_k$ | y | Time |

**Table 2:**
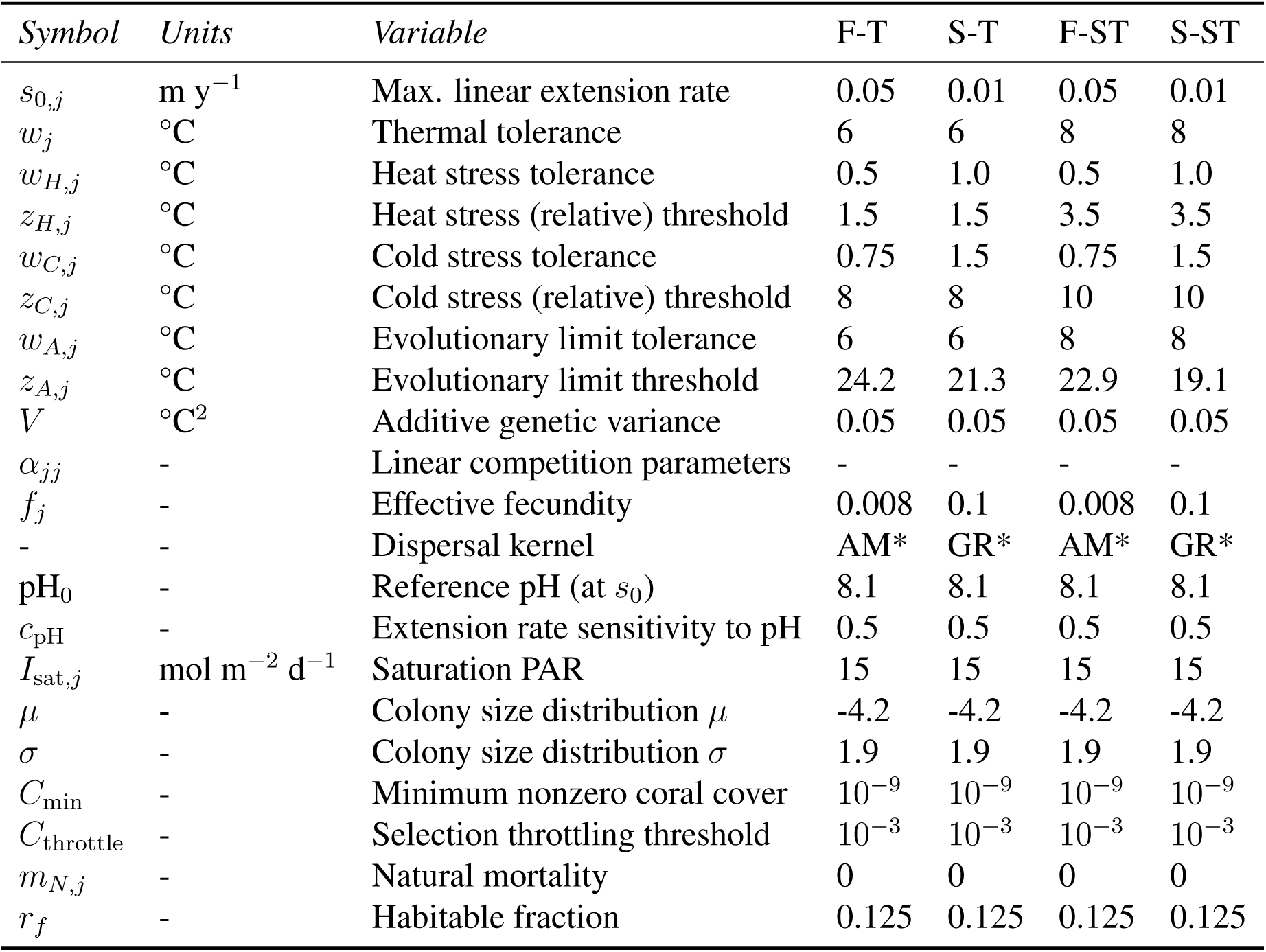
Model parameters. F-T: Fast-growing tropical coral, S-ST: Slow-growing subtropical coral, and so on.

### Experimental design

For each CMIP6 model used, we initialize coral cover from the Allen Coral Atlas (*28*) and the thermal optimum trait value to the 90th percentile sea-surface temperature from the preindustrial control run (*piControl*). This accelerates model convergence, but the convergent state is insensitive to the initial conditions. We spin up the model for 5000 years by looping the *∼*500 year *piControl* forcing, as the model reaches a steady state in terms of coral cover and community composition after approximately 1000-4000 years (figures S35-36). This ensures that any response to imposed historical or projected forcing is genuinely a response to a change in forcing, rather than an adjustment from the initial conditions. We then sequentially impose *historical* (1850–2014) and *SSP1-2.6*, *SSP2-4.5* or *SSP5-8.5* (2015–2100) forcing. For validation purposes, we extend the CMIP6 historical run to 2020 using *SSP2-4.5* forcing, since this more realistically represents true emissions (*61*). Finally, we extend the ScenarioMIP forcing beyond 2100 by computing the difference between *piControl* and 2090-2100 sea-surface temperature and pH, and adding this to *piControl* forcing. This is not intended to represent a realistic post-2100 climate trajectory (all *SSP2-4.5*, and most *SSP1-2.6* and *SSP5-8.5* simulations were not extended beyond 2100 in CMIP6), but does permit us to investigate the long-term equilibrium response of corals to anthropogenic warming.

We assess the model predictive capacity by comparing (i) modelled areal reef-forming coral cover averaged across 2010-2020 against the Allen Coral Atlas; (ii) modelled coral cover trends from 1978-2019 against compilations across Global Coral Reef Monitoring Network regions (*29*); and (iii) comparing hindcast changes in coral cover in Japan against observations of new species occurrences (*15*).

### Environmental data

At the time of writing, twelve models contributing to CMIP6/ScenarioMIP6 have sea-surface temperature and pH data at monthly temporal resolution, covering *piControl*, *historical*, *SSP1-2.6*, *SSP2-4.5* and *SSP5-8.5* scenarios, and with a transient climate response within a range deemed reasonable, 1.2 – 2.4°C (*27*). These models are CESM2, CESM2-WACCM, CMCC-ESM2, CNRM-ESM2-1, GFDL-CM4, GFDL-ESM4, IPSL-CM6A-LR, MIROC-ES2L, MPI-ESM1-2-LR, MPI-ESM1-2-HR, NorESM2-LM, and NorESM2-MM. We do not screen models based on pH since there is less variability between models (*62*). We use model output on the native grid where possible.

Due to the higher resolution of the CERES grid versus the ocean grids used in models contributing to CMIP (ranging from c. 20 km for GFDL-CM4, to c. 100 km for MPI-ESM1-2-LR), there is mismatch between the land-sea masks. We therefore interpolate land cells in the CMIP models to ensure data coverage for all CERES cells. We then compute the sea-surface temperature and pH for all CERES cells using nearest-neighbour interpolation. Finally, we attempt to correct for time-invariant biases in sea-surface temperature and pH using the ‘delta method’ (*63*), i.e., computing the difference between monthly climatological sea-surface temperature and pH from 1986-2014 for CMIP fields and observational products, and then adding this (monthly climatological) offset to CMIP fields. We use sea-surface temperature from the 5 km NOAA Coral Reef Watch CoralTemp product (*64*), and pH from the CMEMS Global Ocean Surface Carbon product (*65*).

We use sea-surface temperature rather than benthic water temperature due to the coarse horizontal resolution of the CMIP models used in this study. The average depth of coastal cells used in these GCMs is significantly deeper than the allowed depth for ‘habitable’ cells in CERES, so sea-surface temperature is therefore likely a better approximation for the water temperature experienced by corals.

We obtain benthic PAR from an observational product (*66*), which estimates monthly climatological benthic PAR based on surface irradiance, water-column light attenuation from ocean colour, and bathymetry. This product is made available at the same resolution as GEBCO 2023 (i.e. 15×15 arc-seconds). We firstly interpolate any missing values using the `setmisstodis` distance-weighted interpolation algorithm in cdo, and then compute the benthic PAR for each CERES cell as the mean across all benthic PAR values within a water depth of 20 m. We note that, due to the nonlinear (exponential) decay in PAR with depth, computed benthic PAR values based on the mean depth will likely be an overestimate compared to the true mean benthic PAR.

### CERES ecological dynamics

As a reminder, the subscripts *i*, *j* and *k* respectively refer to subpopulations, coral groups, and time steps. We assume that live coral cover changes through the growth of existing corals through budding, the establishment of new coral colonies through sexual reproduction, and mortality. During months without spawning, the rate of change of coral cover is given by the following differential equation:

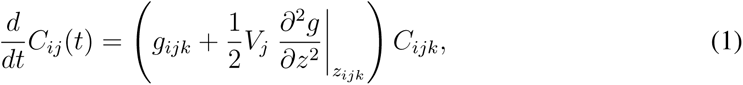

where *g_ijk_*is the fitness function (y*^−^*^1^), and 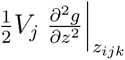 is the genetic load due to variability in *z* around *z* = *z_ijk_*.

The fitness function is in turn defined as:

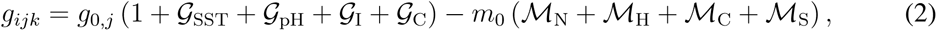

where *g*_0_*_,j_* and *m*_0_*_,j_* (y*^−^*1) are respectively the growth and mortality rate coefficients, and *𝒢* and *ℳ* terms (unitless) respectively determine the environmental dependence of coral cover growth and mortality. We assume that the linear extension rate of corals scales with the sum of the effects of temperature (𝒢_SST_), pH (𝒢_pH_), benthic PAR (𝒢_I_), and competition (𝒢_C_), i.e., there are no interactive effects. We define these functions as follows:

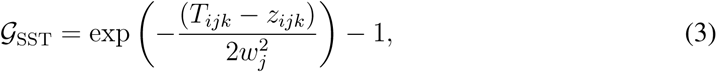

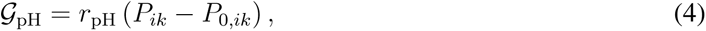

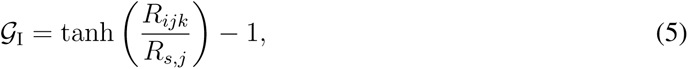

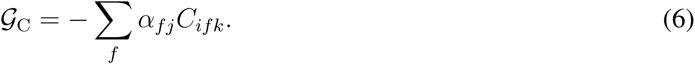

The Gaussian form of the temperature function is based on previous coral metapopulation studies (*22, 49*) and coral thermal performance curves (*67*). We set the reef-forming coral thermal tolerance breadth *w_j_* to 6°C, consistent with empirical data (*67, 68*). Since subtropical/temperate coral communities can survive a higher amplitude seasonal cycle, we add +2°C to *w_j_* for the subtropical-type corals. We assume that the coral extension rate responds linearly to pH, such that a reduction in 1 pH unit reduces the extension rate by 50% (*46*). Coral calcification rate is widely modelled as a hyperbolic tangent response to light intensity (*69, 70*), with a saturation value of *R_s_*. Here, we set *R_s_* = 15 mol m*^−^*^2^ d*^−^*^1^, a lower-bound estimate from (*70*). Although some short-term empirical studies found a positive growth rate in dark conditions (*70*), in this study, we assume that shallow-water corals cannot grow in complete darkness. We do not consider the potential for photoacclimation, photoinhibition, or shifts in energy sources. Finally, we model competition as a linear interaction between coral groups (*22*), set by a competition matrix *α*, where *α*_T,ST_ = 1.2, *α*_ST,T_ = 0.8, *α*_T,T_ = 2, and *α*_ST,ST_ = 10. In other words, when all other parameters are equal, tropical (T) corals have a competitive advantage over subtropical (ST) corals. These competition parameters reproduce the observations that (i) differing life-history strategies coexist across the latitudinal range of corals (*45*), and (ii) despite the greater environmental tolerance required to survive at higher latitudes, non-reefal coral communities exist at relatively low abundance at high latitudes, and reef-associated communities dominate in the tropics (*57*). It is possible for 𝒢_SST_ +𝒢_pH_ +𝒢_I_ +𝒢_C_ *< −*1, representing compound environmental stress. In this case, the negative value can be interpreted as additional mortality.

To convert coral colony linear extension rates into a coral cover growth rate, we assume that coral colony size structure is log-normal, as is commonly observed in coral reefs (*25*), with shape parameters *µ_j_* and *σ_j_*. In this case, 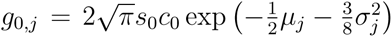, where *s*_0_ is the optimal linear extension rate, and *c*_0_ = 1 m*^−^*^1^ (see supplementary materials). We set *s*_0_ to 5 mm y*^−^*^1^ and 1 mm y*^−^*^1^ for fast-growing and slow-growing corals, the average across their respective life history strategies (*23*). We also set *µ* = *−*4.2 and *σ* = 1.9 for all corals, based on the geometric mean across a range of tropical coral species (*25*). To avoid requiring a complex size-structured model, we assume that these shape parameters do not vary, although acknowledge that this is a model limitation that could be improved in the future (*41, 42*).

We assume that the mortality rate of corals scales with the sum of the effects of natural background mortality (*M*_N_), heat stress (*M*_H_), cold stress (*M*_L_), and an absolute lower thermal limit (*M*_A_). We define these functions as follows:

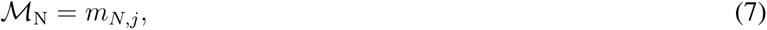

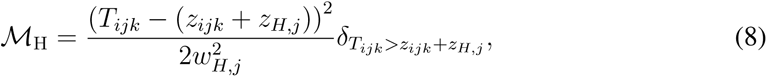

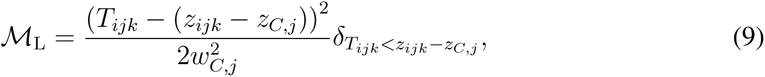

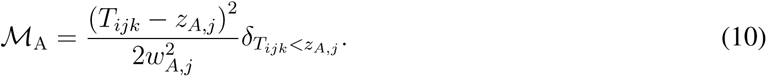

Here, *δ_X_*_=1_ if *X* is true, and 0 otherwise, such that *M*_N_, *M*_H_ and *M*_L_ are only nonzero under stress. We set *m*_0_*_,j_* = 1 y*^−^*^1^ for correct units (no scaling is required, contrary to the extension rate terms, assuming these mortality rates refer to coral cover). In this study, we set *m_N,j_*= 0, which assumes that all mortality is due to environmental stress. We also define the temperature difference between the thermal optimum, *z_ijk_*, and the onset of heat-stress and cold-stress induced mortality as *z_H,j_* and *z_C,j_*, respectively. The heat stress threshold for corals is often set as the mean monthly maximum (MMM) sea-surface temperature + 1°C (*64*), so we set *z_H,j_*= 1.5°C for reef-forming corals since the thermal optimum in our simulations usually equilibrates slightly below the MMM. There are fewer empirical constraints for *z_c,j_*, so we set this value to 8°C for reef-forming corals, such that *z_C,j_* + *z_H,j_* is less than the amplitude of the seasonal cycle at most tropical coral reefs (*38*). As above, subtropical and temperate coral communities survive a greater range of temperatures than their tropical counterparts, so we add +2°C to both *z_H,j_* and *z_C,j_* for subtropical-type corals. The parameters *w_H,j_* and *w_C,j_* are directly related to the accumulated heat stress required to cause significant coral mortality (see supplementary materials). Since fast-growing corals such as *Acroporidae* tend to experience greater bleaching and mortality during marine heatwaves (*39*), we set *w_H,j_* to 0.5°C and 1.0°C for fast-growing and slow-growing corals, equivalent to mass mortality thresholds of around 9 and 18 DHW, respectively (*39, 71*). Mortality due to cold stress is comparatively poorly constrained, but we set *w_C,j_* to 0.75°C and 1.5°C for fast-growing and slow-growing corals respectively, equivalent to conservative mass mortality thresholds of around 13 and 27 DCW (*72*). Finally, although we do not explicitly constrain *z_ijk_*, as this trait is free to evolve, we add a fitness penalty below an absolute lower thermal limit *z_A,j_*(*22*), with tolerance *w_A,j_* (°C). In the absence of better constraints, we set *w_A,j_* = *w_j_*, and set *z_A,j_* such that the minimum temperature at which growth is possible is 19°C and 16°C for reef- and community-type corals, slightly above the winter temperatures where coral reefs and communities are presently found (*57*). The population growth rate for all four modelled coral groups (under otherwise optimal conditions) as a function of the temperature and trait value is shown in figure 8.

**Figure 8:**
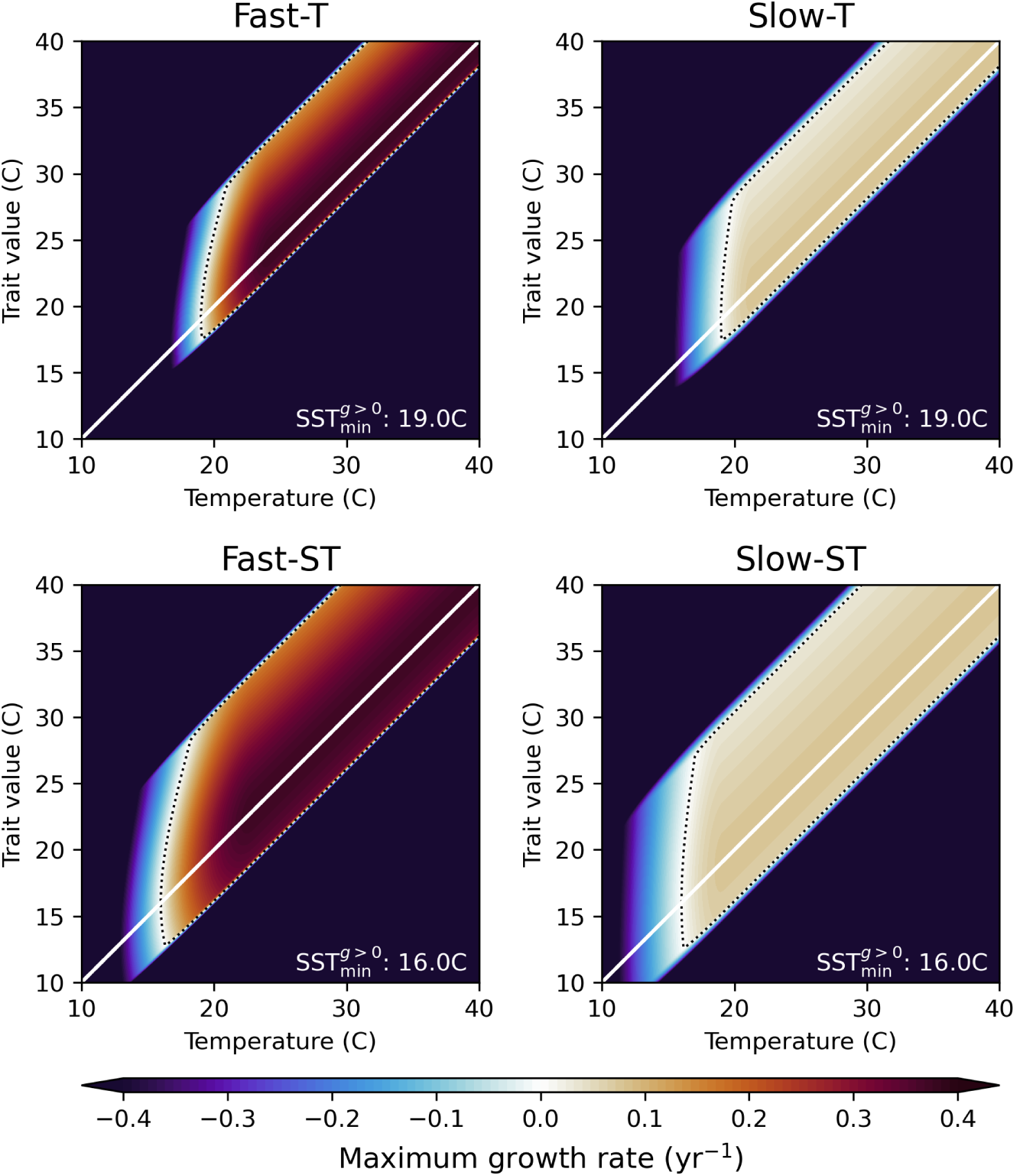
Optimal population growth rate as a function of the monthly mean temperature (*x* axis) and the thermal optimum trait value *z* (*y* axis). T: Tropical, S: Subtropical. The dotted black line bounds the region where positive growth is possible, and the white line corresponds to the temperature and thermal optimum being equal. The absence of positive growth rates at low temperatures is due to the absolute limit imposed in the model configuration.

During months with spawning (taken to be once per year), we add an additional growth term due to the establishment of new coral colonies. We assume that fecundity is time-invariant; that larval settling likelihood is proportional to the available ‘free space’ not occupied by corals and is not affected by chemical cues; that all settling takes place over a ‘short’ time period (such that settling dominates changes in *C_ij_* during a settling event); and that a newly established colony has the size of a single polyp. Under these assumptions, there is an analytical solution for the change in coral cover due to a spawning event (see supplementary materials):

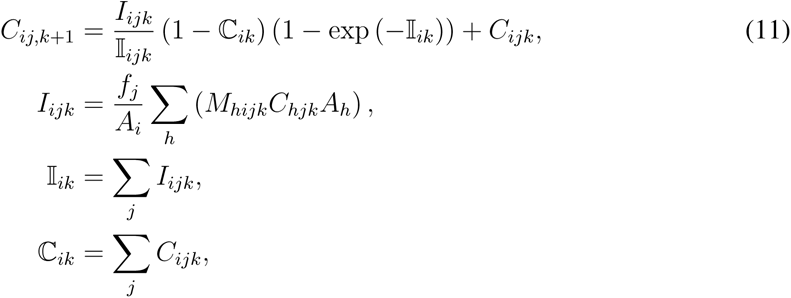

where *f_j_* is the effective fecundity (unitless) and *M_hijk_* (also unitless) is the potential connectivity between sites *h → i*, i.e., the likelihood that a coral *j* larva generated at site *h* is transported to site *i*. We set the effective fecundity to 4 *×* 10*^−^*^4^ and 5 *×* 10*^−^*^3^ for fast-growing and slow-growing corals respectively, based on the mean number of eggs generated per polyp for each life history strategy (*23*), and conservative assumptions for the proportion of eggs that translate into sexually mature corals (*47, 48*) (see supplementary materials). The model assumes all coral is capable of both sexual and asexual reproduction. The onset of sexual reproduction typically occurs 1-10 years post-settling (*73*); neglecting this delay may cause the model to overestimate range expansion.

The potential connectivity matrix is based on the *EZfate* product (*24*), advected using the 1/12° CMEMS GLORYS12 reanalysis (*74*). The proportion of larvae that are alive and competent varies through time (83), so we compute the potential connectivity matrix as

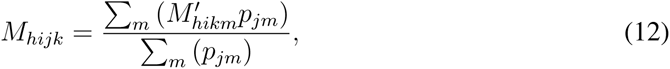

where *M_hikm_′* is the raw potential connectivity matrix, and *p_jm_* is the proportion of larvae alive and competent at time *τ_m_* = [2, 4 *· · ·,* 60] days after spawning. In reality, *p_jm_* will vary by species, as well as environmental parameters such as temperature (*75*), but we represent *p_jm_* for the fast-growing and slow-growing groups using parameters for *Acropora millepora* and *Goniastrea retiformis* respectively, to account for the generally lower dispersal distances (and presence of brooding reproductive strategies) within the slow-growing group (*76*). Given the broad number of species and global scope considered in CERES, and the general dominance of high-frequency variability over seasonality in potential connectivity (*77*), we randomly sample potential connectivity matrices across all months (this randomness does not introduce significant uncertainty, see figure S37) and assume that potential connectivity (mean state and variability) will not change significantly in the future. Although surface currents are expected to change over the coming century, these changes are small compared to the mean state and variability due to eddies, so this assumption is reasonable (*3, 78*).

### CERES evolutionary dynamics

We allow only one trait to evolve, namely the (mean) coral thermal optimum *z_ijk_* (with the subscripts *i*, *j* and *k* again respectively referring to subpopulations, coral groups, and time steps). In reality, other traits may also change through time. However, we focus here on the thermal optimum, as it is changes in the thermal environment that are expected to drive most future decline in coral cover (*7*). The actual thermal optimum trait value for individual coral colonies is distributed around this mean, with additive genetic variance*V_j_* (°C^2^). We set *V_j_* = 0.05°C^2^, the product of the narrow-sense heritability of thermal tolerance in corals, *h*^2^ *∼* 0.3 (*79*), and the phenotypic variance, *V_P_ ∼* 0.15°C^2^ (*43, 44*). In CERES, *V_P_* is held fixed in space and time, based on observed stability in coral genetic variance in response to major disturbances (*80*). We assume that *z_ijk_* can change through time due to (i) stabilizing selection, and (ii) the introduction of genotypes through immigration.

During months without spawning, *z_ijk_* only changes through stabilizing selection:

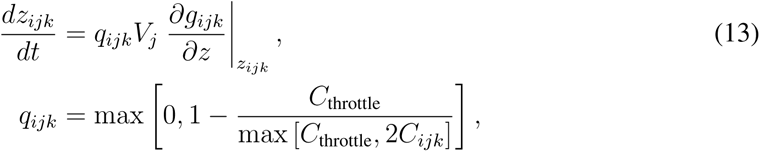

Here, *C*_throttle_ is a coral cover threshold below which selection is throttled (*22*), simulating bottlenecks/founder effects, which we arbitrarily set to 10*^−^*^3^. Along with *C*_min_, we are not aware of any empirical constraints for these parameters, so the chosen values are arbitrary. However, the model is insensitive to these parameters over a large range of values (figures S38-39), with results only changing noticeably when *C*_throttle_ is very large (greater than 1% coral cover).

During time-steps with spawning events, the mean trait value *z_ijk_* can also change through mechanism (ii), immigration. We assume that the mean trait value post-immigration is a simple admixture of the pre-immigration mean trait value, and the mean trait value across new immigrants, weighted by coral cover (*22*), i.e.

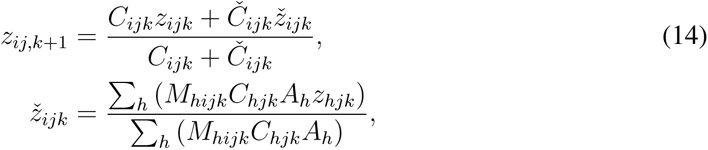

Here, *΂_ijk_* is the mean trait value amongst immigrant larvae, and *Č_ijk_* is the change in coral cover due to immigration, i.e., equation 11 minus *C_ijk_*.

## Acknowledgements

We are grateful to the Global Coral Reef Monitoring Network for sharing their coral cover data with us. We also thank the Marine Ecological Theory lab at the Hawai’i Institute of Marine Biology, Ariel Greiner, and Renato Morais, for their thoughts and comments on this research.

## Author Contributions

NSVV and LCM conceived the research. NSVV designed and conducted analyses, with input from all authors, and supervision from LCM. NSVV wrote the original draft, and all authors edited the manuscript.

## Data and materials availability

All scripts (including CERES) required to reproduce analyses and figures in the manuscript and supplementary materials can be found at 10.5281/zenodo.12560847. The full model output and forcing files will be made available in an institutional repository prior to publication.

