## Supplementary materials for "Anthropogenic climate change will likely outpace coral range expansion"

### This PDF file includes:

- Supplementary Text
- Figs. S1 to S39
- Movies S1 to S2

### Other Supplementary Materials for this manuscript include the following:

- Movies S1 to S2

### Model performance outside of the NW Atlantic and Indo-Pacific

Although coral reefs exist along eastern ocean boundaries, they are found in low abundance and represent less than 1% of global coral cover (*29*) . Eastern boundaries are upwelling regions, and are therefore associated with lower surface temperature, pH, and visibility. However, as demonstrated by CERES, these variables are not sufficiently deleterious in most upwelling regions to explain the lack of reefs. Indeed, coral reefs exist elsewhere in much more extreme thermal and turbidity conditions (*38*) . Upwelling also results in high nitrate and phosphate concentrations, which can increase competitive pressure on corals by macroalgae (*54*). These competitive interactions are poorly understood so, to avoid over-tuning this process-based model, we did not attempt to model direct or indirect effects of nutrient concentrations on coral growth rate. Given the strong correspondence between surface nutrient concentrations and coral cover overestimation in CERES, this is a possible explanation for this failure of the model to reproduce realistic coral reefs in the East Tropical Pacific, the Equatorial Pacific, and West Africa, although we note that the correlation between reef presence and nutrient concentrations is generally poor (*38*).

Nutrients cannot, however, explain the relatively low coral cover in the Southwest Atlantic, principally Brazil. Here, salinity appears to be a first-order control on the distribution of reef-building corals (*81*) . Apart from near river mouths, salinity along the coast of Brazil is generally within the range of conditions that are conducive to reef growth elsewhere (*38*). However, coral communities in the Southwest Atlantic are highly distinct (*57*), and it is therefore unsurprising that they may have a different sensitivity to their physical and chemical environment than other coral reef systems. The distinctiveness of coral communities in the marginal environments of the East Tropical Pacific and Equatorial Atlantic (*38*) suggests that, for optimal performance, model parameters should be tuned to different biogeographic realms. Again, for the sake of parsimony, we avoided this approach in the present study, but future models may wish to explore this possibility to improve range expansion projections outside of the NW Atlantic and Indo-Pacific.

Finally, although we incorporate the effect of sediment load on light availability in our simulations, we do not include the direct effects of sedimentation on smothering corals and inhibiting settlement (*6,82*). There is a high degree of overlap between regions of high sediment thickness (*83*) (a proxy for terrestrial sediment input) and coral reef absence, so it is possible that the direct effect of sedimentation could also explain the incorrect predictions by CERES of coral reef presence in some regions.

### Dependence of population growth rate on coral colony size structure

Following the same notation as used in the main text. Let site $i$ have $l$ colonies of coral group $j$ at time $k$. Coral colonies grow asexually through linear extension, which do not depend on the size of the coral colony (*84*). Assuming each colony occupies the space of a hemisphere with radius $r_{ijkl}$, growing at a site-specific linear extension rate $\varepsilon_{ijk}$,

$$\frac{d}{dt}r_{ijl}(t) = \varepsilon_{ijk}.$$

Since the footprint of each colony is given by $a_{ijl}(t) = \pi r_{ijl}(t)^2$, we substitute $r_{ijl}(t)$ for $a_{ijl}(t)$, giving

$$\frac{d}{dt}a_{ijl} = 2\sqrt{\pi}\varepsilon_{ijk}a_{ijl}(t)^{\frac{1}{2}}.$$

Using the fact that $C_{ij}(t) = \frac{1}{A_i}\sum_l a_{ijl}(t)^{\frac{1}{2}}$:

$$\frac{d}{dt}C_{ij} = \frac{1}{A_i}2\sqrt{\pi}\varepsilon_{ijk}\sum_l a_{ijl}(t)^{\frac{1}{2}} \tag{1}$$

We now assume that the distribution of coral colony areal footprint is log-normally distributed, which is the case for most coral populations (*25*), i.e. $a_{ijkl}$ can be modelled as a random log-normally distributed variable with parameters $\mu_{ijk}$ and $\sigma_{ijk}$. In this case, assuming that $n_l$ is large enough to approximate the distribution of $a_{ijl}$ across $l$ as continuous:

$$\begin{aligned}\frac{1}{n_l}\sum_l a_{ijl}(t)^{\frac{1}{2}} &\approx \int_0^\infty a^{\frac{1}{2}} f_A(a)\, da,\\ &\approx c_1 \exp\left(\frac{\mu_{ijk}}{2} + \frac{\sigma_{ijk}^2}{8}\right),\\ \sum_l a_{ijl}(t)^{\frac{1}{2}} &\approx n_l c_1 \exp\left(\frac{\mu_{ijk}}{2} + \frac{\sigma_{ijk}^2}{8}\right), \qquad (2)\end{aligned}$$

where the right-hand side of this equation is the 0.5-th arithmetic moment of the log-normal distribution, and $c_1 = 1$ m (a unit dimension to associate dimensions with the probability distribution). Similarly, from the 1-st arithmetic moment of the log-normal distribution (the mean), we have

$$\begin{aligned} \frac{1}{n_l}\sum_l a_{ijl}(t) &\approx c_2 \exp\left(\mu_{ijk} + \frac{\sigma_{ijk}^2}{2}\right), \\ \sum_l a_{ijl}(t) &\approx n_l c_2 \exp\left(\mu_{ijk} + \frac{\sigma_{ijk}^2}{2}\right), \\ C_{ij}(t)A_i &\approx n_l c_2 \exp\left(\mu_{ijk} + \frac{\sigma_{ijk}^2}{2}\right), \end{aligned} \tag{3}$$

where the last line uses the fact that the total area covered by coral colonies ($\sum_l a_{ijl}$) is equal to the fractional coral cover $C_{ij}$ multiplied by the habitable area $A_i$, and $c_2 = 1$ m$^2$ (again, a unit dimension required to give correct dimensions to the probability distribution). Substituting $n_l$ in equation 3 into equation 2, we find

$$\sum_l a_{ijl}(t)^{\frac{1}{2}} \approx C_{ij}(t)A_i s_0 \exp\left(-\frac{\mu_{ijk}}{2} - \frac{3\sigma_{ijk}^2}{8}\right), \tag{4}$$

where $c_0 = c_1/c_2 = 1\text{m}^{-1}$. Finally, substituting equation 4 into equation 1, we obtain the result

$$\begin{aligned} \frac{d}{dt}C_{ij} &\approx s_{ijk}\varepsilon_{ijk}C_{ij}(t) \\ s_{ijk} &= 2\sqrt{\pi}c_0 \exp\left(-\frac{1}{2}\mu_{ijk} - \frac{3}{8}\sigma_{ijk}^2\right) \end{aligned}$$

Therefore, to convert a linear extension rate into a population growth rate, we multiply it by factor $s_{ijk} = 2\sqrt{\pi}c_0 \exp\left(-\frac{1}{2}\mu_{ijk} - \frac{3}{8}\sigma_{ijk}^2\right)$. In the main text, we incorporate $s_{ijk}$ into the calculation of $g_{0,j}$ (i.e. the maximum population growth rate).

Since $s \propto \exp(-\mu)$, and $\mu = \ln A_{\text{med}}$ (where $A_{\text{med}}$ is the median coral colony area) for a log-normal distribution, we also find that $s \propto \frac{1}{\sqrt{A_{\text{med}}}}$, i.e. the population growth rate is inversely proportional to the square root of the median coral colony area.

### Dependency of mortality rate on temperature

The heat stress mortality term used in this model, is as follows:

$$\frac{dC}{dt} = -m_0 \frac{(T_k - Z_{H,k})^2}{2w_H^2} \delta_{T_k > Z_{h,k}}, \tag{5}$$

letting $Z_H = z + z_H$ for brevity. Under the influence of mortality due to heat stress alone, and assuming that $T > Z_H$ is constant within a time-step (as is the case in our model), the solution to equation 5 is:

$$C_{k+1} = C_k \exp\left(-\frac{(T_k - Z_{H,k})^2}{2w_H^2} m_0 \Delta t\right). \tag{6}$$

Rearranging equation 6, we find that the mortality $M_k$ over time-period $\Delta t$ is given by:

$$M_k = 1 - \frac{C_{k+1}}{C_k} = 1 - \exp\left(-\frac{(T_k - Z_{H,k})^2}{2w_H^2} m_0 \Delta t\right). \tag{7}$$

Many studies investigate coral mortality due to accumulated heat stress, measured in degree-heating weeks (DHW). A degree heating week is (roughly) the time-integral of the temperature anomaly above a thermal threshold. Following the above formulation, within a single time-step, we have the following approximate relationship when $T_k > Z_{H,k}$:

$$\text{DHW} = c_{dw} \left(T_k - Z_{H,k}\right) \Delta t, \tag{8}$$

where $c_{wy} = 52$ week y$^{-1}$ due to the different units of time used in our model (years) and DHW (weeks) - recall that the units of $m_0$ are y$^{-1}$. Substituting equation 8 into equation 7, we have

$$M_k = 1 - \exp\left(-\frac{\text{DHW}^2}{2c_{wy}^2 w_h^2 \Delta t} m_0\right) \tag{9}$$

In other words, for a constant time period of heat exposure, we predict the coral mortality $M_k$ to vary with $1 - \exp\left(k \times \text{DHW}^2\right)$. This fits well with available data (*43,85*), so we see equation 5 as a sensible parameterisation for mortality due to heat stress, although we acknowledge that this comparison is rough due to the assumptions made above.

Defining $\mathrm{DHW}_{50}$ as the accumulated heat stress required to cause 50% mortality and rearranging equation 9 to solve for $w_h^2$, we find

$$w_h = \sqrt{\frac{m_0}{2\ln(2)c_{wy}^2\Delta t}}\mathrm{DHW}_{50},$$
$$\approx \frac{\mathrm{DHW}_{50}}{17.7}, \tag{10}$$

where the last step takes $m_0 = 1$ y$^{-1}$ and $\Delta t = \frac{1}{12}$ y. equation 10 allows us to compute a value for $w_h^2$ in the model, using an empirical value of $\mathrm{DHW}_{50}$. Note that this derivation is equally valid for parameterising $w_c$ for cold stress, by replacing DHW with DCW (degree cooling weeks), and $T_k - Z_{h,k}$ with $Z_{c,k} - T_k$, $Z_c = z - z_c$.

### Computation of effective fecundity

The 'effective fecundity' $f_j$ as used in CERES can be interpreted as the proportional increase in coral cover due to the establishment of new coral colonies (or alternatively, the area of coral cover generated per unit area of existing coral cover), assuming 100% potential connectivity. Assuming hemispheric colony geometry, the number of coral polyps in a colony is $2\pi r_l^2/a_{0,j}$, where $r_l$ is the colony radius and $a_{0,j}$ is the surface area of a single polyp. The number of polyps per unit area footprint of the colony is therefore $2/a_{0,j}$ m$^{-2}$. If the number of eggs produced per spawning event per polyp is $f_{0,j}$ then the coral fecundity is $2f_{0,j}/a_{0,j}$ eggs m$^{-2}$. Finally, if the proportion of eggs that successfully transition to sexually mature corals is $r$, and the areal footprint of a newly established coral colony is $a_{1,j}$, then the new coral area generated per unit area of existing coral from a spawning event is

$$f_j = 2f_{0,j}r\frac{a_{1,j}}{a_{0,j}}.$$

Assuming that a newly established coral colony as the same size as a single colonial coral polyp, the equation simplifies to

$$f_j = 2f_{0,j}r.$$

This quantity is the *effective fecundity*. We use $f_{0,j}$ from (*23*). $r$ is in turn given by the following expression:

$$r = r_f \times r_t \times r_s \times r_j, \tag{11}$$

where $r_f$ is the proportion of eggs that are fertilised, $r_t$ is the proportion of larvae that survive transport, $r_s$ is the proportion of larvae arriving at a reef that successfully undergo recruitment, and $r_j$ is the proportion of recruits that survive to sexual maturity. There is considerable uncertainty (and variability in all of these parameters).

We set $r_f = 0.1$ based on in-situ measurements of fertilisation rates (*47*). For the computation of the potential connectivity matrix, we normalise potential connectivity based on the likelihood that a larva is alive and competent when it reaches another site, neglecting en-route settlement. $r_t$ therefore varies between 0 (if a virtual particle in `EZfate` never reaches another suitable site) and 1 (if a virtual particle in `EZfate` spends a full 60 days above suitable sites). We set $r_s = 0.1$, although acknowledge that this number is likely species- and substrate-dependent (*48*), and that a considerably lower figure was suggested by (*86*). Finally, we set $r_j = 0.02$, again acknowledging that this value will in reality be environment-dependent (*48*). This results in an estimate of $r = 2 \times 10^{-4}$.

### Computation of change in coral cover due to density-dependent settlement

We assume that the rate of change of coral cover $\mathbb{C}_{ik}(t)$ ($\mathbb{C}_{ik}(t) = \sum_j C_{ijk}(t)$) at a site during a spawning event is given by the following equation:

$$\frac{d\mathbb{C}_{ik}}{d\xi} = \mathbb{I}'_{ik}(\xi)\left(1 - \mathbb{C}_{ik}\right), \tag{12}$$

where $\xi = [0, 1]$ is the fraction of the spawning event, $\mathbb{I}'_{ik}(\xi)$ is the rate at which free space at $i$ is being taken up by new recruits, as a function of progress within a spawning event. Integrating equation 12 from the start of the spawning event to $\xi$:

$$\mathbb{C}_{i,k+1} = 1 - (1 - \mathbb{C}_{ik}) \exp(-\mathbb{I}_{ik}), \tag{13}$$

where $\mathbb{I}_{ik} = \int_0^1 \mathbb{I}'_{ik} \, d\xi$ is the total larval supply across a spawning event to $i$ (as a fraction of the habitable area at $i$). With this solution for $\mathbb{C}_{i,k+1}$, it is now possible to solve for $C_{ijk}(t)$ (i.e. for the individual coral groups), assuming larvae for all groups arrive at a similar time. This is represented by the following equation:

$$\frac{dC_{ijk}}{dt} = I'_{ijk}(\xi) \left(1 - \mathbb{C}_{ik}\right), \tag{14}$$

where $\mathbb{I}'_i(\xi) = \sum_j I_{ij}(\xi)$. This has the solution

$$C_{ij,k+1} = \frac{I_{ijk}}{\mathbb{I}_{ijk}} \left(1 - \mathbb{C}_{ik}\right) \left(1 - \exp(-\mathbb{I}_{ik})\right) + \mathbb{C}_{ik} \tag{15}$$

$I_{ijk}$ is the 'supply' of fractional coral $j$ cover at $i$ from incoming larvae. The 'supply' of areal coral $j$ cover arriving at $i$ from incoming larvae is equal to $f_j \sum_h (M_{hi} C_{hjk} A_h)$, where $M_{hi}$ is the potential connectivity $h \to i$, and $C_{hjk} A_h$ is the areal coral $j$ cover at $h$. Therefore,

$$I_{ijk} = \frac{f_j}{A_i} \sum_h \left(M_{hi} C_{hjk} A_h\right). \tag{16}$$

### Supplementary animations

#### Supplementary animation 1

Tropical coral assemblage population size (bubble size) and relative change versus 1850-1899 baseline (colours) under SSP2-4.5 from the GFDL-ESM4 model. Only Indo-Pacific and NW Atlantic subpopulations are plotted. The line chart on the right plots the global tropical coral cover, relative to the 1850-1899 baseline.

### Supplementary animation 2

Total coral population size (bubble size) and community composition (colours), following the same method as used in figure 6 in the Tropical coral assemblage population size (bubble size) and relative change versus 1850-1899 baseline (colours) under SSP2-4.5 from the GFDL-ESM4 model. Only Indo-Pacific and NW Atlantic subpopulations are plotted. The line chart on the right plots the global tropical coral cover, relative to the 1850-1899 baseline.

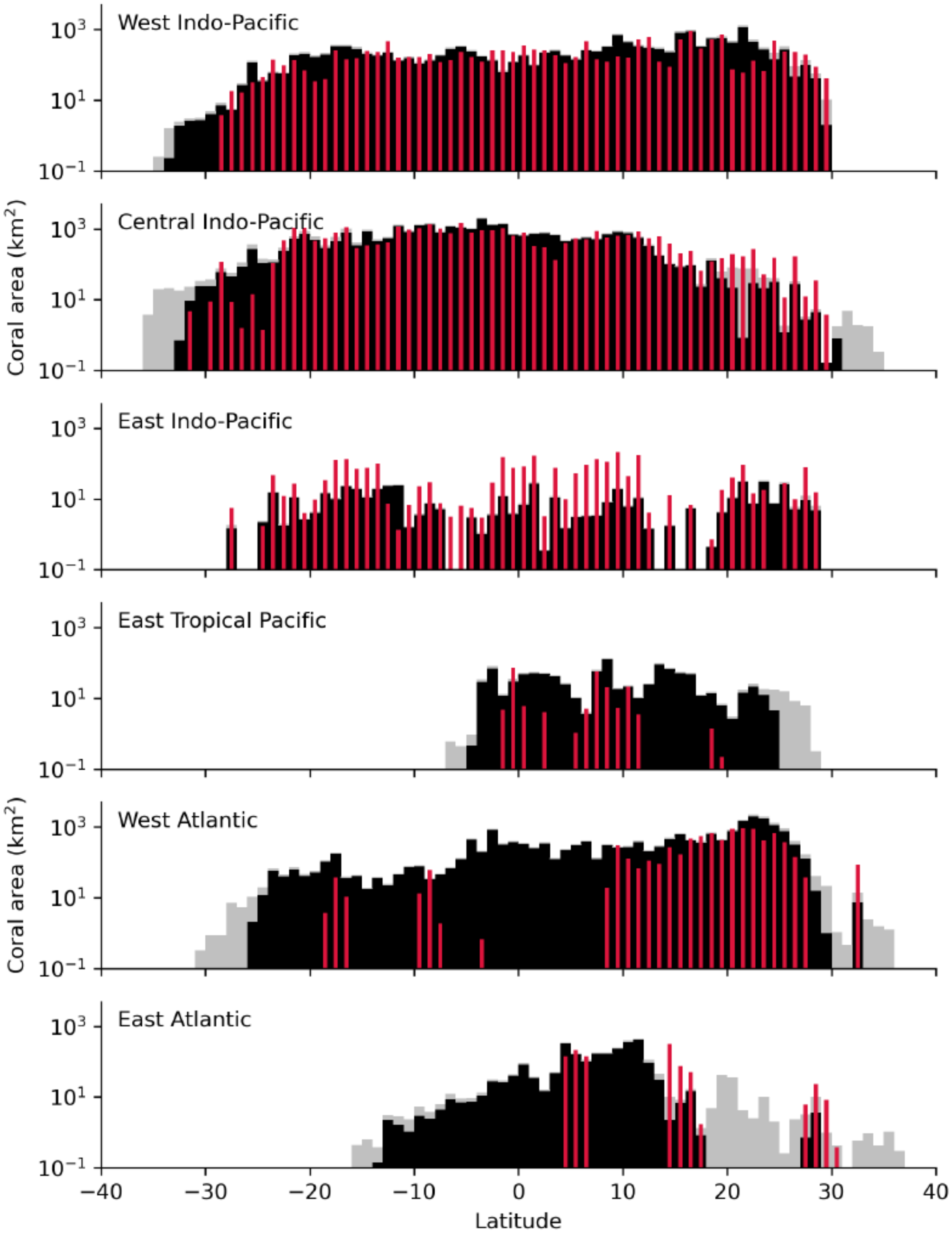


Fig. S1: Modelled tropical reef-building (black) and subtropical community-forming (grey) type coral area per latitude band for 2010-2019, compared against satellite-derived cover in coral reefs from the Allen Coral Atlas (*28*), for six major biogeographic realms (*87*). Here, we append subtropical and temperate provinces with corals to neighbouring tropical realms, e.g. appending all provinces in Japan and Australia to the Central Indo-Pacific realm.

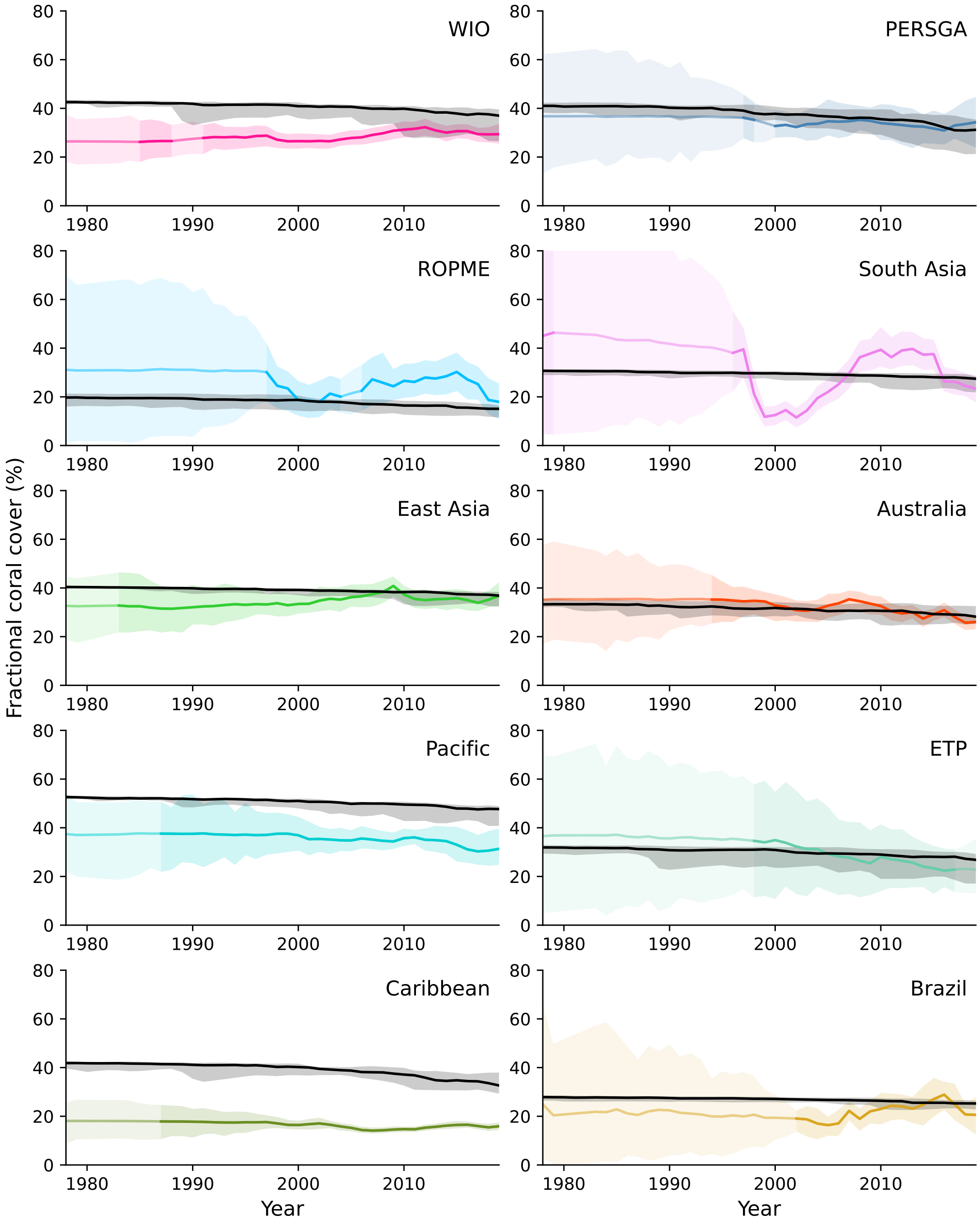


Fig. S2: Modelled mean coral cover across cells with $\geq 1\%$ coral cover (black) and estimated live hard coral cover from the Global Coral Reef Monitoring Network (colours) (*29*), for the 10 GCRMN monitoring regions. The shaded area represents the range across the model ensemble (black), and the 5-95% confidence interval for GCRMN-derived data (colours). Note that CMIP models are not assimilative, so would not be expected to reproduce global bleaching events in the correct years.

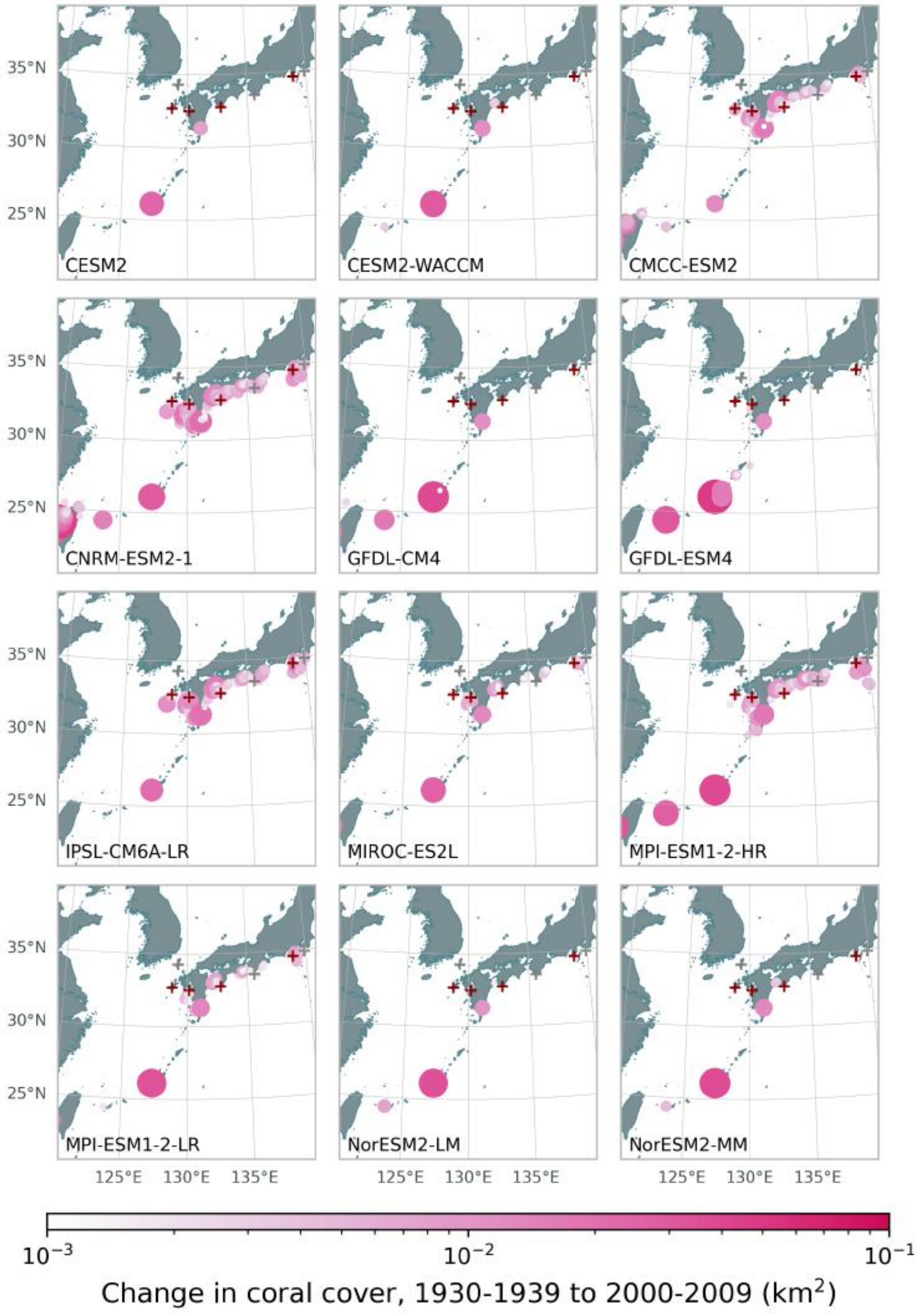


Fig. S3: Modelled change in total coral cover for all 12 ensemble members between 1930-1939 and 2000-2009. Sites where at least 1 coral species was considered to have expanded its range by (*15*) are marked with crosses; grey for one species only, and red for multiple.

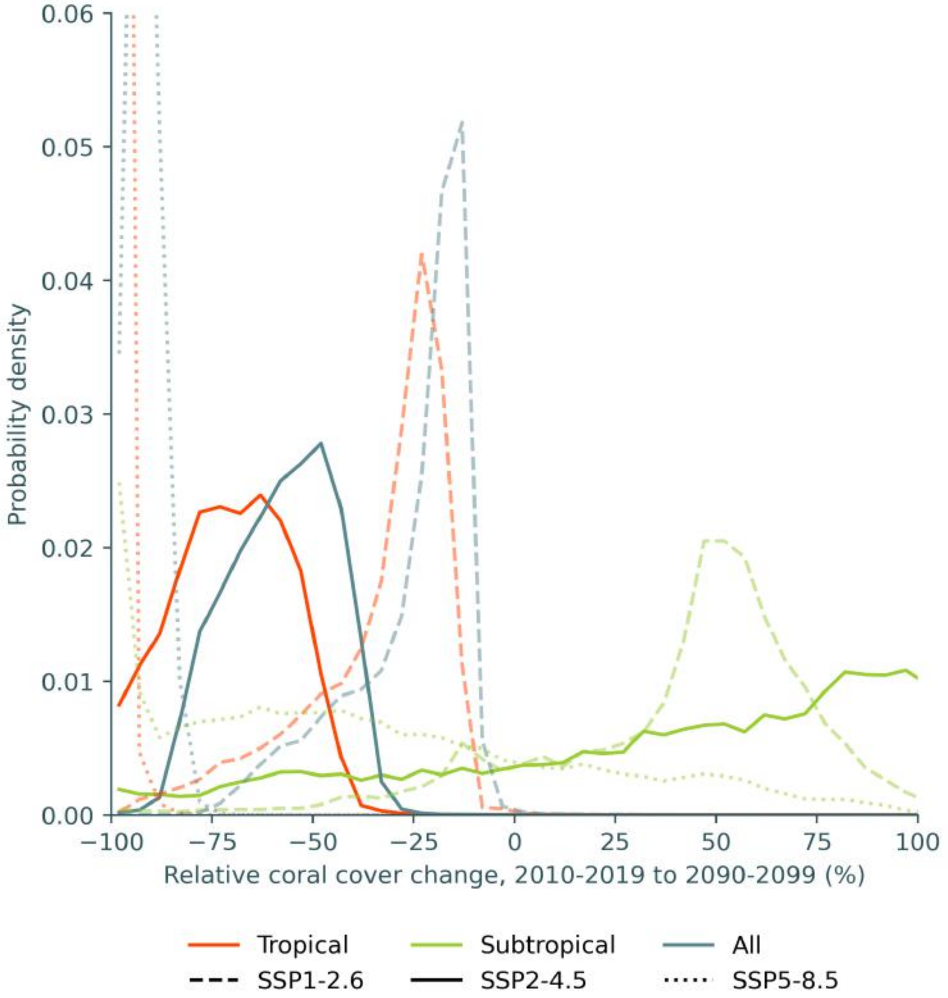


Fig. S4: Probability density for the (ensemble-mean) relative change in coral cover from 2010-2019 to 2090-2099, weighted by coral cover in 2010-2019.

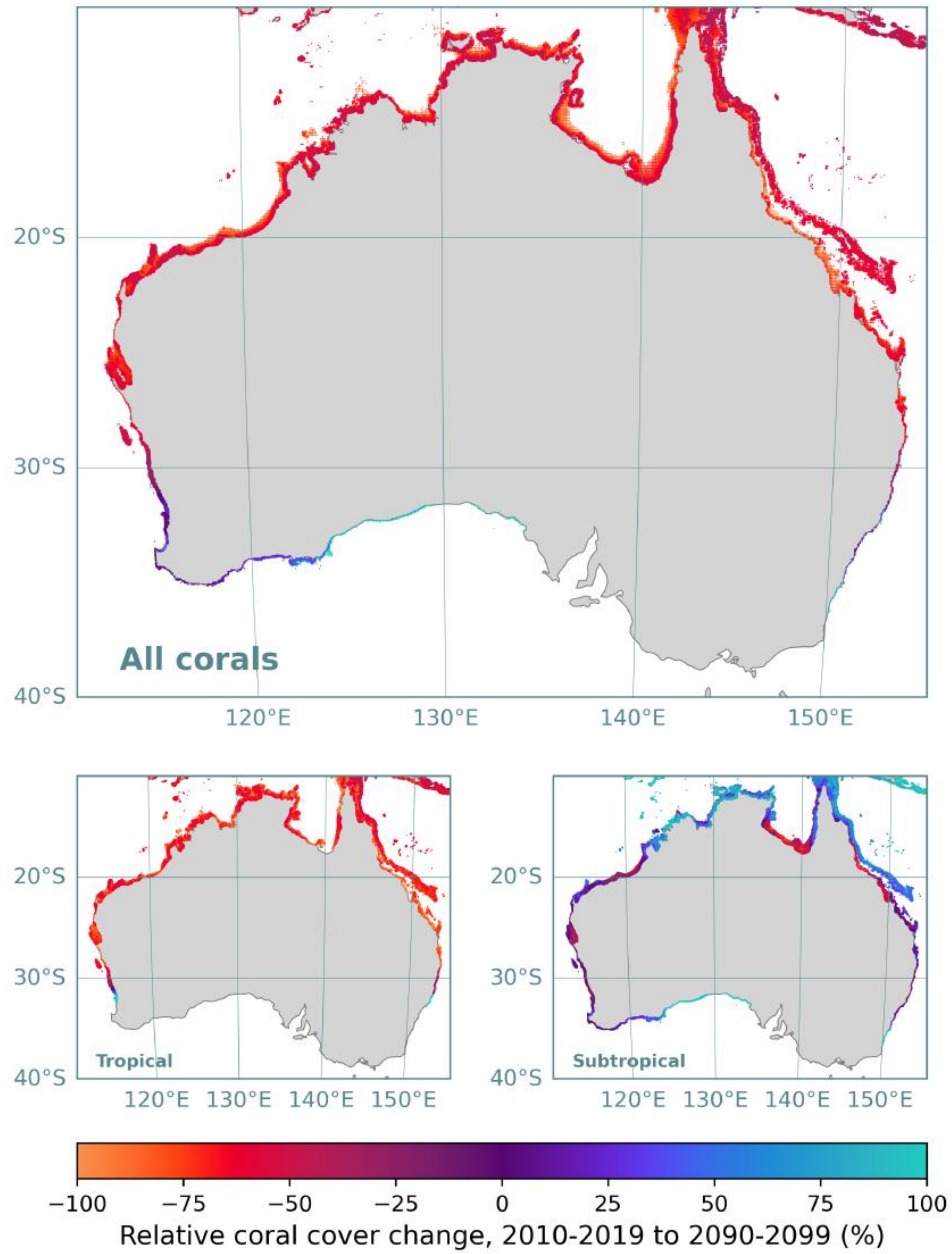


Fig. S5: Mean relative change in coral cover between 2010-2019 and 2090-2099 across all ensemble members in CERES around Australia for (a) all corals, (b) tropical corals, and (c) subtropical corals. Sites are scaled by the absolute coral cover between 2090-2099. For clarity, only sites with a final coral cover exceeding 0.001 $km^2$ are plotted.

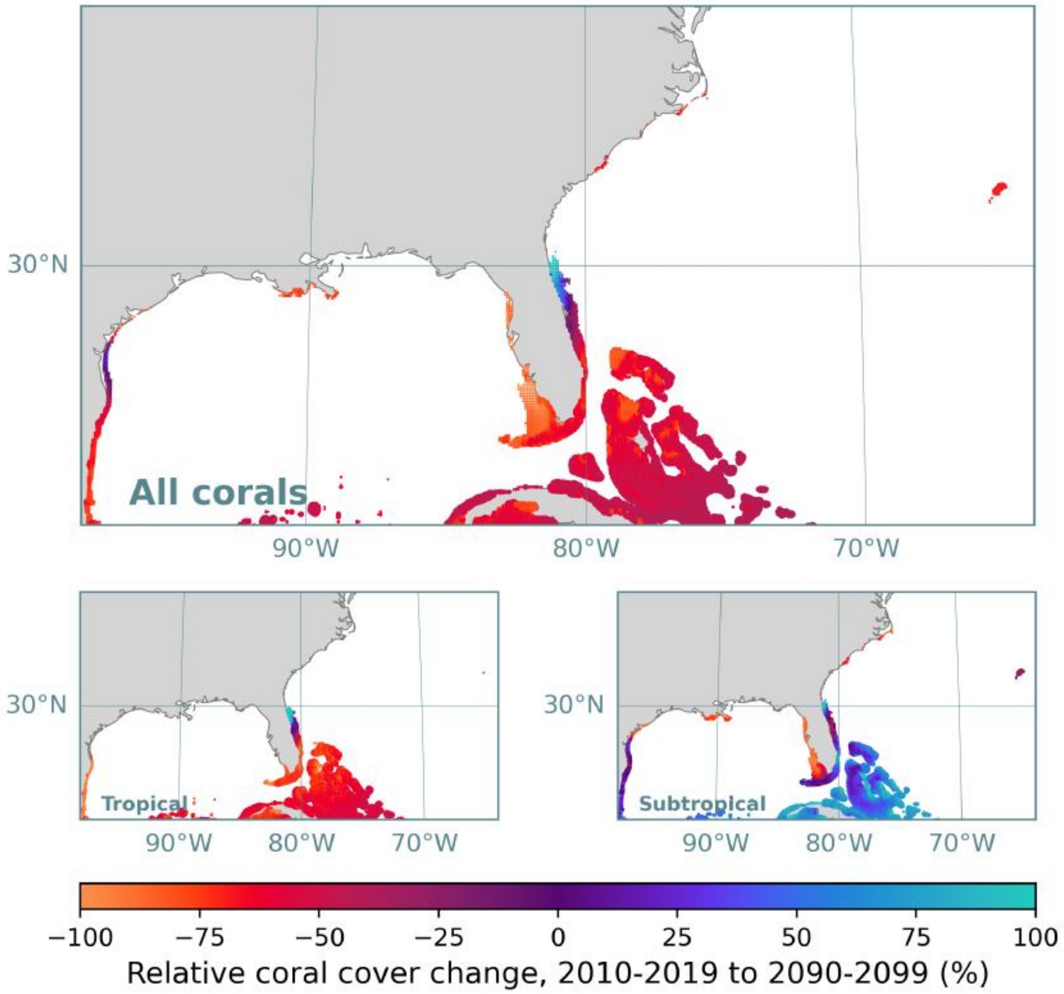


Fig. S6: Mean relative change in coral cover between 2010-2019 and 2090-2099 across all ensemble members in CERES around the NW Atlantic for (a) all corals, (b) reef assemblages, and (c) non-reef assemblages. Sites are scaled by the absolute coral cover between 2090-2099. For clarity, only sites with a final coral cover exceeding 0.001 $km^2$ are plotted.

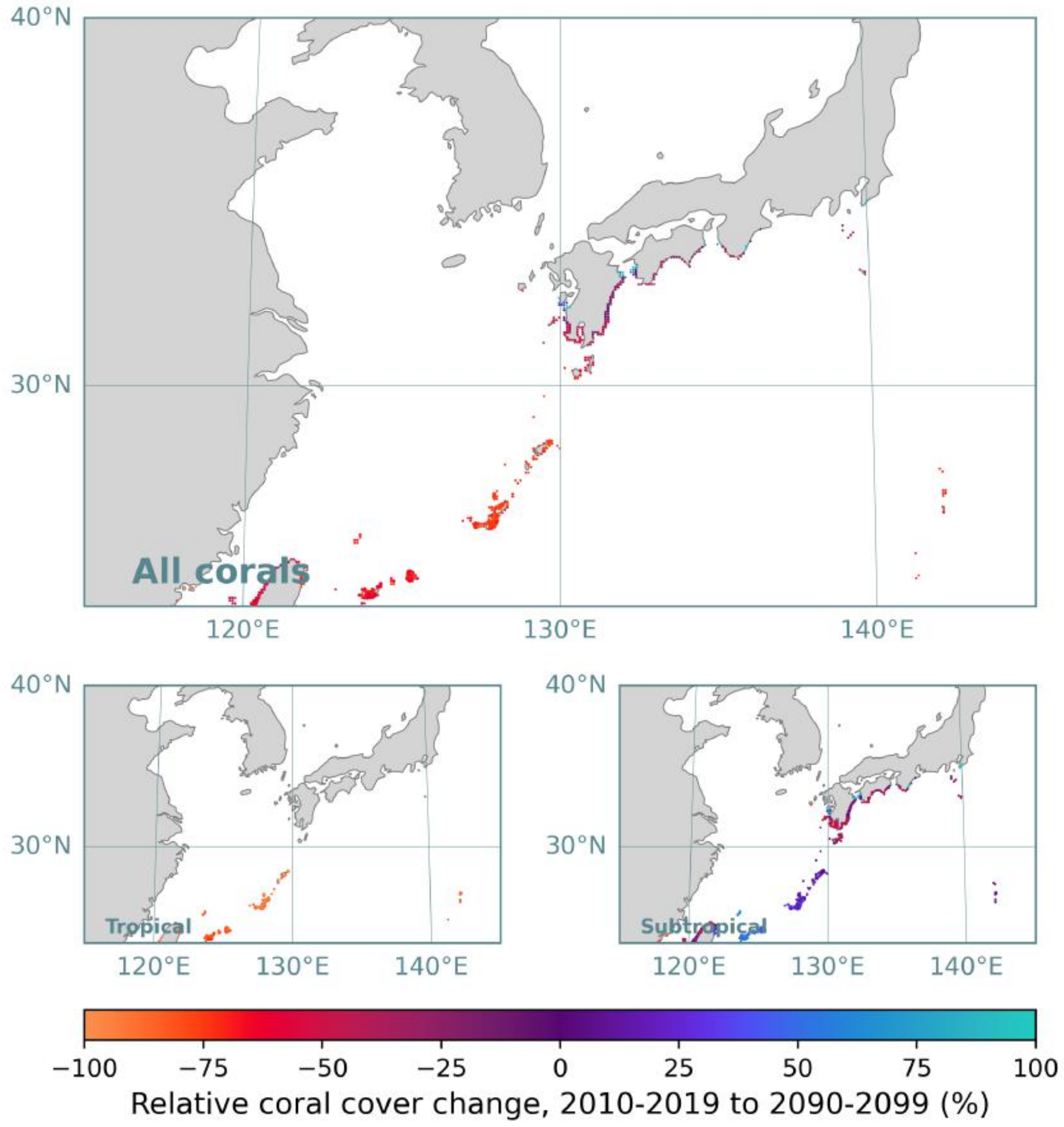


Fig. S7: Mean relative change in coral cover between 2010-2019 and 2090-2099 across all ensemble members in CERES around the NW Pacific for (a) all corals, (b) reef assemblages, and (c) non-reef assemblages. Sites are scaled by the absolute coral cover between 2090-2099. For clarity, only sites with a final coral cover exceeding 0.001 $km^2$ are plotted.

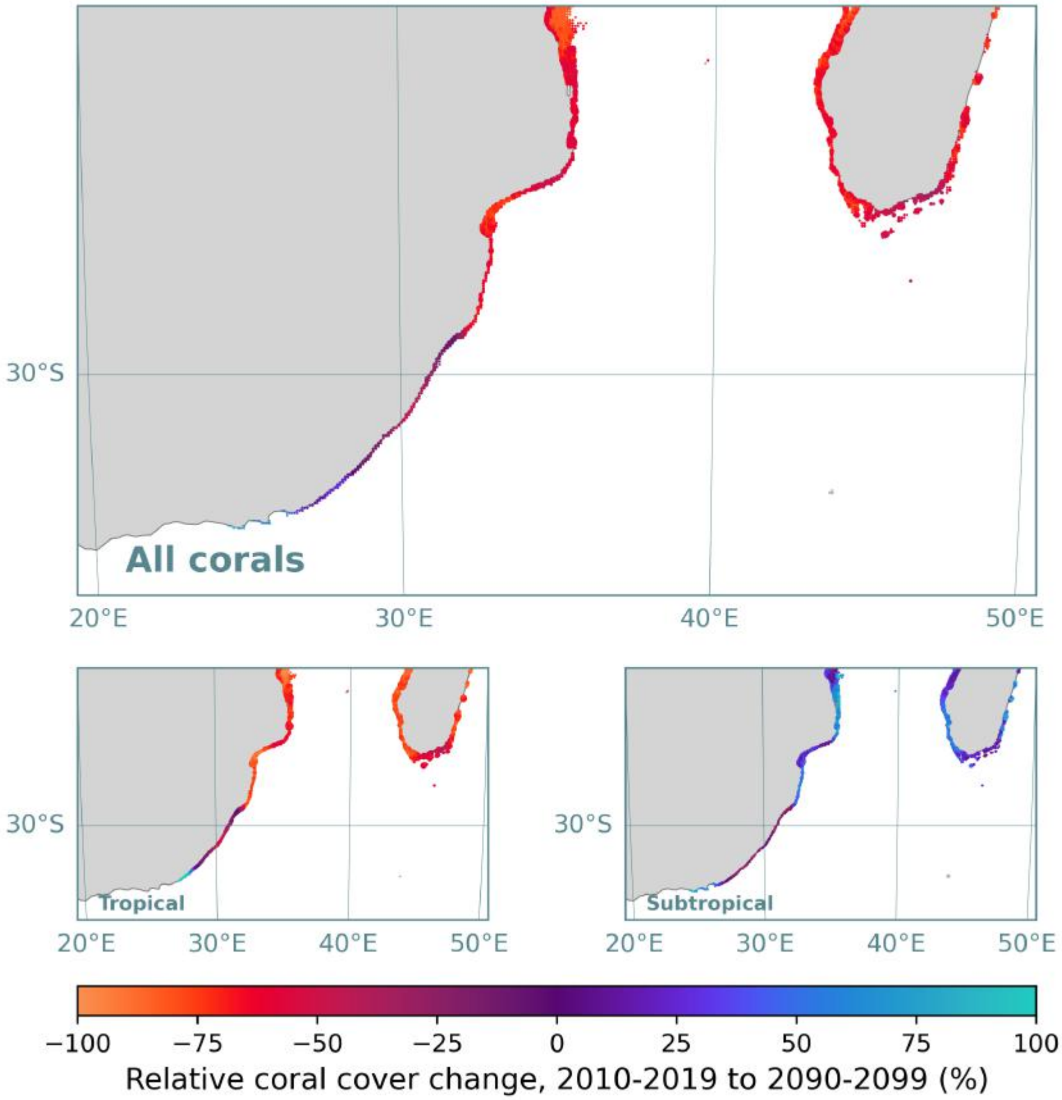


Fig. S8: Mean relative change in coral cover between 2010-2019 and 2090-2099 across all ensemble members in CERES around the SW Indian Ocean for (a) all corals, (b) reef assemblages, and (c) non-reef assemblages. Sites are scaled by the absolute coral cover between 2090-2099. For clarity, only sites with a final coral cover exceeding 0.001 $km^2$ are plotted.

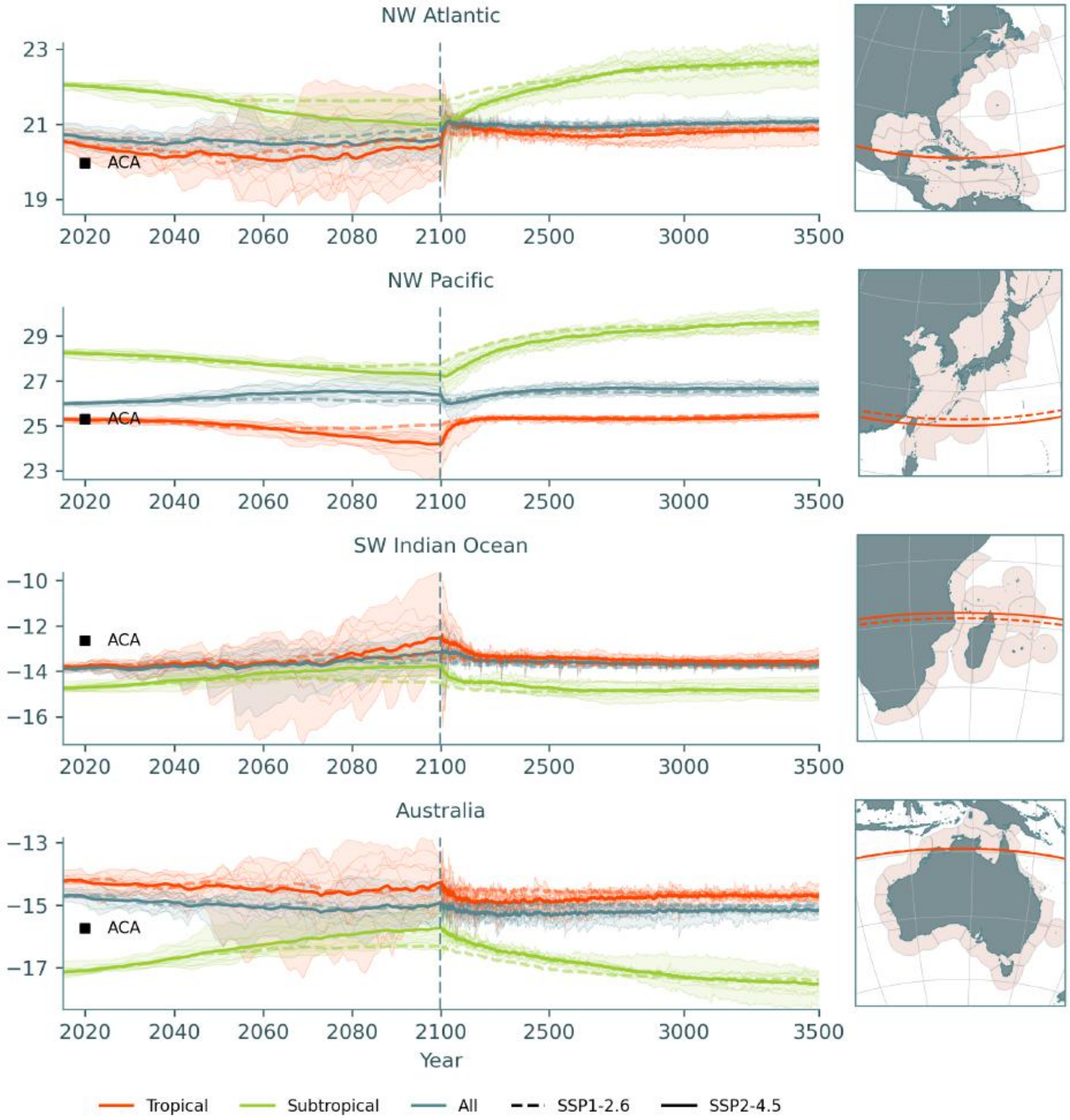


Fig. S9: Latitude of the coral cover centroid for SSP1-2.6 (dashed line) and SSP2-4.5 (solid line) for four high-latitude reef-systems (included ecoregions (*87*) are shaded in orange in the map panels). SSP5-8.5 is not shown, as the variations are too large to be represented on the same scale. The black squares represent the tropical coral centroid computed from the Allen Coral Atlas (*28*). Thin lines represent individual model trajectories for SSP1-2.6. Horizontal orange lines on the maps represent the tropical coral centroid latitude in 2010-2019 (dashed) and 2090-2099 (solid).

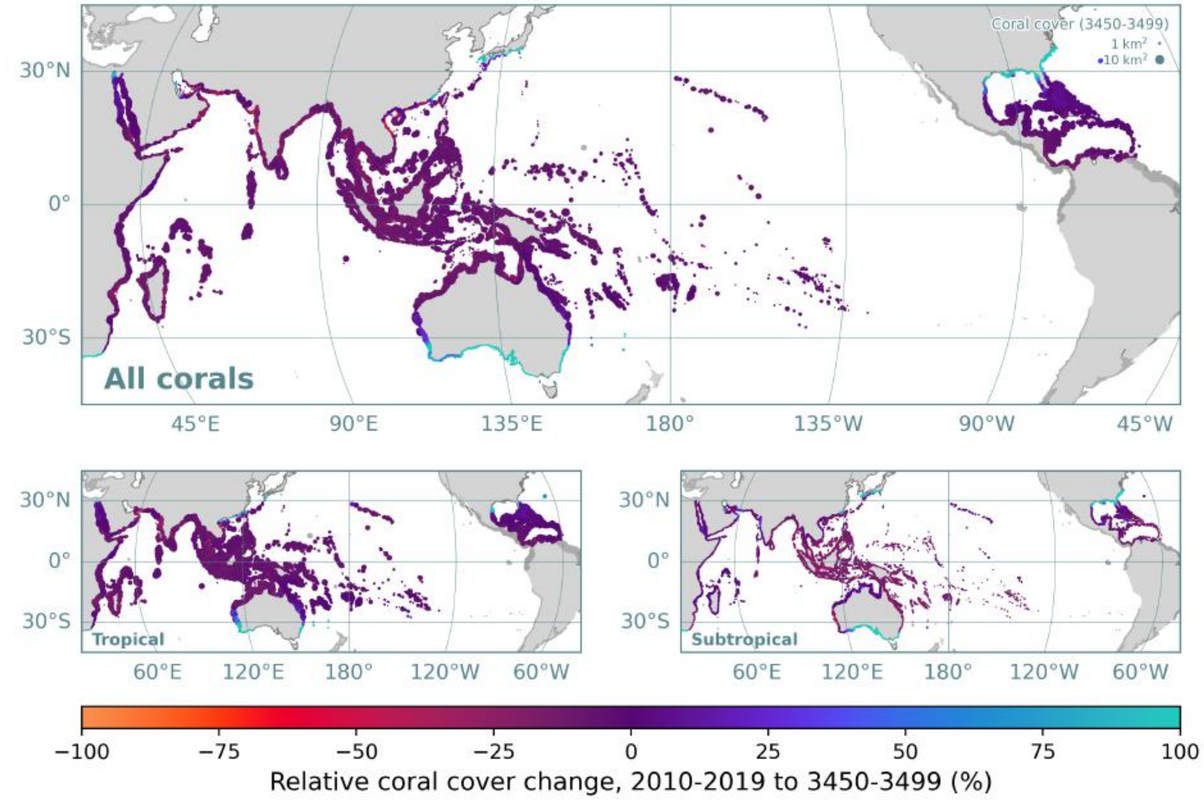


Fig. S10: Mean relative change in coral cover between 2010-2019 and 3450-3499 across all ensemble members in CERES for (a) all corals, (b) reef assemblages, and (c) non-reef assemblages. Sites are scaled by the absolute coral cover between 3450-3499. For clarity, only sites with a final coral cover exceeding 0.001 km$^2$ are plotted.

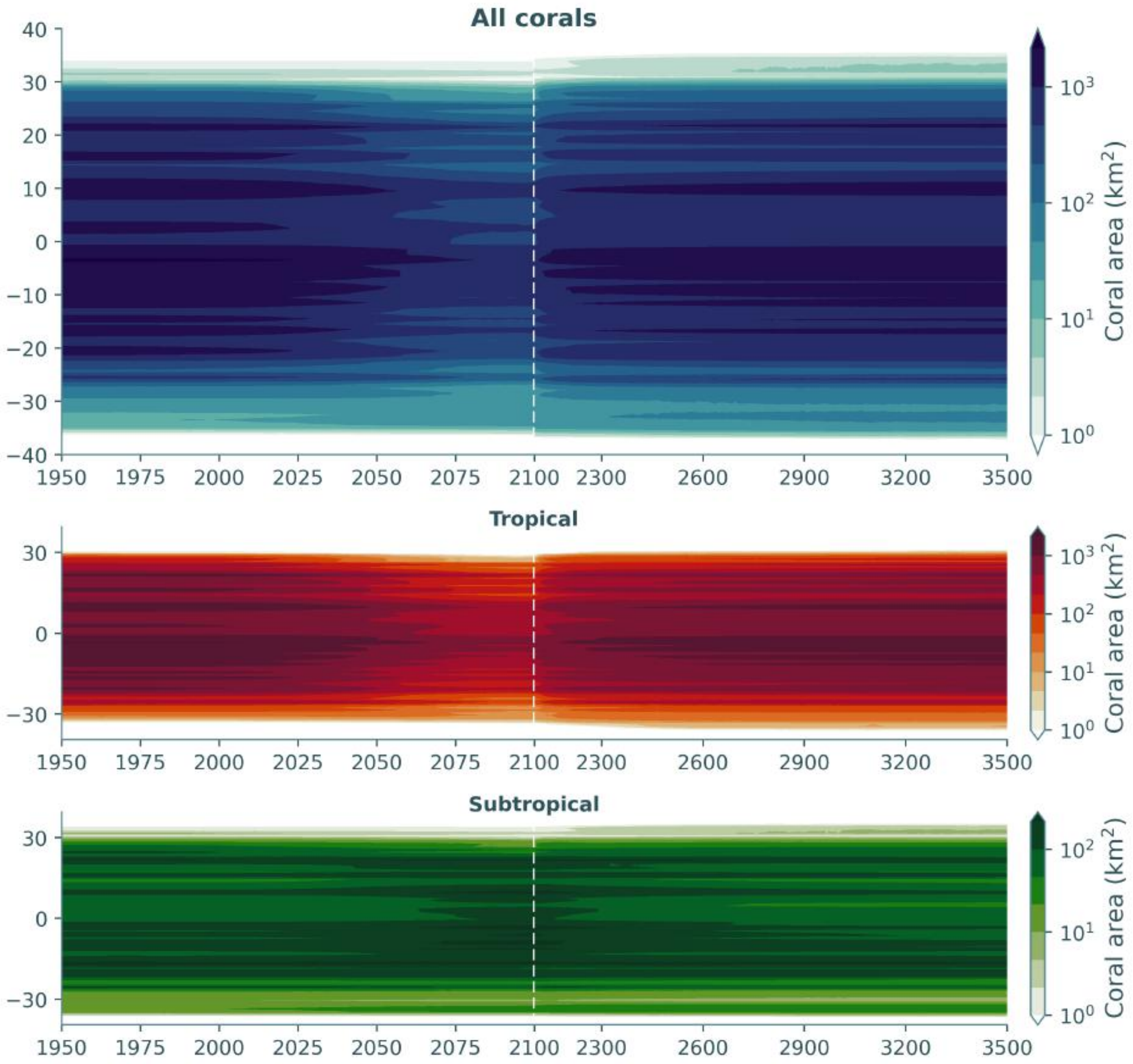


Fig. S11: Ensemble-mean coral cover per latitude band under SSP2-4.5 for all corals, tropical corals, and subtropical corals.

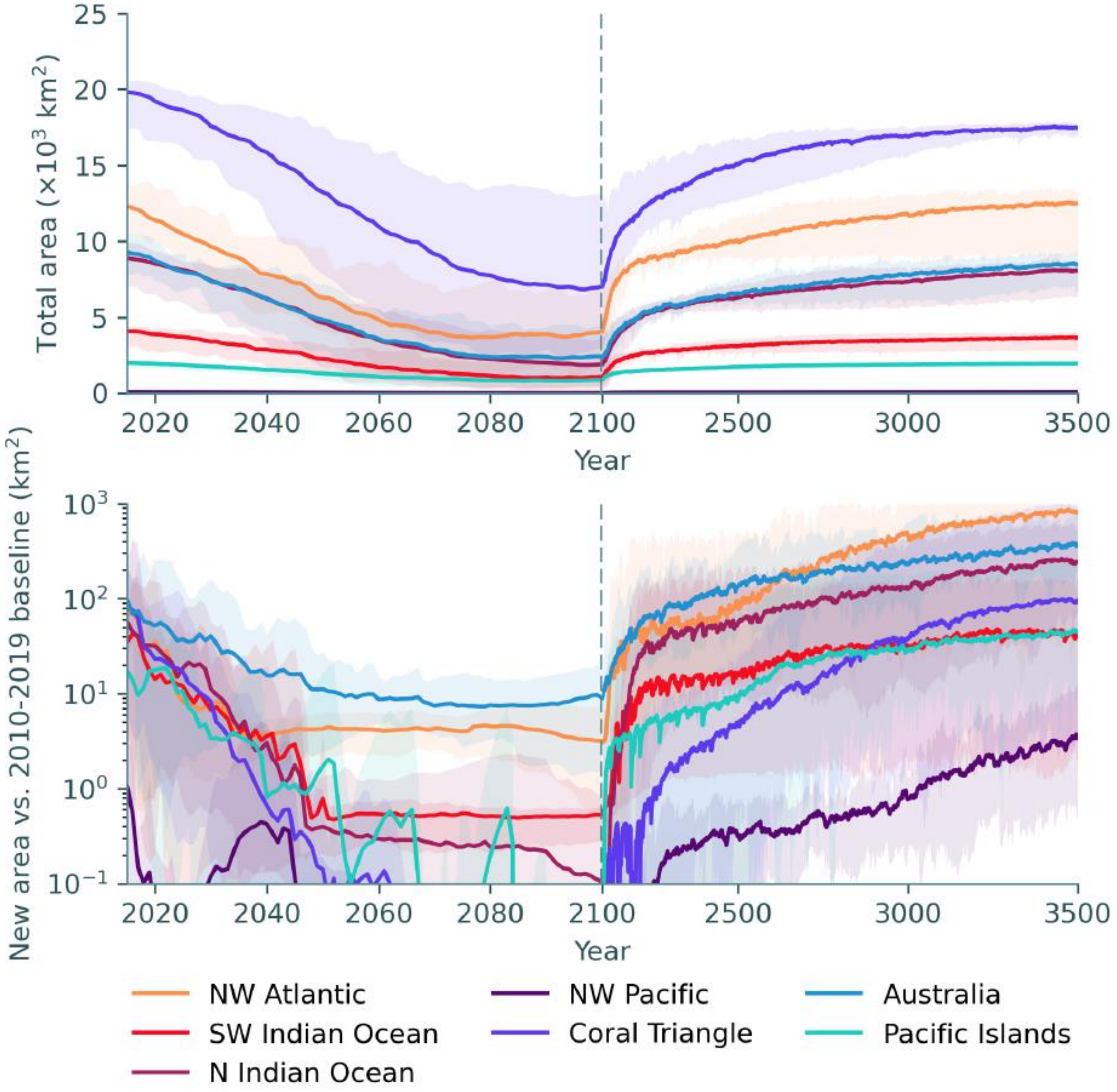


Fig. S12: (a) Total modelled area of reef assemblages *only* under SSP2-4.5 in seven major coral ecoprovinces. (b) Total new coral cover relative to the mean model state from 2010-2019 (the sum across all sites where coral cover increased). Note that the time axis scale changes in the year 2100.

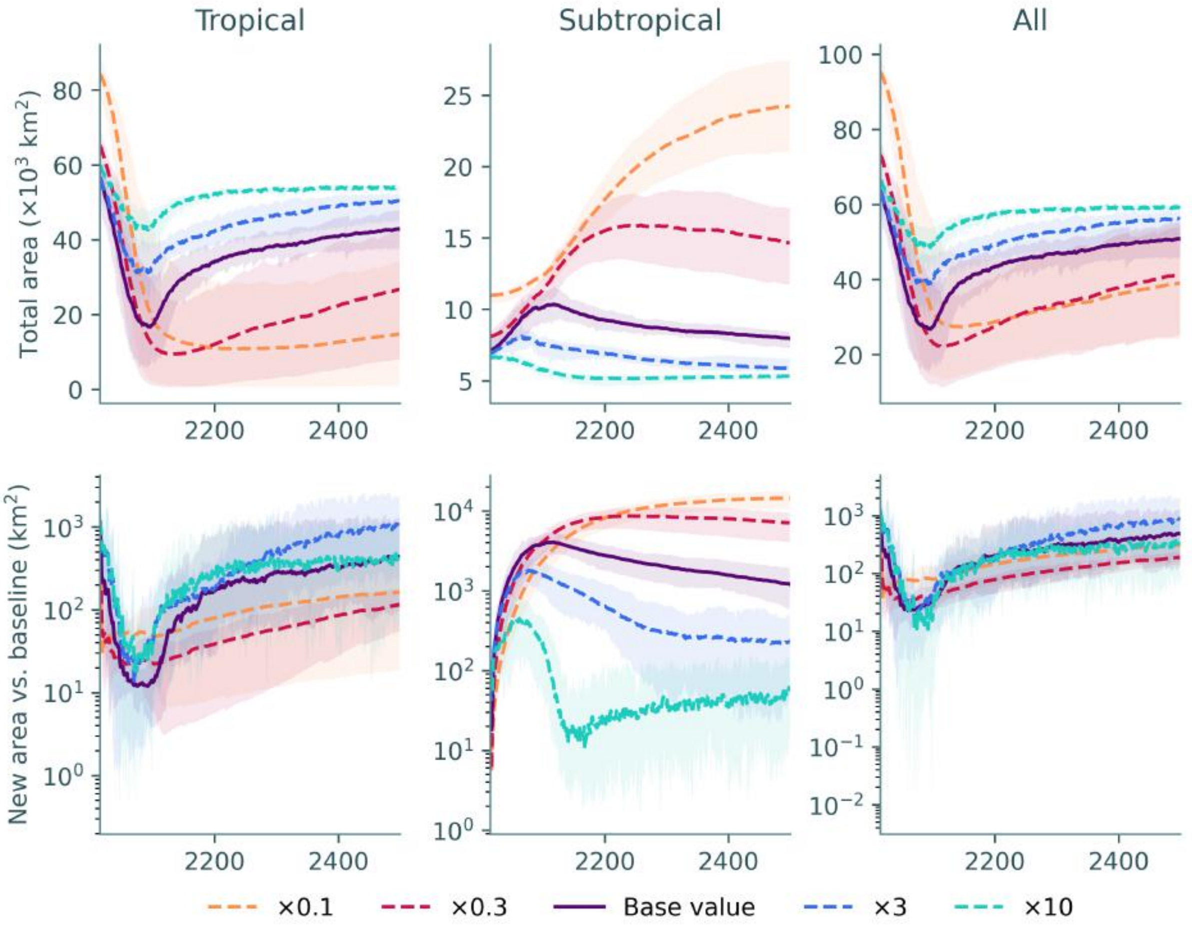


Fig. S13: Total modelled area (*top*) and new coral cover relative to the 2015-2019 baseline (*bottom*) of coral assemblages under SSP2-4.5, under different values of the (maximum) colony linear extension rate ($s_0$), from 2015 to 2500. Under low environmental stress, there is a higher equilibrium population size under lower growth rate, possibly due to the lower intensity of selection resulting in a more stable thermal optimum.

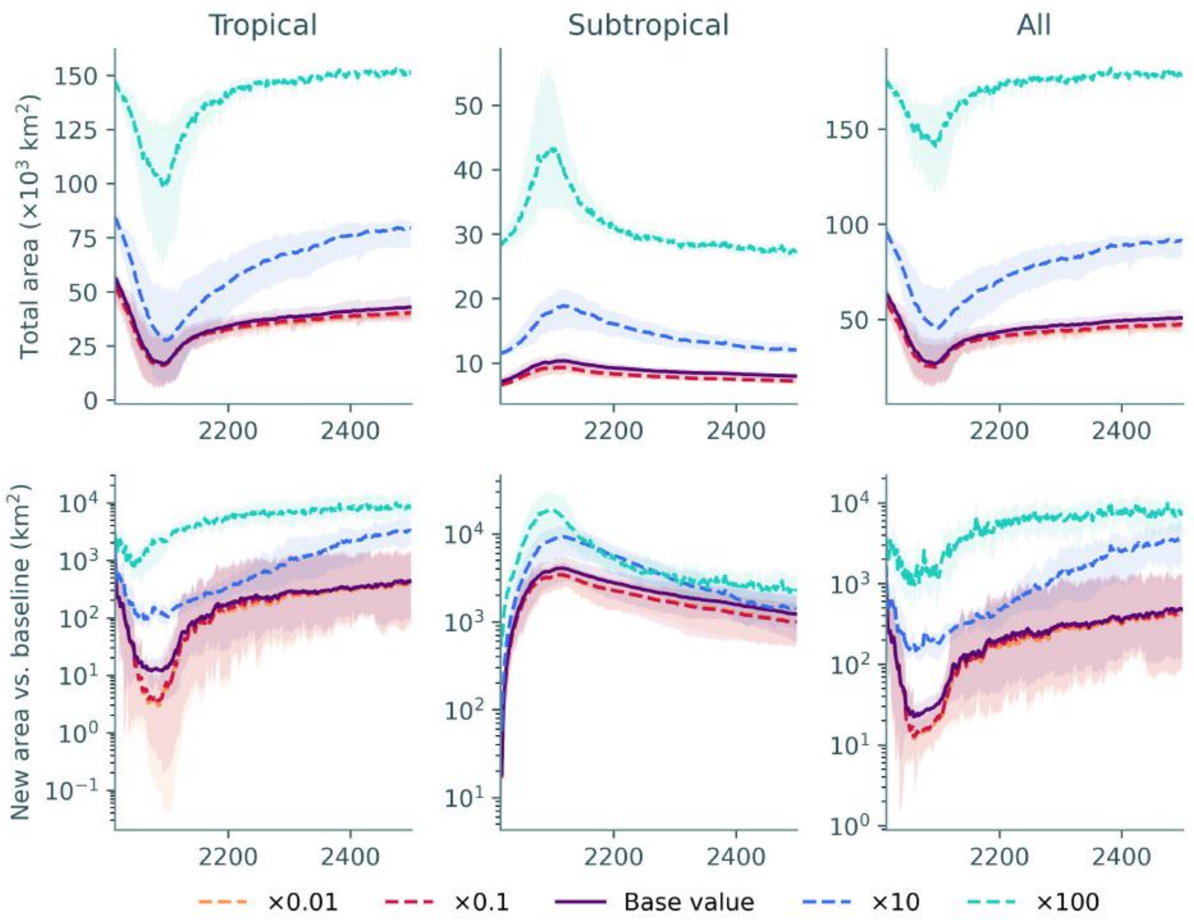


Fig. S14: Total modelled area (*top*) and new coral cover relative to the 2015-2019 baseline (*bottom*) of coral assemblages under SSP2-4.5, under different values of effective fecundity ($f$), from 2015 to 2500.

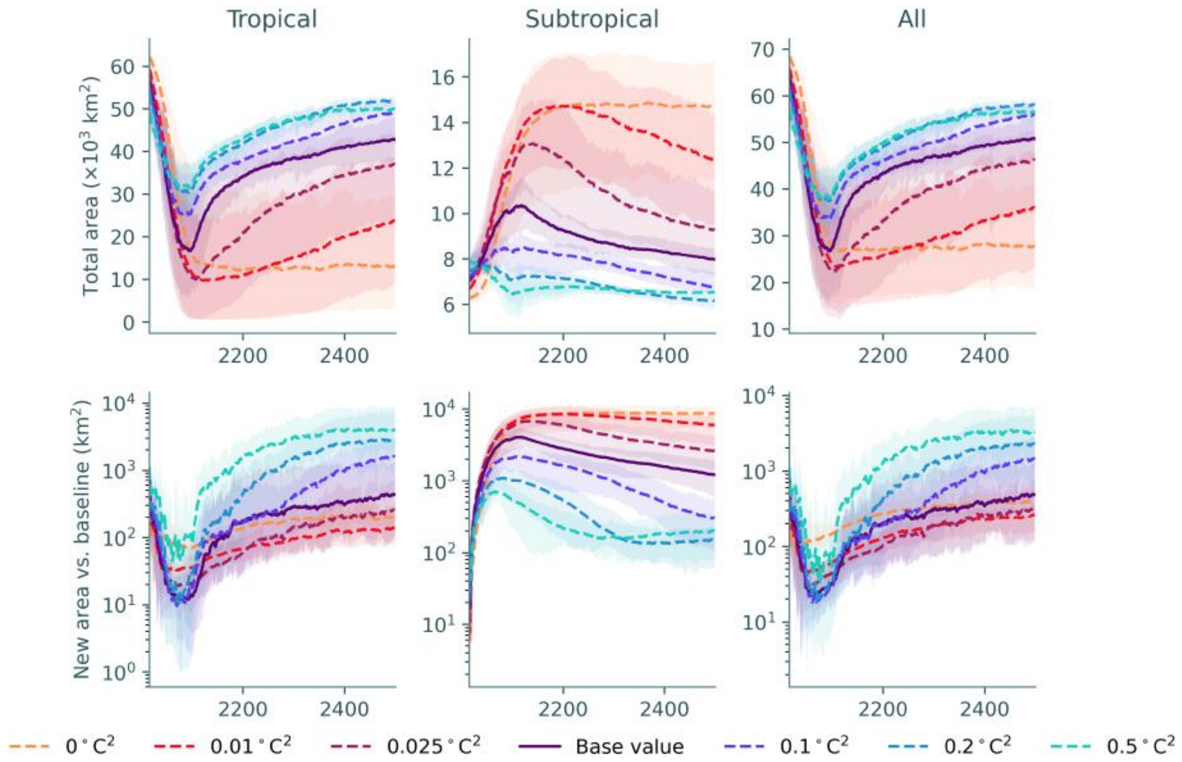


Fig. S15: Total modelled area (*top*) and new coral cover relative to the 2015-2019 baseline (*bottom*) of coral assemblages under SSP2-4.5, under different values of the additive genetic variance ($V$), from 2015 to 2500.

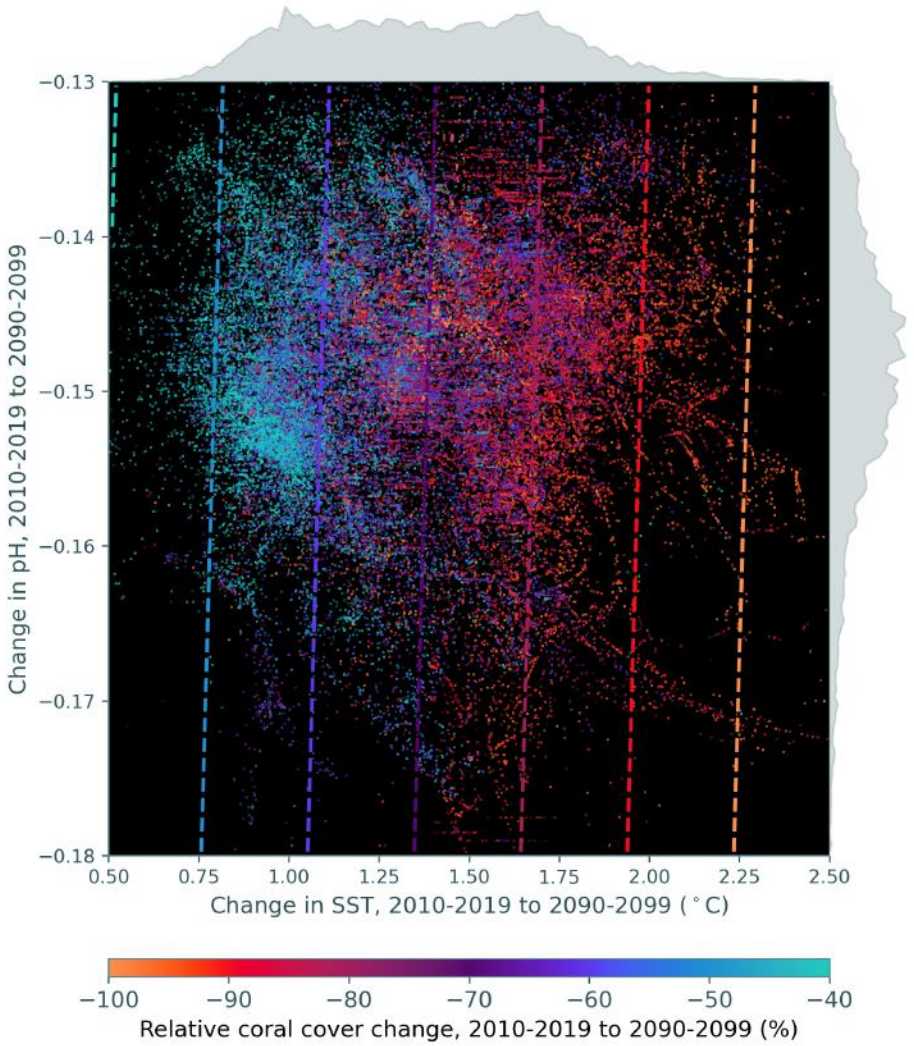


Fig. S16: Predicted (background) and simulated (points) relative change in tropical coral cover over the 21$^{st}$ century, as a function of the change in sea-surface temperature and pH. Only sites where the tropical coral cover originally exceeded 10% are included. The predicted change in coral cover was calculated from a linear mixed model, with random intercepts varying by ensemble member. Simulated data are plotted with random effects removed. Histograms show the distribution of predictor variables across reef sites.

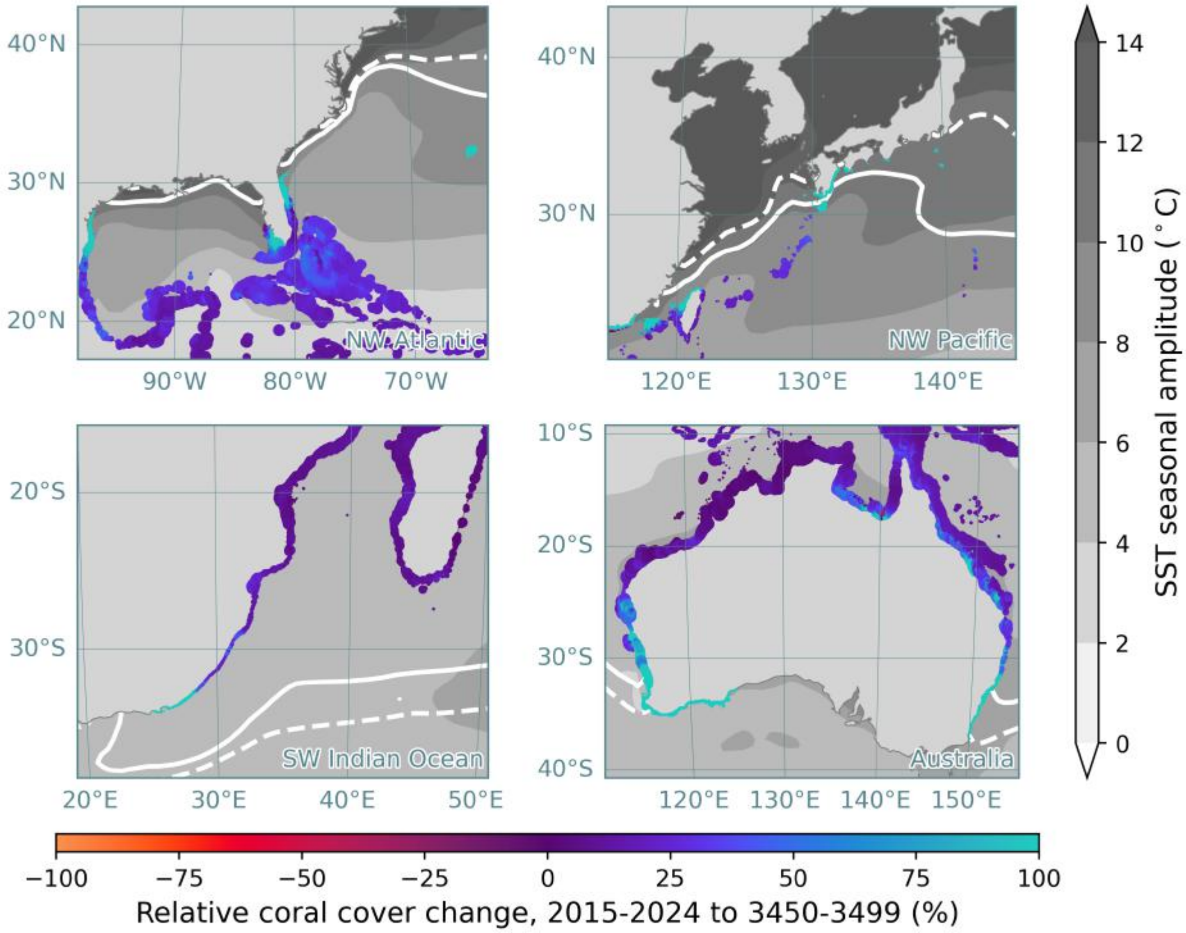


Fig. S17: Mean relative change in tropical coral cover between 2015-2024 and 3450-3499 under SSP2-4.5 *with pH fixed to pre-industrial levels* across all ensemble members in CERES for tropical coral assemblages. Sites are scaled by the absolute coral cover between 3450-3499. The ocean is shaded by the amplitude of the seasonal cycle of sea-surface temperature, and the solid and dotted white contours represent the annual minimum 18°C isotherm for the pre-industrial and post-2090 periods respectively. For clarity, only sites with a final coral cover exceeding 0.001 $km^2$ are plotted.

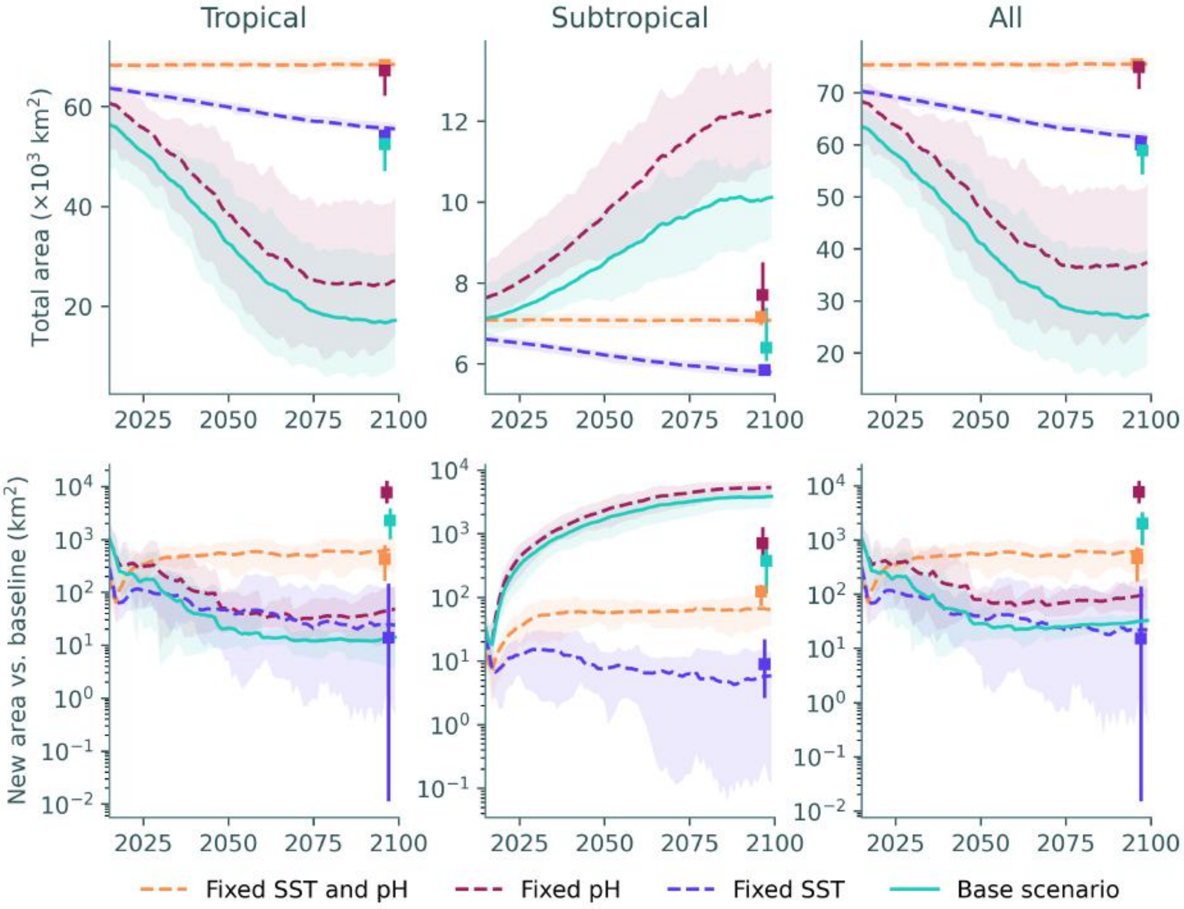


Fig. S18: Total modelled area (*top*) and new coral cover relative to the 2015-2019 baseline (*bottom*) of coral assemblages under SSP2-4.5, and alternative cases where pH and/or sea surface temperature are maintained in the pre-industrial scenario. Lower values represent a lower sensitivity to light intensity. Squares at the right of each axis represent the mean (long-term) value between 3450-3499, with vertical lines representing the range across the ensemble.

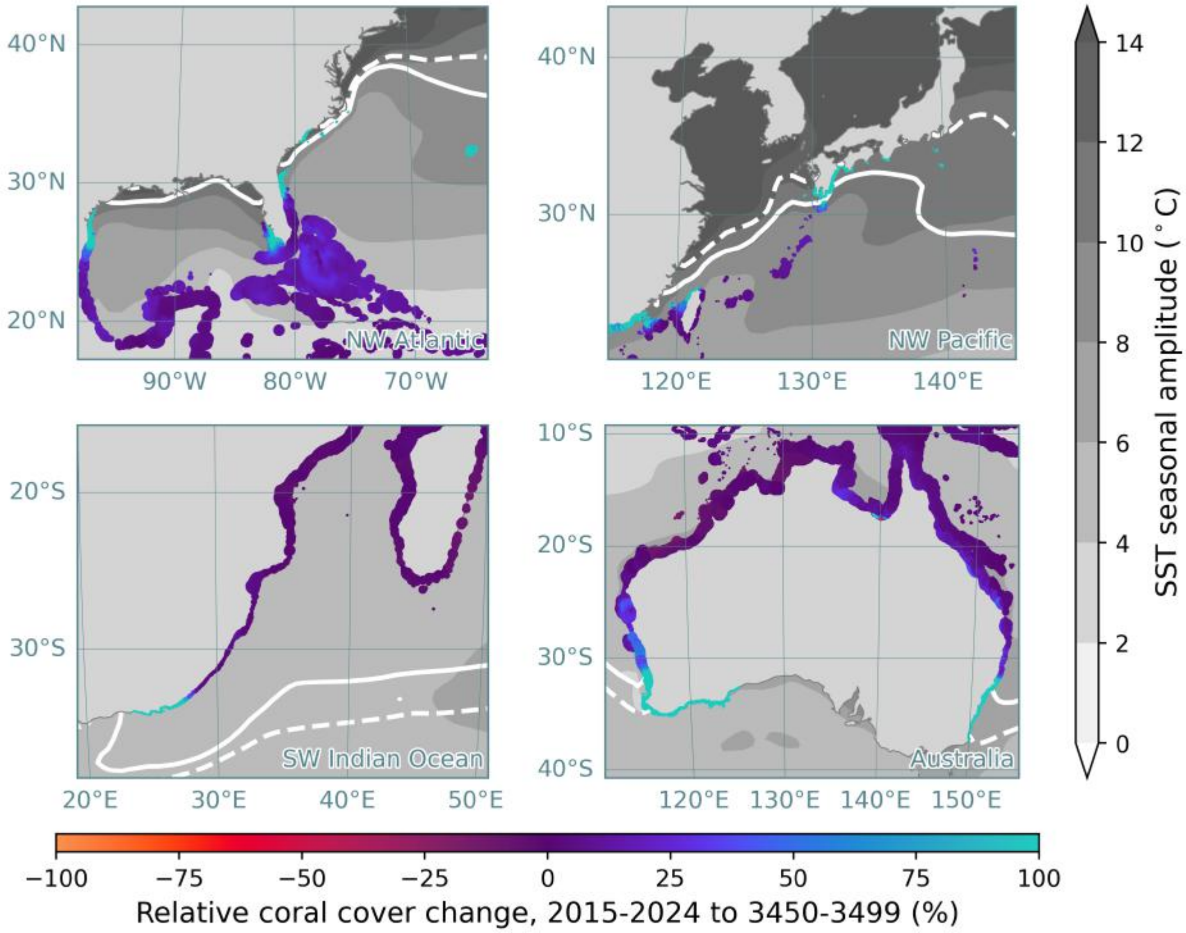


Fig. S19: Mean relative change in tropical coral cover between 2015-2024 and 3450-3499 under SSP2-4.5 *with reduced sensitivity to PAR* ($I_{\mathrm{sat}} = 5$ mol m$^{-2}$ d$^{-1}$) across all ensemble members in CERES for tropical coral assemblages. Sites are scaled by the absolute coral cover between 3450-3499. The ocean is shaded by the amplitude of the seasonal cycle of sea-surface temperature, and the solid and dotted white contours represent the annual minimum 18°C isotherm for the pre-industrial and post-2090 periods respectively. For clarity, only sites with a final coral cover exceeding 0.001 km$^2$ are plotted.

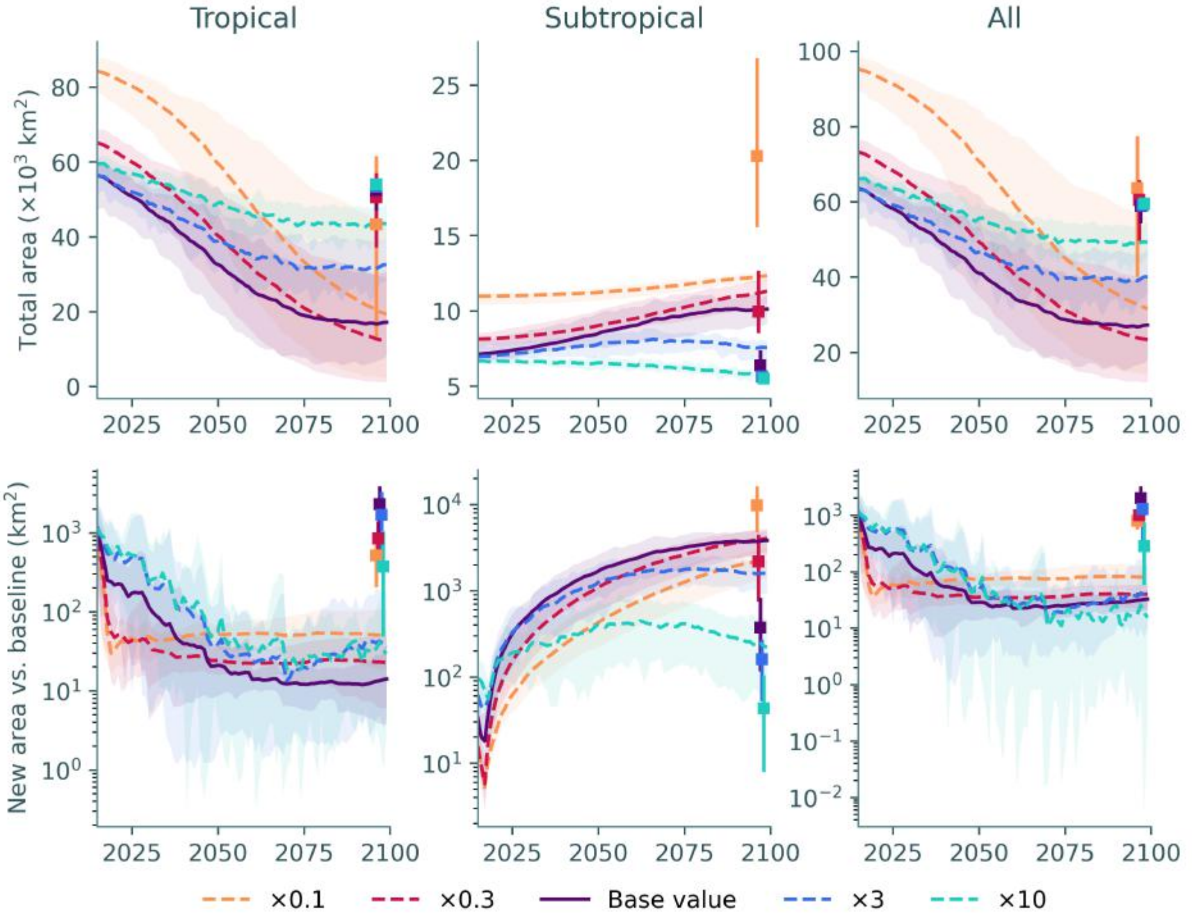


Fig. S20: Total modelled area (*top*) and new coral cover relative to the 2015-2019 baseline (*bottom*) of coral assemblages under SSP2-4.5, under different values of the (maximum) colony linear extension rate ($s_0$). Under low environmental stress, there is a higher equilibrium population size under lower growth rate, possibly due to the lower intensity of selection resulting in a more stable thermal optimum. Squares at the right of each axis represent the mean (long-term) value between 3450-3499, with vertical lines representing the range across the ensemble.

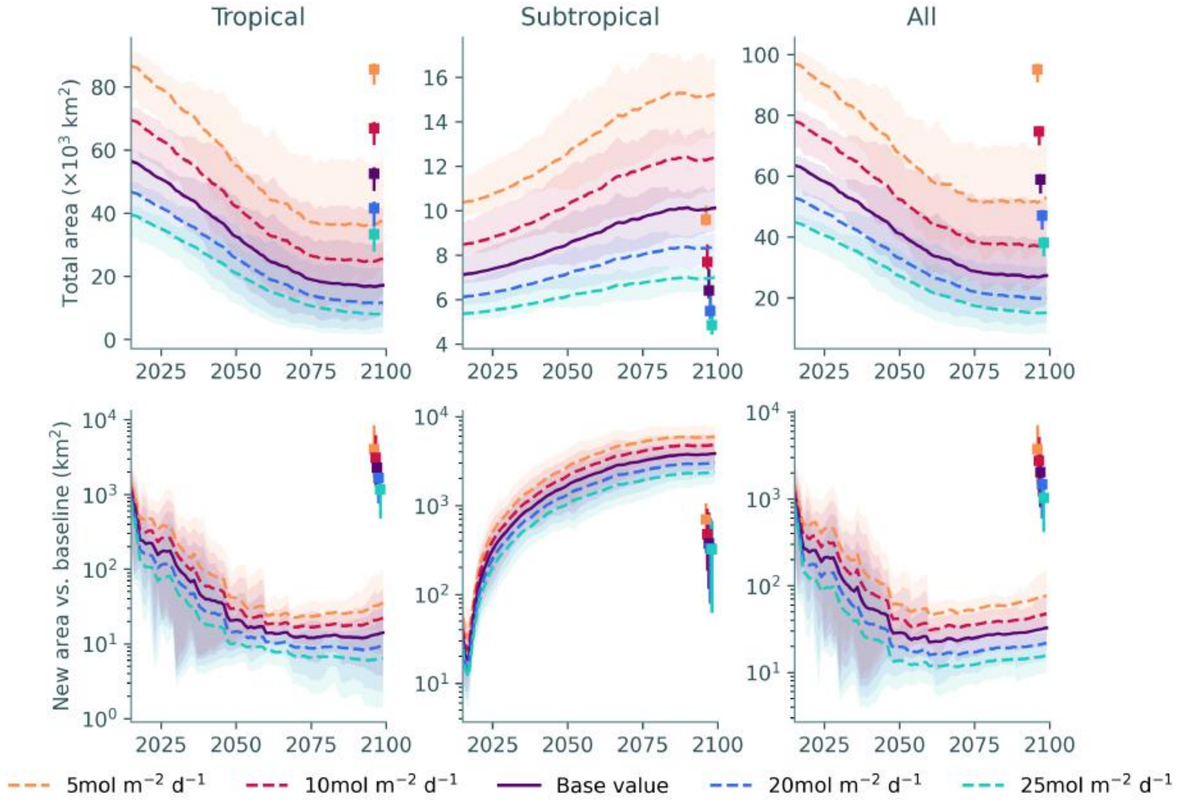


Fig. S21: Total modelled area (*top*) and new coral cover relative to the 2015-2019 baseline (*bottom*) of coral assemblages under SSP2-4.5, under different values of the saturation benthic PAR ($I_{\text{sat}}$). Lower values represent a lower sensitivity to light intensity. Squares at the right of each axis represent the mean (long-term) value between 3450-3499, with vertical lines representing the range across the ensemble.

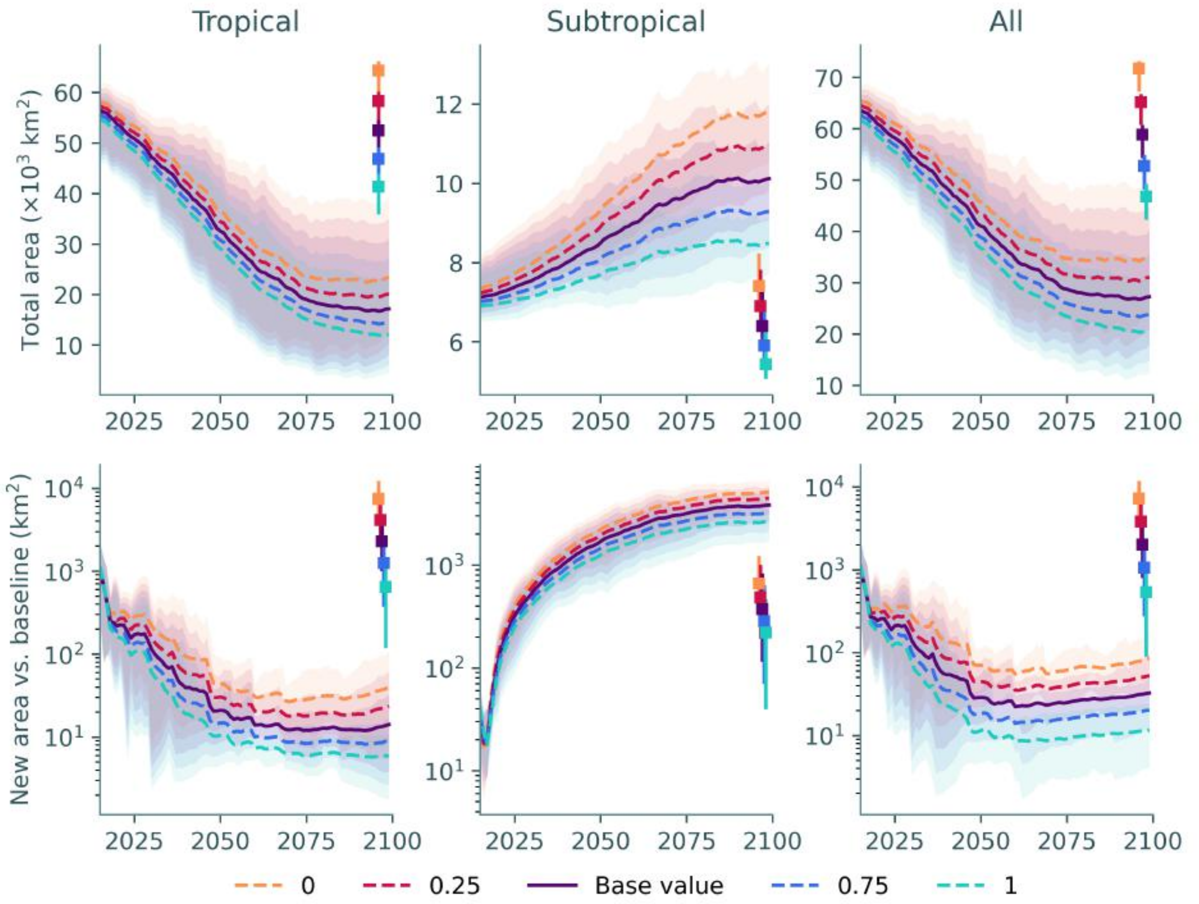


Fig. S22: Total modelled area (*top*) and new coral cover relative to the 2015-2019 baseline (*bottom*) of coral assemblages under SSP2-4.5, under different values of the sensitivity to pH ($c_{\mathrm{pH}}$). Lower values represent a lower sensitivity to pH intensity. Squares at the right of each axis represent the mean (long-term) value between 3450-3499, with vertical lines representing the range across the ensemble.

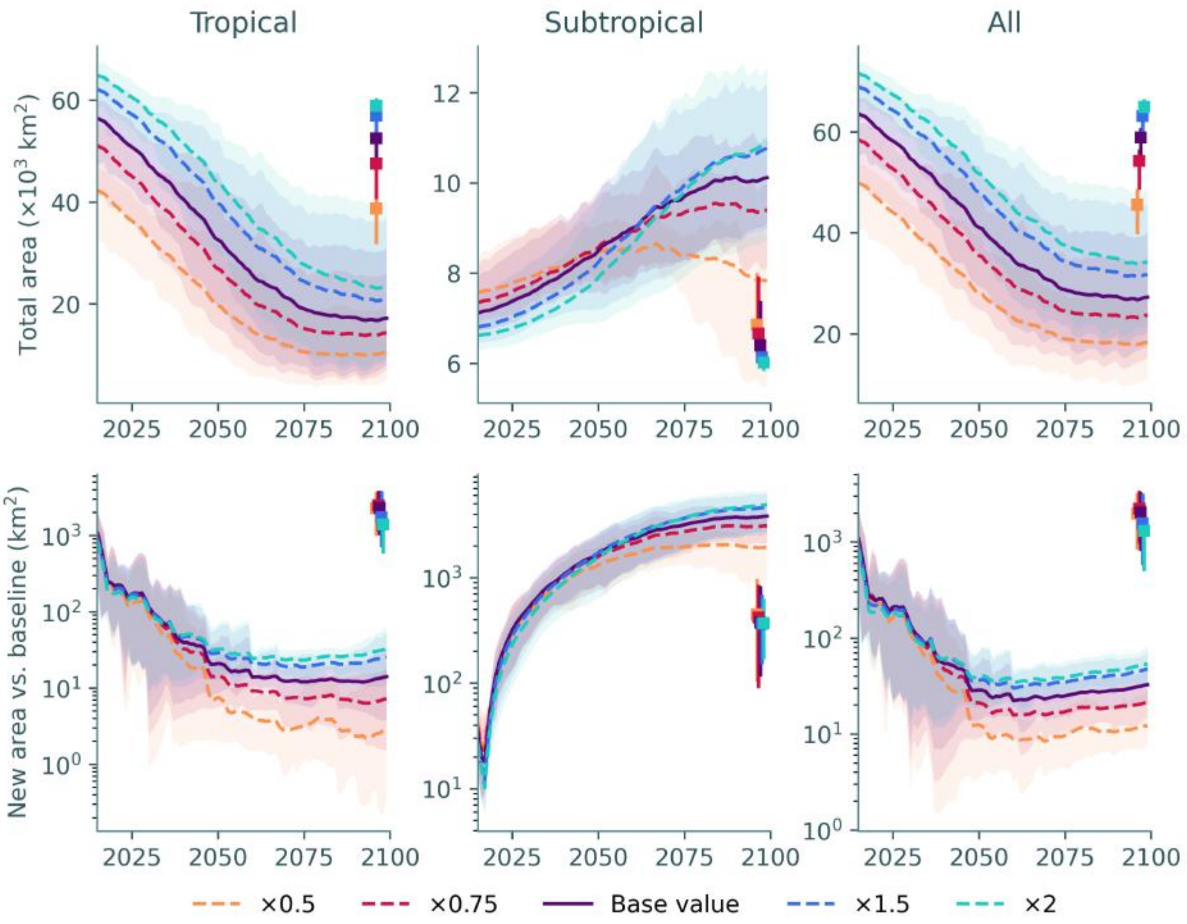


Fig. S23: Total modelled area (*top*) and new coral cover relative to the 2015-2019 baseline (*bottom*) of coral assemblages under SSP2-4.5, under different values of the thermal tolerance ($w$). Lower values represent a higher sensitivity of the linear extension rate to temperature. Squares at the right of each axis represent the mean (long-term) value between 3450-3499, with vertical lines representing the range across the ensemble.

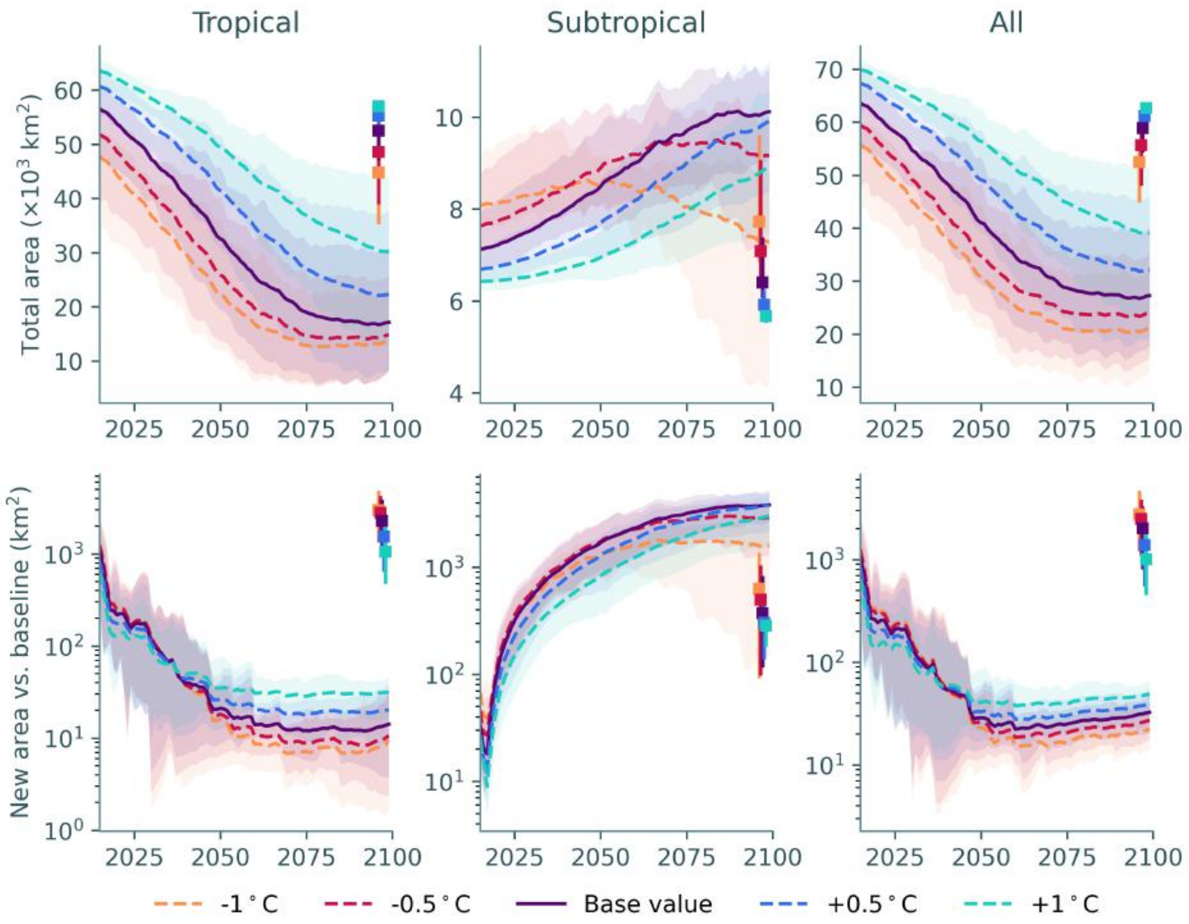


Fig. S24: Total modelled area (*top*) and new coral cover relative to the 2015-2019 baseline (*bottom*) of coral assemblages under SSP2-4.5, under different values of the heat stress threshold ($z_h$). Lower values represent a lower heat stress threshold. Squares at the right of each axis represent the mean (long-term) value between 3450-3499, with vertical lines representing the range across the ensemble.

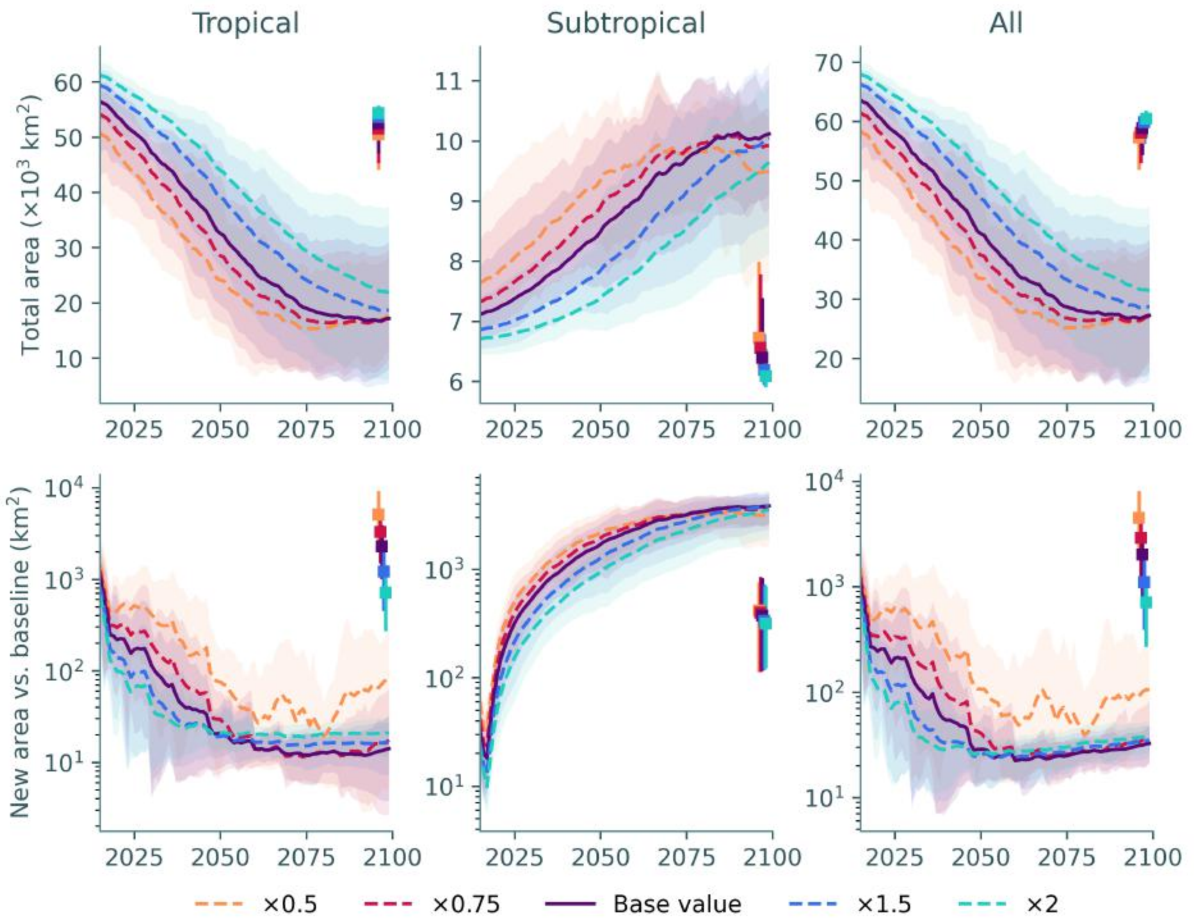


Fig. S25: Total modelled area (*top*) and new coral cover relative to the 2015-2019 baseline (*bottom*) of coral assemblages under SSP2-4.5, under different values of the heat stress tolerance ($w_h$). Lower values represent a higher sensitivity of the mortality rate to heat stress. Squares at the right of each axis represent the mean (long-term) value between 3450-3499, with vertical lines representing the range across the ensemble.

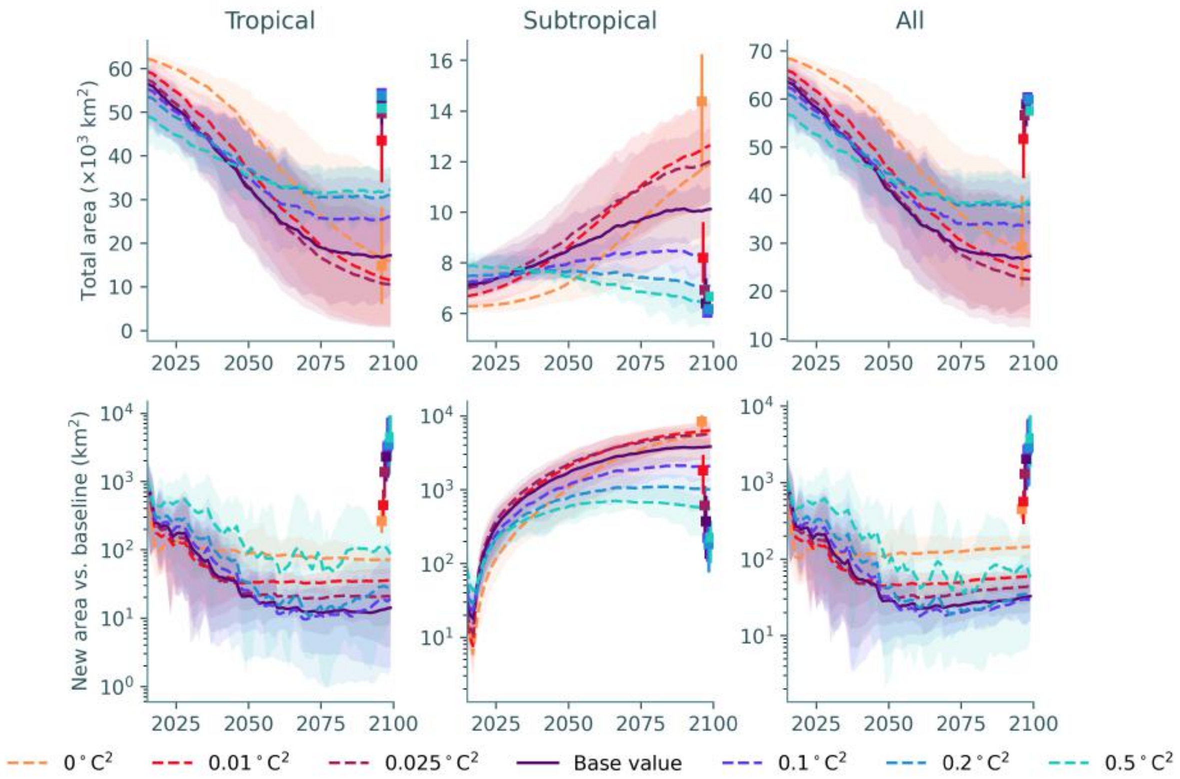


Fig. S26: Total modelled area (*top*) and new coral cover relative to the 2015-2019 baseline (*bottom*) of coral assemblages under SSP2-4.5, under different values of the additive genetic variance ($V$). Squares at the right of each axis represent the mean (long-term) value between 3450-3499, with vertical lines representing the range across the ensemble.

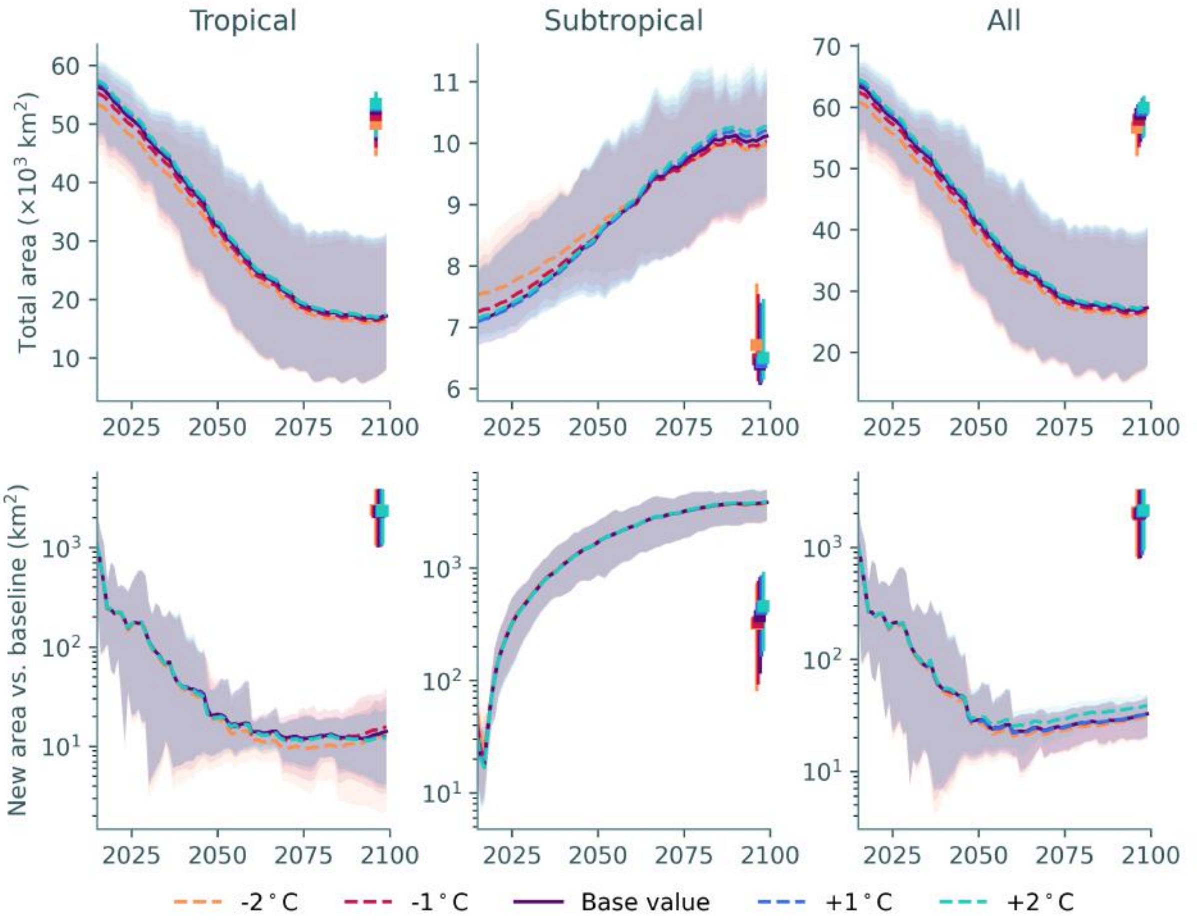


Fig. S27: Total modelled area (*top*) and new coral cover relative to the 2015-2019 baseline (*bottom*) of coral assemblages under SSP2-4.5, under different values of the cold stress threshold ($z_c$). Lower values represent a higher cold stress threshold. Squares at the right of each axis represent the mean (long-term) value between 3450-3499, with vertical lines representing the range across the ensemble.

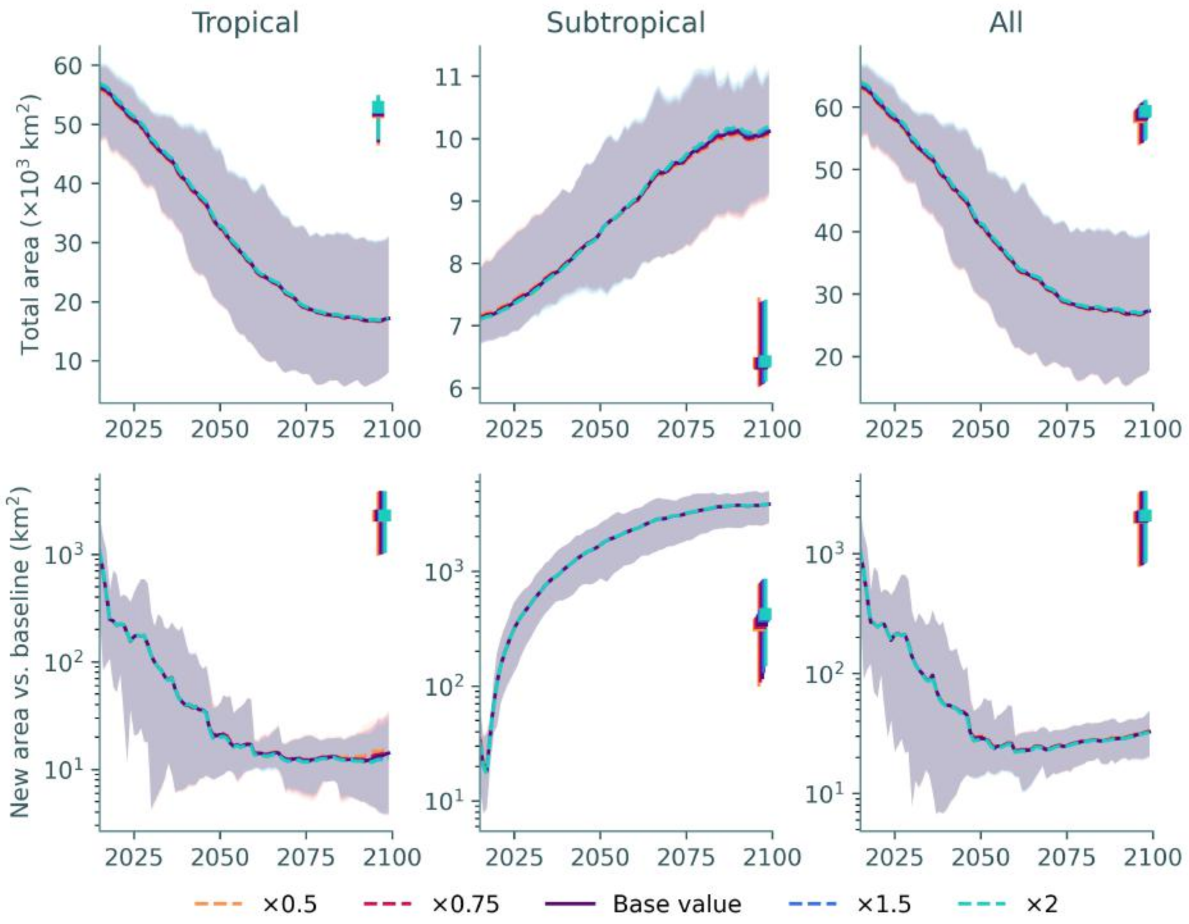


Fig. S28: Total modelled area (*top*) and new coral cover relative to the 2015-2019 baseline (*bottom*) of coral assemblages under SSP2-4.5, under different values of the cold stress tolerance ($w_c$). Lower values represent a higher sensitivity of the mortality rate to cold stress. Squares at the right of each axis represent the mean (long-term) value between 3450-3499, with vertical lines representing the range across the ensemble.

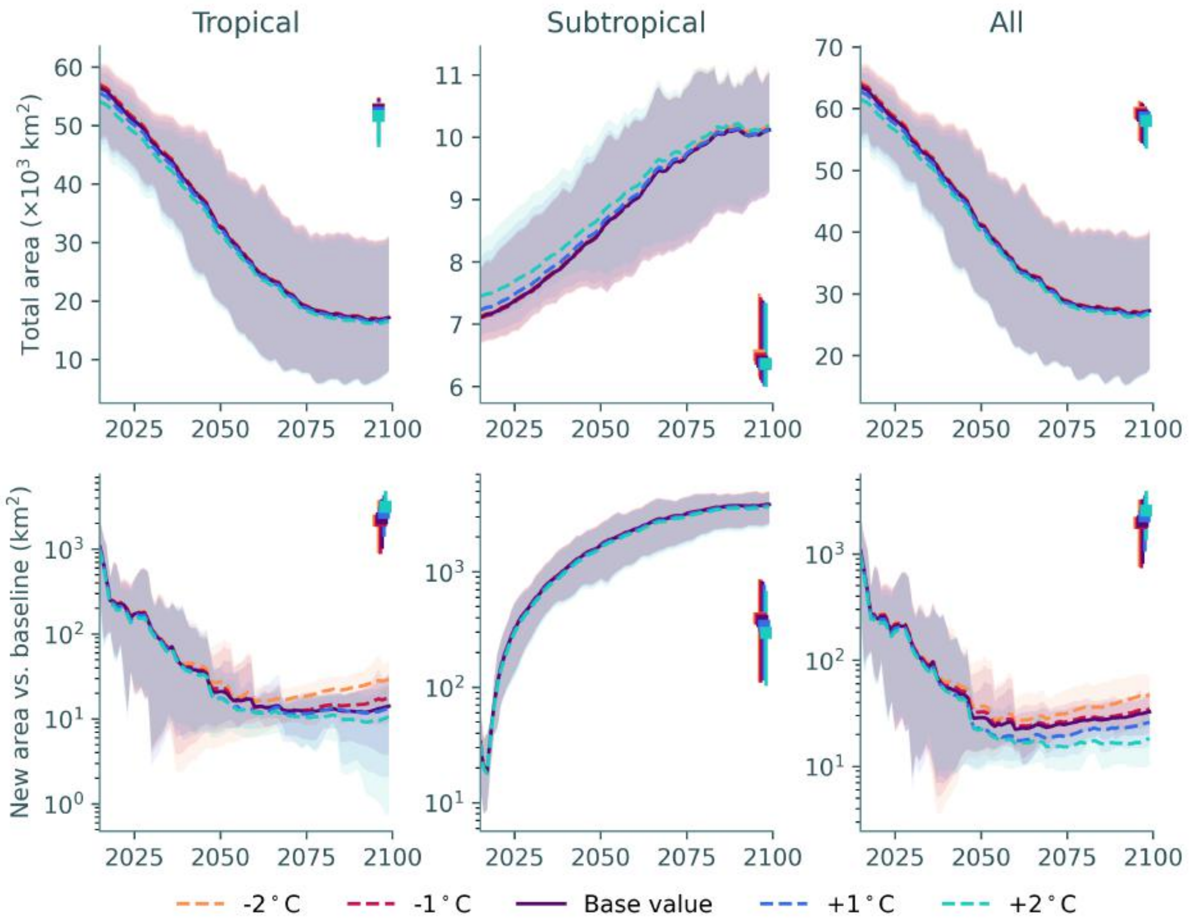


Fig. S29: Total modelled area (*top*) and new coral cover relative to the 2015-2019 baseline (*bottom*) of coral assemblages under SSP2-4.5, under different values of the absolute cold stress threshold ($z_a$). Lower values represent a lower absolute cold stress threshold. Squares at the right of each axis represent the mean (long-term) value between 3450-3499, with vertical lines representing the range across the ensemble.

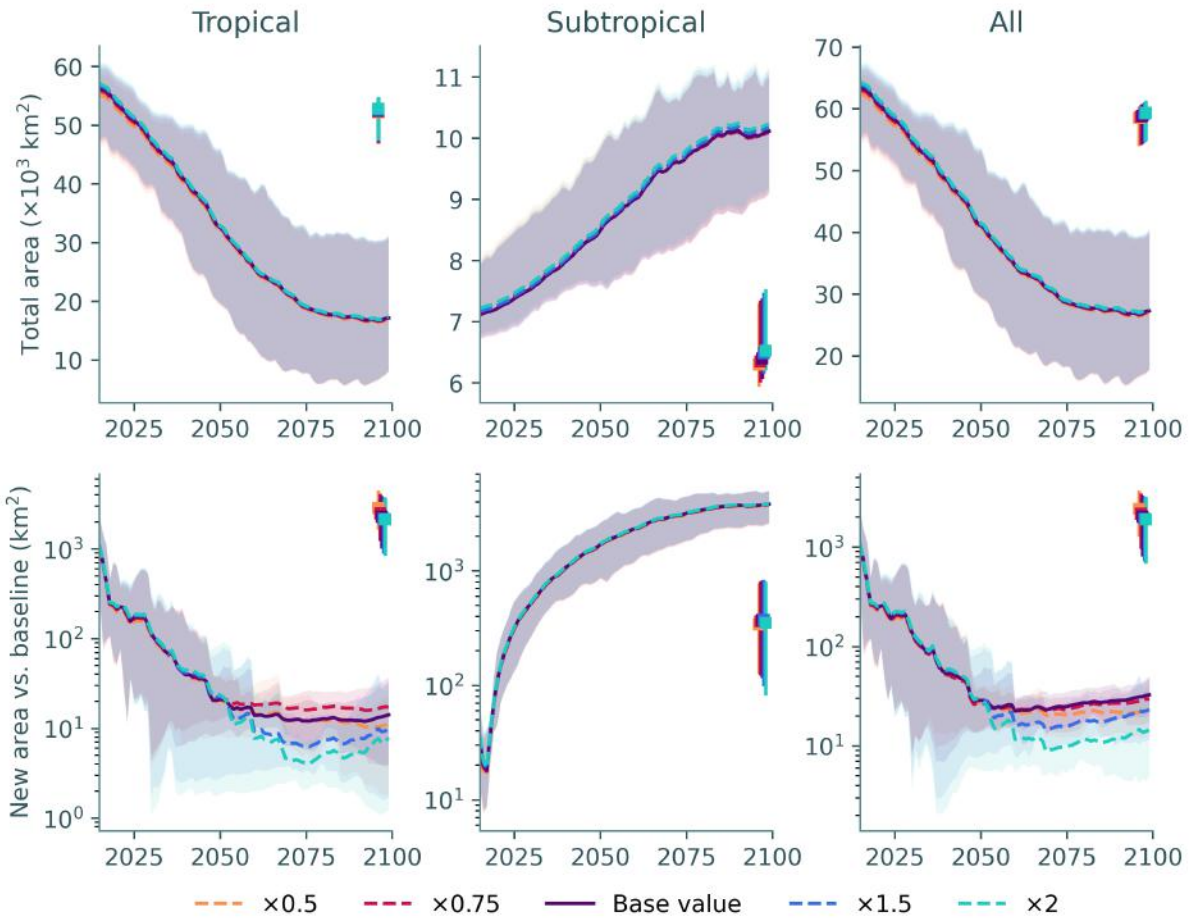


Fig. S30: Total modelled area (*top*) and new coral cover relative to the 2015-2019 baseline (*bottom*) of coral assemblages under SSP2-4.5, under different values of the absolute cold stress tolerance ($w_a$). Lower values represent a higher sensitivity of the mortality rate to absolute cold stress. Squares at the right of each axis represent the mean (long-term) value between 3450-3499, with vertical lines representing the range across the ensemble.

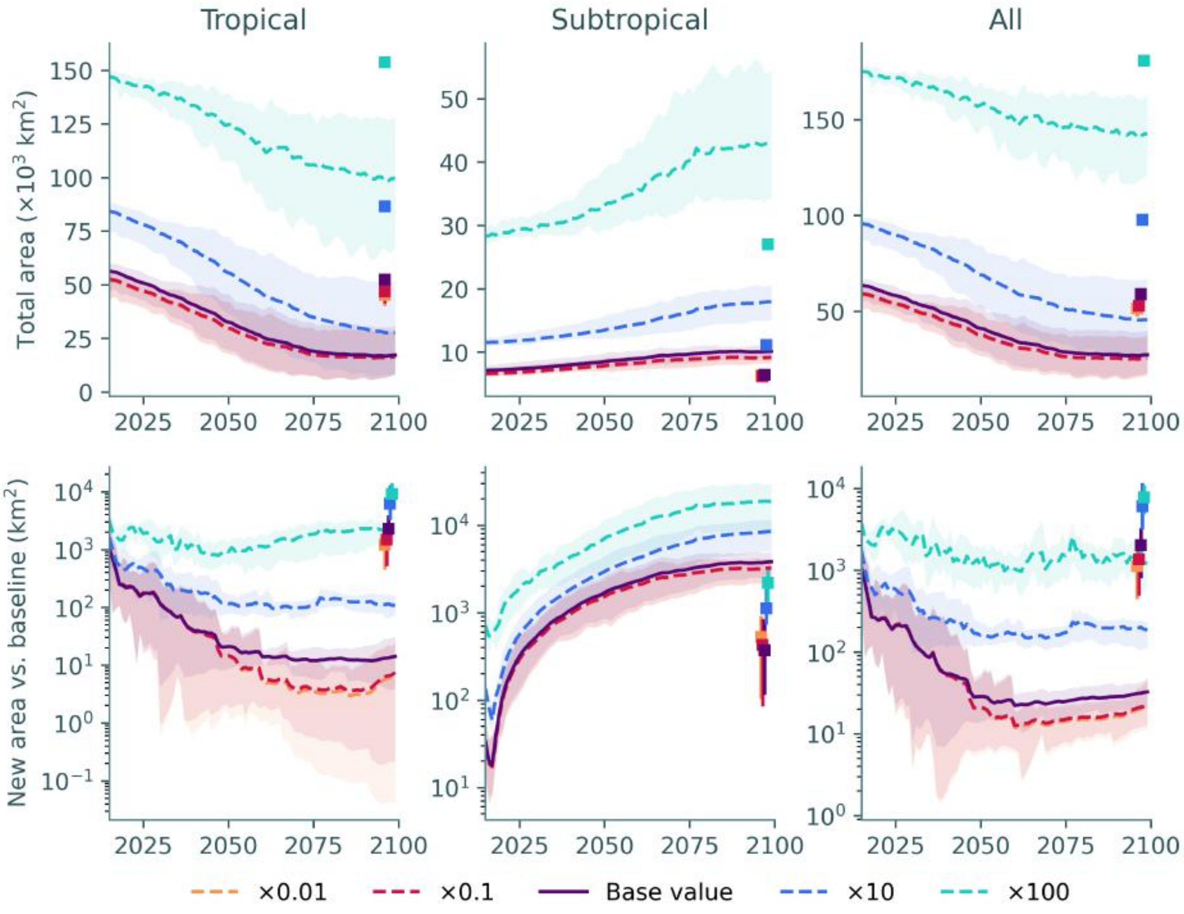


Fig. S31: Total modelled area (*top*) and new coral cover relative to the 2015-2019 baseline (*bottom*) of coral assemblages under SSP2-4.5, under different values of effective fecundity ($f$). Squares at the right of each axis represent the mean (long-term) value between 3450-3499, with vertical lines representing the range across the ensemble.

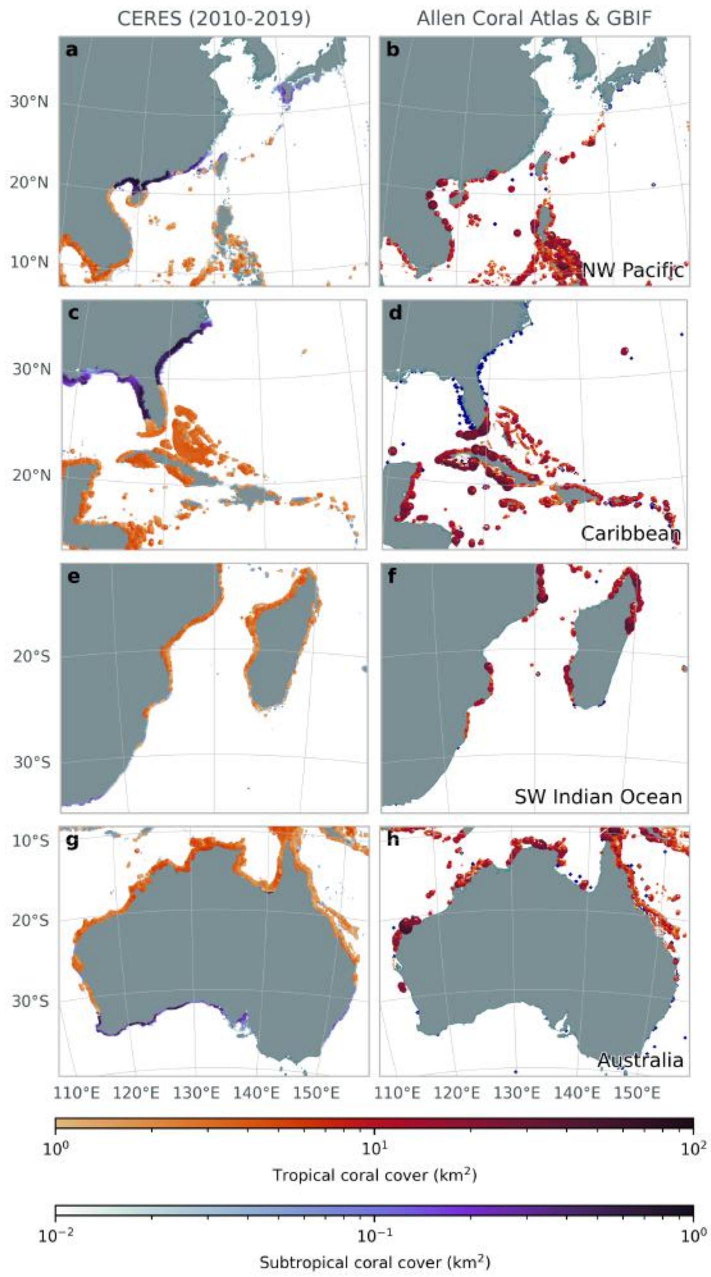


Fig. S32: Mean coral cover from 2010-2019 across all ensemble members in CERES with effective fecundity ($f$) increased by a factor of 10 (*left*); and coral cover from the Allen Coral Atlas (*28*) and (as blue points) scleractinian coral occurrence records within 20 m depth (*30*) (*right*). To show both reef and non-reef coral cover data from CERES, we only plot reef coral cover where it exceeds 1 km$^2$.

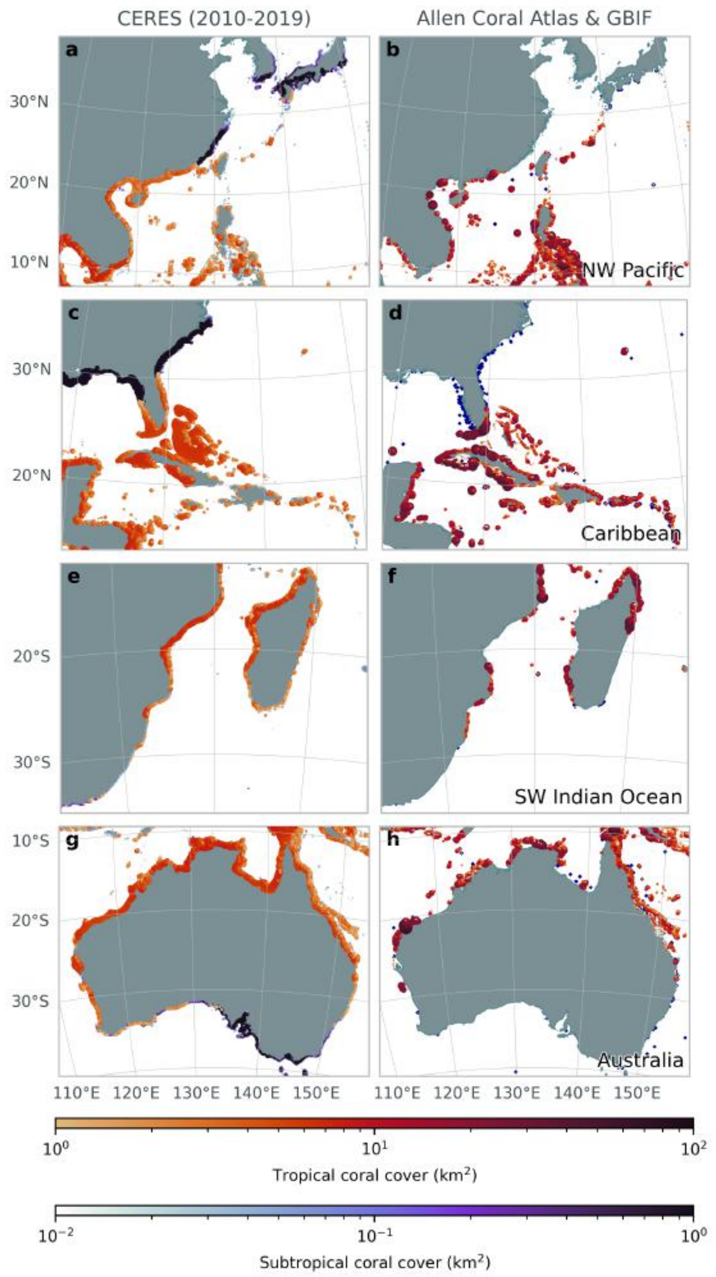


Fig. S33: Mean coral cover from 2010-2019 across all ensemble members in CERES with effective fecundity ($f$) increased by a factor of 100 (*left*); and coral cover from the Allen Coral Atlas (*28*) and (as blue points) scleractinian coral occurrence records within 20 m depth (*30*) (*right*). To show both reef and non-reef coral cover data from CERES, we only plot reef coral cover where it exceeds 1 km$^2$.

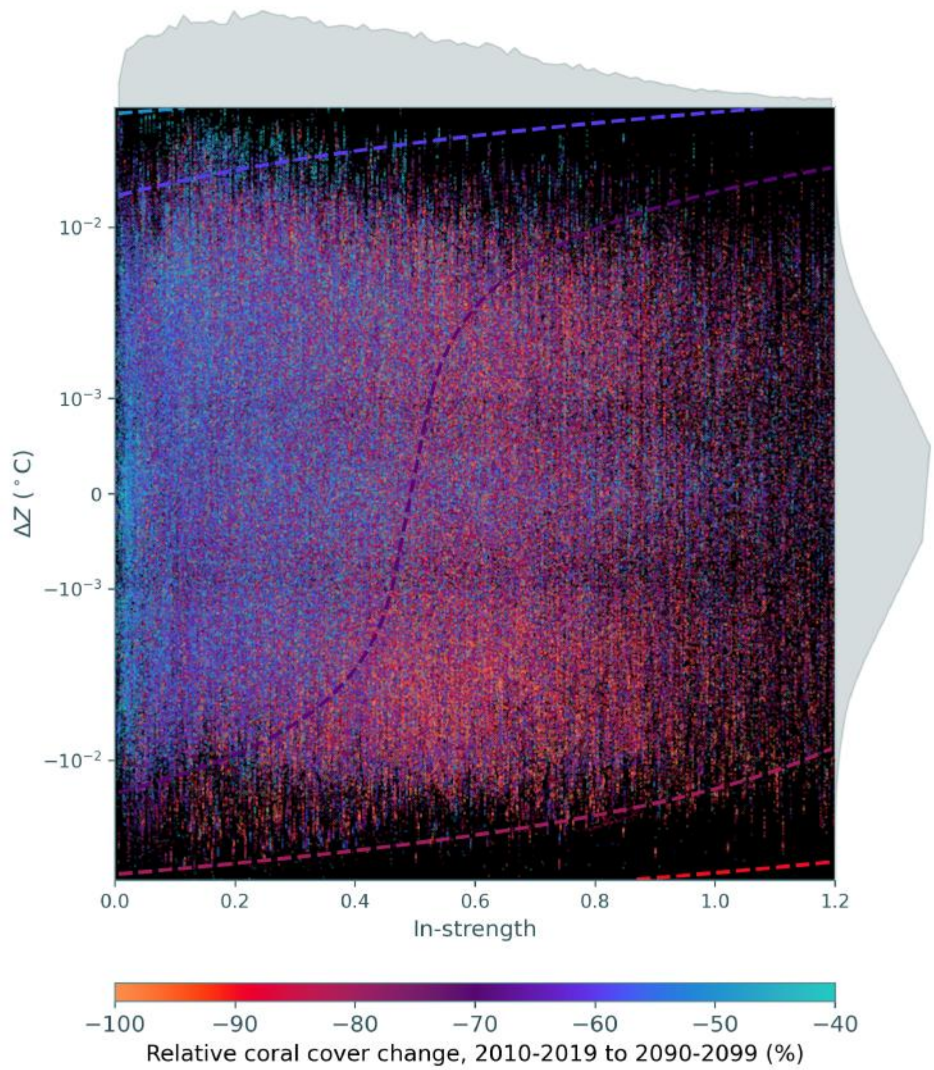


Fig. S34: Predicted (background) and simulated (points) relative change in tropical coral cover over the 21$^{\text{st}}$ century, as a function of the change in ratio of incoming to outgoing coral larvae (in-strength), and the difference between the average thermal optimum trait value of incoming larvae and existing corals at the site ($\Delta Z$). Only sites where the tropical coral cover originally exceeded 10% are included. The predicted change in coral cover was calculated from a linear mixed model, with random intercepts varying by ensemble member. Simulated data are plotted with random effects removed. Histograms show the distribution of predictor variables across reef sites. Note that the $y$ axis is plotted on a symlog scale due to the distribution of $\Delta Z$.

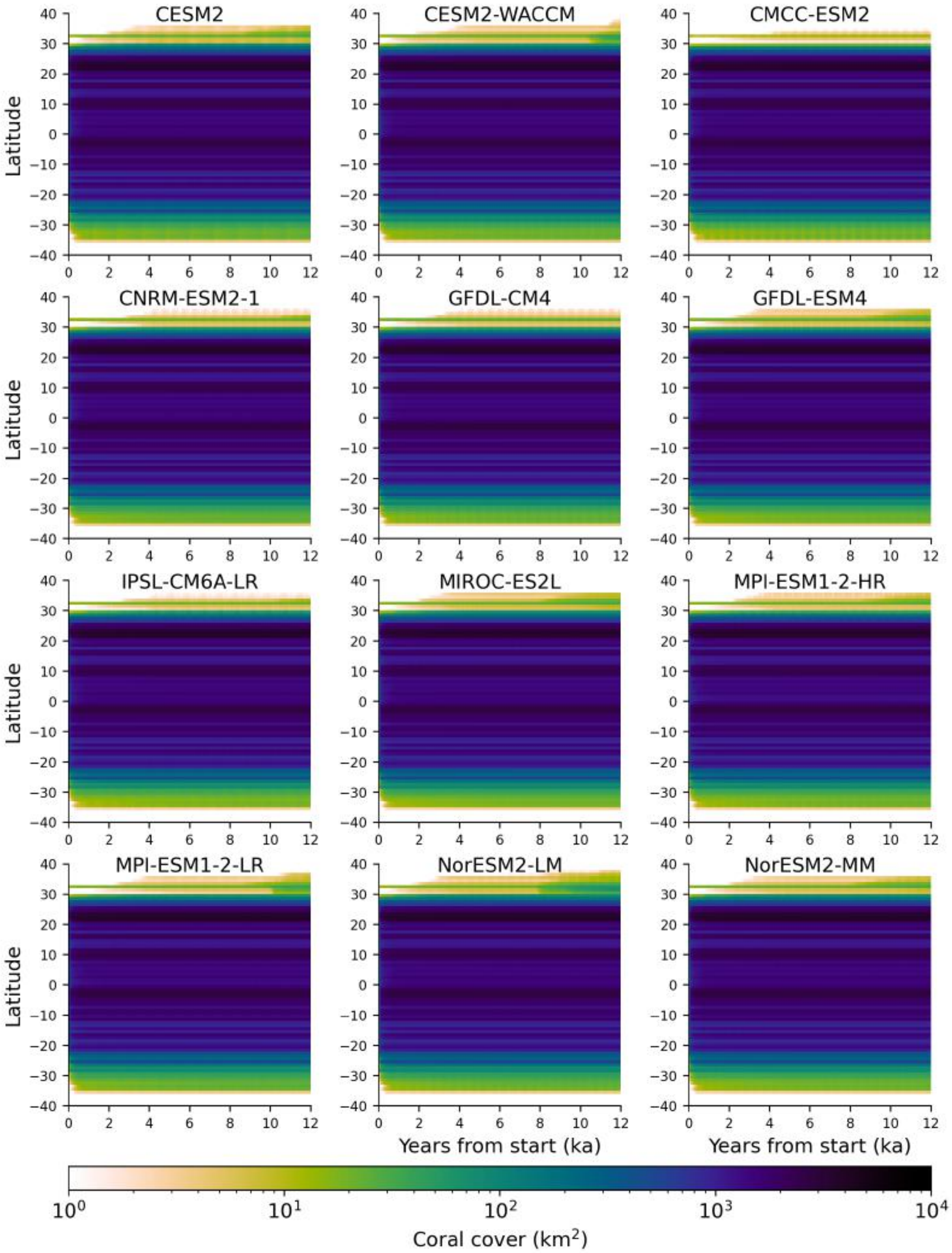


Fig. S35: Total coral cover per latitude band under looping *piControl* forcing, after initialisation from the Allen Coral Atlas (*28*), for the twelve CMIP6 models used.

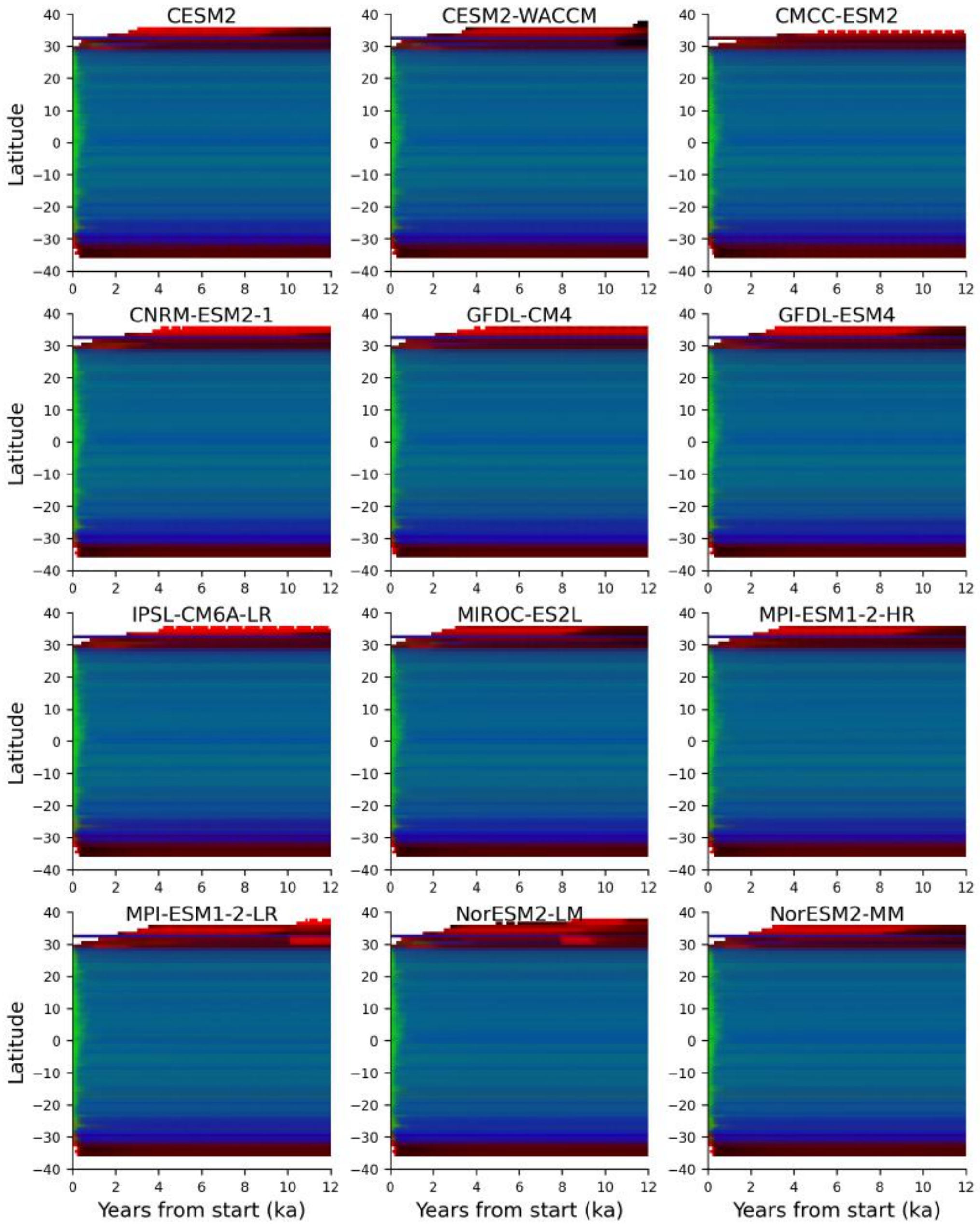


Fig. S36: Coral community composition within each latitude band under looping *piControl* forcing. There are three degrees of freedom in proportional community composition normalised to total coral cover (four coral groups, with proportions summing to 1). Proportions belonging to **Fast-ST**, **Fast-T**, and **Slow-T** are respectively mapped to red, blue and green channels respectively, with **Slow-ST** effectively mapped to lightness. Community composition is only shown where coral cover exceeds 1 $km^2$ for a latitude band.

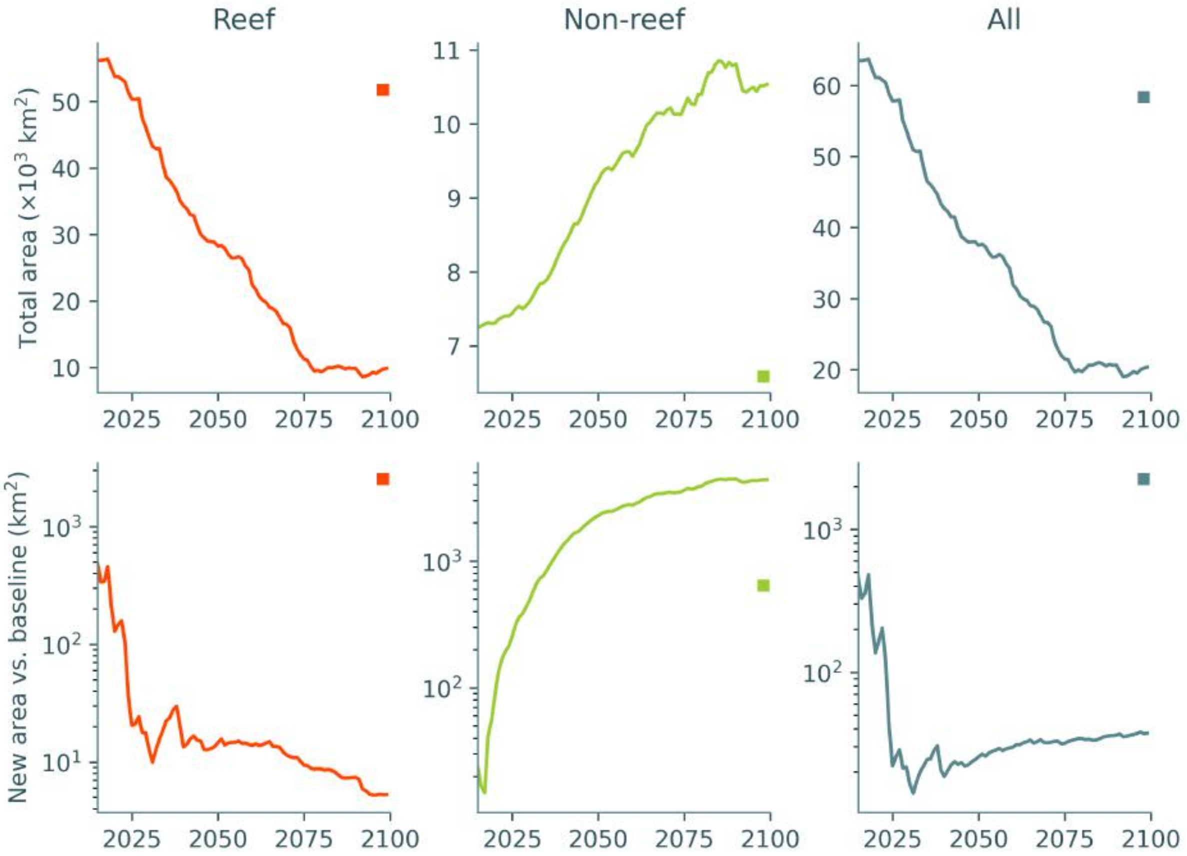


Fig. S37: Total modelled area (*top*) and new coral cover relative to the 2015-2019 baseline (*bottom*) of coral assemblages under SSP2-4.5, with the shaded area (usually not visible due to the low variability) showing the range across different random combinations of temporal connectivity matrices. Squares at the right of each axis represent the mean (long-term) value between 3450-3499, with vertical lines (not visible) representing the range across the ensemble.

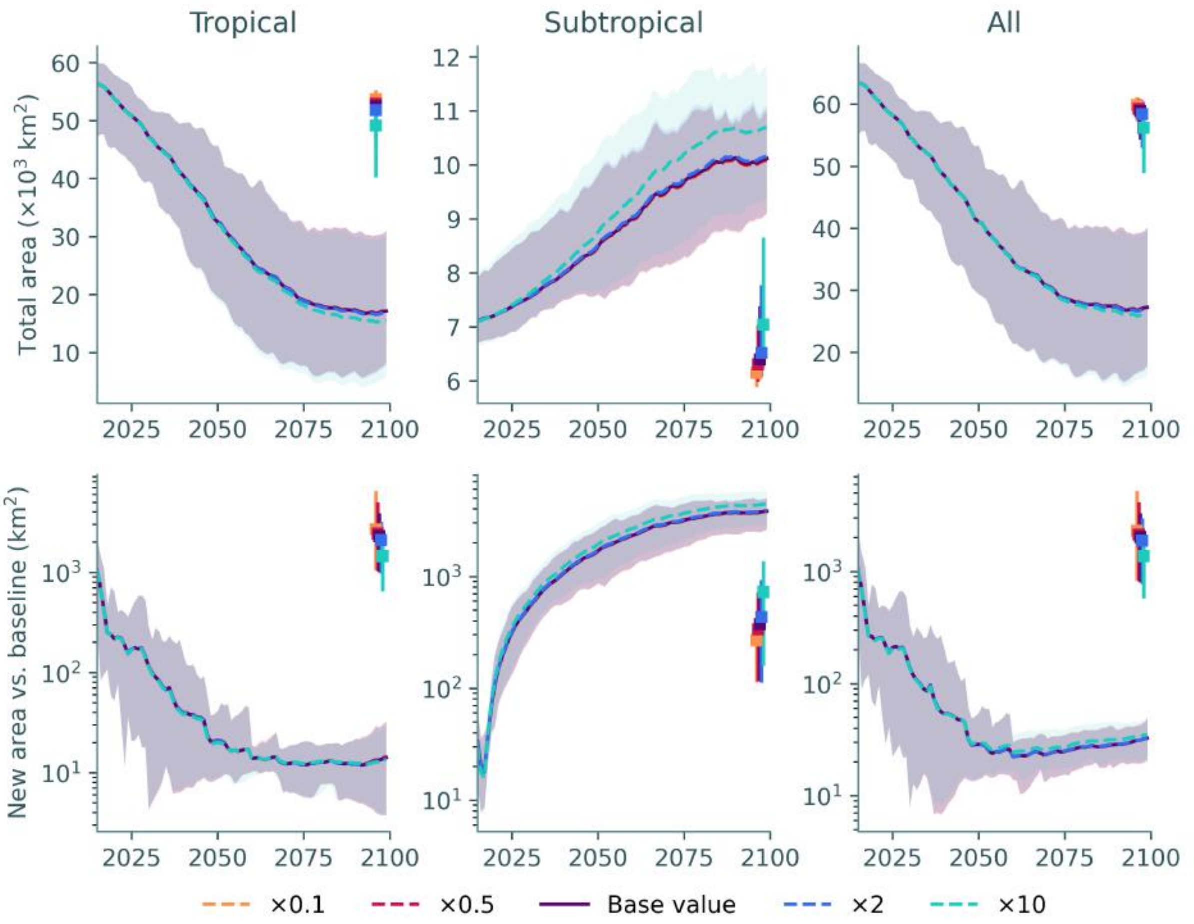


Fig. S38: Total modelled area (*top*) and new coral cover relative to the 2015-2019 baseline (*bottom*) of coral assemblages under SSP2-4.5, under different values of the selection throttling threshold ($C_{\mathrm{throttle}}$). Squares at the right of each axis represent the mean (long-term) value between 3450-3499, with vertical lines representing the range across the ensemble.

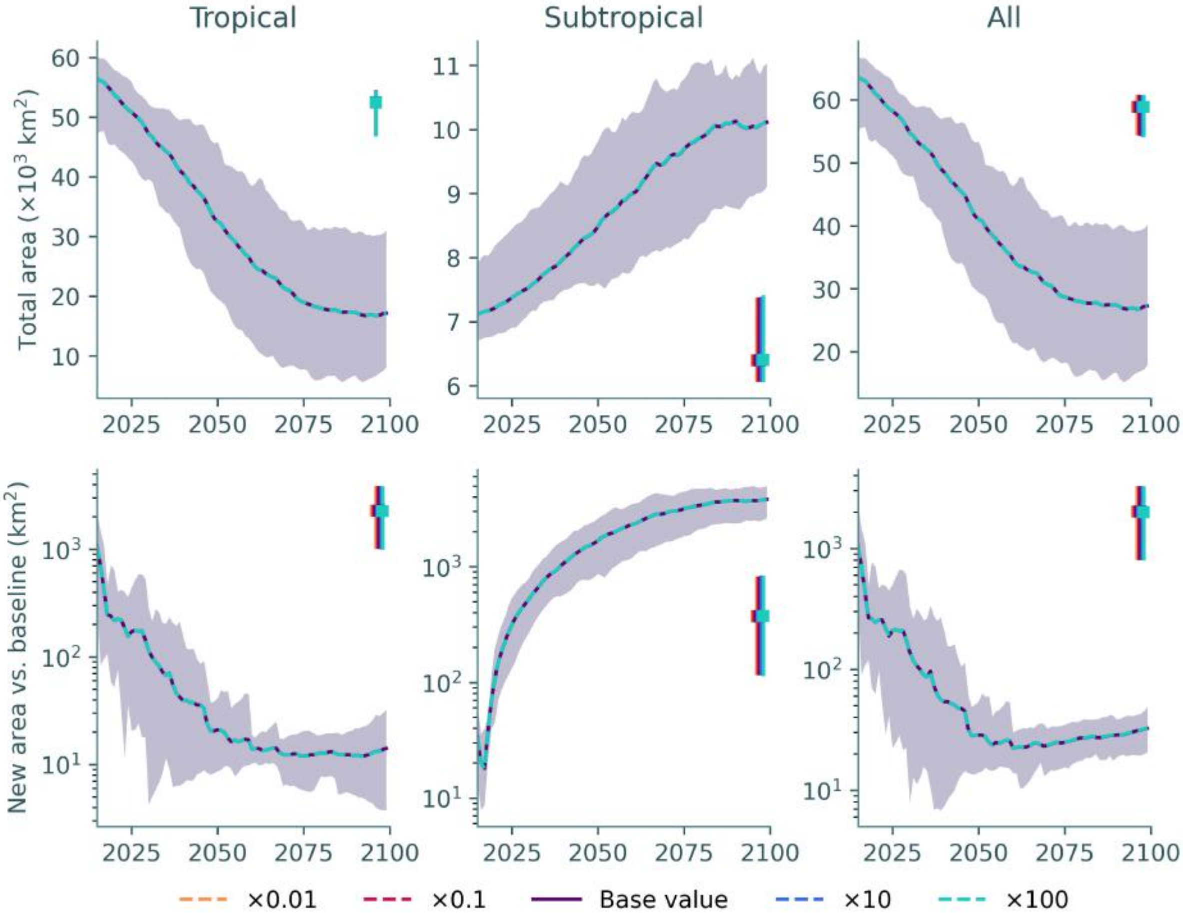


Fig. S39: Total modelled area (*top*) and new coral cover relative to the 2015-2019 baseline (*bottom*) of coral assemblages under SSP2-4.5, under different values of the minimum nonzero coral cover ($C_{\mathrm{min}}$). Squares at the right of each axis represent the mean (long-term) value between 3450-3499, with vertical lines representing the range across the ensemble.
